# Maintenance of adaptive dynamics and no detectable load in a range-edge out-crossing plant population

**DOI:** 10.1101/709873

**Authors:** Margarita Takou, Tuomas Hämälä, Evan M. Koch, Kim A. Steige, Hannes Dittberner, Levi Yant, Mathieu Genete, Shamil Sunyaev, Vincent Castric, Xavier Vekemans, Outi Savo-lainen, Juliette de Meaux

**Affiliations:** Institute of Botany, University of Cologne, Cologne, Germany; Department of Plant and Microbial Biology, University of Minnesota, St. Paul, Minnesota, USA; Department of Biomedical Informatics, Harvard Medical School, Boston, Massachusetts, USA; School of Life Sciences, University of Nottingham, Nottingham NG7 2RD, UK; Univ. Lille, CNRS, UMR 8198 - Evo-Eco-Paleo, F-59000 Lille, France; Department of Ecology and Genetics, University of Oulu, FIN-90014 Oulu, Finland

**Author notes:** **Corresponding author:** Juliette de Meaux.

**Keywords:** range expansion, adaptation, deleterious mutations, self-incompatibility locus, negative frequency-dependent selection, selective sweeps

## Abstract

During range expansion, edge populations are expected to face increased genetic drift, which in turn can alter and potentially compromise adaptive dynamics, preventing the removal of deleterious mutations and slowing down adaptation. Here, we contrast populations of the European sub-species *Arabidopsis lyrata* ssp *petraea*, which expanded its Northern range after the last glaciation. We document a sharp decline in effective population size in the range-edge population and observe that non-synonymous variants segregate at higher frequencies. We detect a 4.9% excess of derived non-synonymous variants per individual in the range-edge population, suggesting an increase of the genomic burden of deleterious mutations. Inference of the fitness effects of mutations and modeling of allele frequencies under the explicit demographic history of each population predicts a depletion of rare deleterious variants in the range-edge population, but an enrichment for fixed ones, consistent with the bottleneck effect. However, the demographic history of the range-edge population predicts a small net decrease in per-individual fitness. Consistent with this prediction, the range-edge population is not impaired in its growth and survival measured in a common garden experiment. We further observe that the allelic diversity at the self-incompatibility locus, which ensures strict outcrossing and evolves under negative frequency-dependent selection, has remained unchanged. Genomic footprints indicative of selective sweeps are broader in the Northern population but not less frequent. We conclude that the outcrossing species *A. lyrata* ssp *petraea* shows a strong resilience to the effect of range expansion.

## Introduction

Range expansion events, like the postglacial colonization of Northern Europe and Scandinavia from Southern refugia, have had wide influence on the distribution of genetic diversity within species (Hewitt 2000). Through its impact on multiple population genetic processes, range expansion has cascading effects on adaptive dynamics (Excoffier et al. 2009). Indeed, it increases drift (Hallatschek et al. 2007), and leads to both a progressive loss of genetic diversity and increased levels of population differentiation along the expansion route (Austerlitz et al. 1997; Corre and Kremer 1998; Muller et al. 2008; Excoffier et al. 2009; Slatkin and Excoffier 2012). As a consequence, fitness is expected to decrease at the front of the expanding range, causing what is known as the expansion load. The majority of those mutations remain at low frequencies or are lost, but some quickly fix, a phenomenon sometimes termed allele surfing (Klopfstein et al. 2006; Peischl et al. 2013). Although non-synonymous and potentially deleterious mutations are more likely to fix in bottlenecked populations, where the removal of new deleterious mutations is less efficient, it takes some evolutionary time until a significant load accumulates (Lohmueller 2014; Simons et al. 2014; Balick et al. 2015; Do et al. 2015).

Expansion load can interfere with adaptive dynamics. Locally adapted populations that move out of their core range are expected to evolve towards new adaptive peaks (Colautti and Barrett 2013; Savolainen et al. 2013; Wos and Willi 2018). In a population carrying an expansion load, larger adaptive steps might be required to establish a novel range edge, resulting in a slowdown of expansion, especially when dispersal is limiting (Henry et al. 2015). Theoretical studies report complex interactions among parameters such as the strength and heterogeneity of selection, the rate of expansion, as well as the architecture of traits under selection. Expansion rate and adaptive requirements to the newly colonized environments can jointly modulate the fitness decrease observed at the range edge (Gilbert et al. 2017; Gilbert et al. 2018). However, to the best of our knowledge, these predictions remain practically untested in natural populations.

The speed of range expansion can also be limited by species interactions, if these are necessary for reproductive success and survival (Louthan et al. 2015). Many flowering plants rely on insects for pollination and thus fertility (Gibbs 2014). As species expand their range, efficient pollinators can become rare, and a shift towards selfing may help restore reproductive assurance and avoid Allee effects (Jain 1976; Morgan et al. 2005; Gascoigne et al. 2009). Transitions to selfing or mixed-mating systems have often been associated with range expansion (Goodwillie et al. 2005; Levin 2010; Laenen et al. 2018; Baker 1995 but see Cheptou 2012). However, mating system shifts can compromise adaptive processes by exposing populations to inbreeding depression and loss of genetic diversity as they face stress at the margin of their ecological niche (Baker 1995; Slatkin 1995; Ingvarsson 2002; Barrett 2003; Glémin and Ronfort 2013). Yet, increases in the selfing rate can also contribute to the purging of deleterious mutations (Pujol et al. 2009; Glémin and Ronfort 2013; Hadfield et al. 2017; Roessler et al. 2019) and promote the emergence of high fitness individuals at the front range of expansion (Klopfstein et al. 2006). In fact, selfing species generally show the greatest overall range size (Grossenbacher et al. 2015). In this context, plant species that have maintained a strictly outcrossing mating system across their expanded distribution range are particularly intriguing.

The European sub-species *Arabidopsis lyrata ssp. lyrata* has expanded its range Northwards after the last glaciation (Clauss and Koch 2006; Schierup et al. 2006; Koch 2019). Its patchy populations are found from Central Europe to the North of Scandinavia (Hoffmann 2005). Northern populations in *A. l. ssp. petraea* show a strong reduction in diversity (Wright et al. 2003; Muller et al. 2008; Ross-Ibarra et al. 2008; Pyhäjärvi et al. 2012; Mattila et al. 2017). Yet, there is evidence that *A. l. ssp. petraea* populations at the Northern range edge are locally adapted. Reciprocal transplant studies between the Northern and Central European populations showed that Northern populations have the highest survival rate in their location of origin consistent with signals of local adaptation (Leinonen et al. 2009). Major developmental traits such as flowering time, as well as the response to abiotic stress factors, seem to have been targets of natural selection (Sandring et al. 2007; Toivainen et al. 2014; Mattila et al. 2016; Davey et al. 2018; Hämälä and Savolainen 2018). Reciprocal transplant experiments across four sites in Europe, as well as between populations of different altitude in Norway, indicated that populations at the range margins were locally adapted (Hämälä and Savolainen 2018).

*A. lyrata ssp. lyrata* enforces self-incompatibility (SI) via the multiallelic S-locus specific to the Brassicaceae family (Bateman 1955; Kusaba et al. 2001). Phylogenetic and genomic analyses of the S-locus have shown that strong negative frequency-dependent selection caused early diversification of the S-locus within the family and a high degree of sharing of S-allele lineages across species (Dwyer et al. 1991; Vekemans et al. 2014). The loss of SI, however, evolved repeatedly in the family (Tsuchimatsu et al. 2012; Vekemans et al. 2014; Durvasula et al. 2017). In fact, some populations of the closely related North American subspecies *A. l. ssp. lyrata*, lost obligate outcrossing at their range margin (Mable et al. 2005; Griffin and Willi 2014; Willi et al. 2018). This transition to selfing has been recently associated with a sharp decrease in average population fitness (Willi et al. 2018). In the sub-species *A. l. ssp. petraea*, instead, SI appears to have been maintained, presumably due to the inbreeding depression, which has been demonstrated using forced selfing (Kärkkäinen et al. 1999; Sletvold et al. 2013).

To gain insight into the combined effects of demographic history and selection processes in an out-crossing range-edge population, we quantified the demographic impact of range expansion in a Northern population of the sub-species *A. l. ssp. petraea* and examined its impact on both negative and positive selection. We compare this population to two populations representative of the core of the species range and specifically ask: i) can we document a decreased efficacy of negative selection in the range-edge population and an increase in the individual burden of deleterious mutations?, ii) does the range-edge population show a decrease in S-allele diversity as expected by an ongoing mating system shift?, and iii) do we detect a slowdown of adaptive dynamics in range-edge *A. l. ssp. petraea* populations?

We document a strong bottleneck and increased frequency of non-synonymous variants indicative of progressive range expansion. Population genetics modeling, genomic measures and common garden analysis of plant fitness indicate that the bottleneck was too short and not severe enough to allow the accumulation of a burden with significant effect on observed fitness. We further observe that negative frequency-dependent selection on S-alleles has remained efficient and find no evidence that the response to positive selection is impaired in the range-edge population. The outcrossing subspecies *A. l.* ssp *petraea* shows a strong resilience to the effect of range expansion.

## Results

### Demographic history of three European *A. lyrata ssp. petraea* populations confirms a scenario of range expansion

We analyzed whole genome sequence data for 46 *Arabidopsis lyrata* individuals, of which, 22 were collected in a range edge population in Norway (Spiterstulen, SP), and 17 and 7 individuals from two core populations in Germany (Plech, PL) and Austria (a scattered sample, AUS; Fig. S1a), respectively. A principal component analysis (PCA) confirmed that our sample was partitioned in three geographically and genetically distinct populations. The first principal component (PC) explained 24.95% of the variance, separating the Northern site from the two Central European sites (PL and AUS). The second PC (6.82%) differentiated the AUS and PL sites. AUS individuals were more scattered than SP and PL individuals, presumably because AUS individuals were collected over a broader area (see methods, Fig. S1b). Admixture analysis showed that our samples formed three populations, without any indication of admixture within populations. Our samples were well described with K=2 clusters (cross-validation error, cv = 0.397). The SP individuals formed a unique cluster, while PL-AUS individuals grouped together in one cluster. The second most probable scenario (cv = 0.419) was K=3, with each population forming its own cluster (Fig. S1c). We further calculated *F_ST_* across 10 kb non-overlapping windows along the genome. Mean *F_ST_* was 0.231 (median of 0.232) and 0.234 (median of 0.236) for SP vs. PL or AUS, respectively. Between PL and AUS, differentiation was much lower, with a mean *F_ST_* value of 0.079 (median of 0.047). Thus, most of the genetic differentiation resides between Northern and Central European populations and not between PL and AUS. The average number of nucleotide differences between pairs of individuals from distinct sites (d_xy_) confirmed the pattern of inter-population differentiation (Table S1). Within populations, nucleotide diversity was estimated as the average number of pairwise differences per sites (π) across the same non-overlapping 10 kb windows. Mean nucleotide diversity of the genomic windows was π=0.0081, π=0.0067 and π = 0.0055 for PL, AUS and SP, respectively (Table S1).

PCA, Fst and STRUCTURE provide measures genetic differentiation between individuals and populations. Genetic differentiation, in turn, is a result of the time since divergence, the intensity of gene flow, and the size of the population. Two populations could have split a long time ago, and nevertheless remain genetically similar if their population size is large and/or if there is gene flow. Conversely, populations could be genetically differentiated if they experienced a strong reduction in population size, even if they split recently. To identify the most likely history explaining the observed pattern of genetic differentiation between populations, we used our dataset to model the demographic history of the three populations with *fastsimcoal2* (Excoffier et al. 2013).We tested models assuming different population split times. The Akaike information criterion (AIC) indicated that the data was more probable under a model assuming that the ancestral population of SP and PL (SP, PL) split from the AUS lineage first (Fig. 1c; Table S2). Divergence between (SP, PL) and AUS (T) was estimated to have occurred approximately 292,210 generations ago (CI: 225,574 – 336,016). The split between SP and PL was estimated to have occurred more recently, approximately T = 74,042 generations ago (CI: 51,054 – 100,642). Demographic modeling further indicated that the most probable migration scenario entailed historical migration between all populations (Table S3). The model indicated that geneflow was higher between PL and AUS (PL to AUS, 4N_e_m =2.113, (CI: 1.668 – 6.771) and from AUS to PL 4N_e_m= 0.039 (CI: 0.05 – 0.125)) than between SP and PL (SP to PL 4N_e_m = 0.038 (CI: 0.013 – 1.699), and PL to SP 4N_e_m = 0.162, (CI: 0.062 – 1.924).

**Fig. 1:**
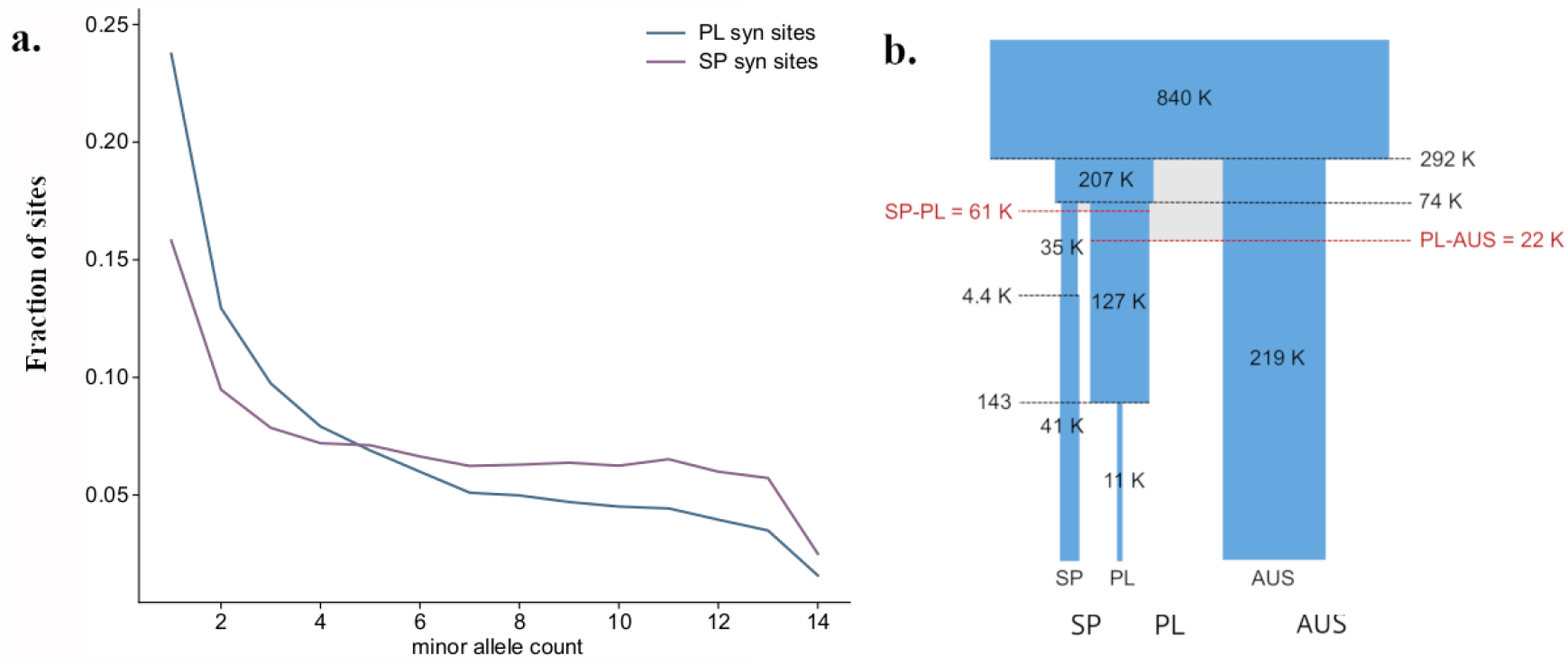
Demographic analysis of 3 *Arabidopsis lyrata* ssp. *petraea* populations. **a.** Folded site frequency spectrum of synonymous sites for PL and SP **b.** Schematic representation of the best-fit demography model. Shown within the boxes are the effective number of diploid individuals (Ne), divergence times (horizontal black lines) are indicated in thousands (k) of generations, with the exception of the final bottleneck in PL. This bottleneck is inferred to have occurred only 143 years ago but it must be noted that, in contrast to the other demographic events, it is not supported by other methods. The time since migration ended (horizontal red lines and numbers in red) is also indicated in thousands of individuals or generations. Width of the elements represents relative differences in Ne (in logarithmic scale), while time-differences in logarithmic-scale are represented by the height of the elements.

In addition, estimated effective population sizes before and after divergence events indicated bottlenecks in all populations. The size estimate of the ancestral population reached N_e_= 839,169 (CI: 823,959 – 877,924). The effective population size (N_e_) of SP was reduced approximately 6-fold after it diverged from PL, from Ne= 206,610 (CI: 100,945 – 308,029) to N_e_=35,479 (CI: 21,624 – 54,855), respectively before and after the split. In contrast, the PL population experienced a weaker initial bottleneck with N_e_ reduced by 40% after the split from SP: 127,100 (CI: 87,666 – 162,171). Both SP and PL also experienced more recent population size changes, with a slight increase in SP to a current N_e_ of 40,886 (23,081 – 47,713), approximately 4,421 (CI: 2,755 – 39,967) generations ago, and a very recent drop in PL to a current N_e_ of 11,190 (2,573 – 20,751), approximately 143 (CI: 4 – 361) generations ago. This ultimate drop in N_e_ of PL may be due to trade-offs in fitting jointly the SFS of all three populations, because it was not confirmed with other methods (see below, Fig. S2, S3). The population size in AUS decreased to 219,078 (CI: 148,664 – 249,105) after splitting from an ancestral population shared with PL. We note, however, that the AUS sample consists of individuals collected from three closely located sites, and thus might reflect diversity at a coarser grain than the SP and PL samples. We confirmed that fold-reductions in population size were robust to sample size (Table S4). We also observed a good correspondence between the observed population-specific SFS (Fig. 2a) and those simulated under the best-fit demography model, indicating that the model captures the evolutionary history of these populations reasonably well (Fig. S1c-d).

**Fig. 2:**
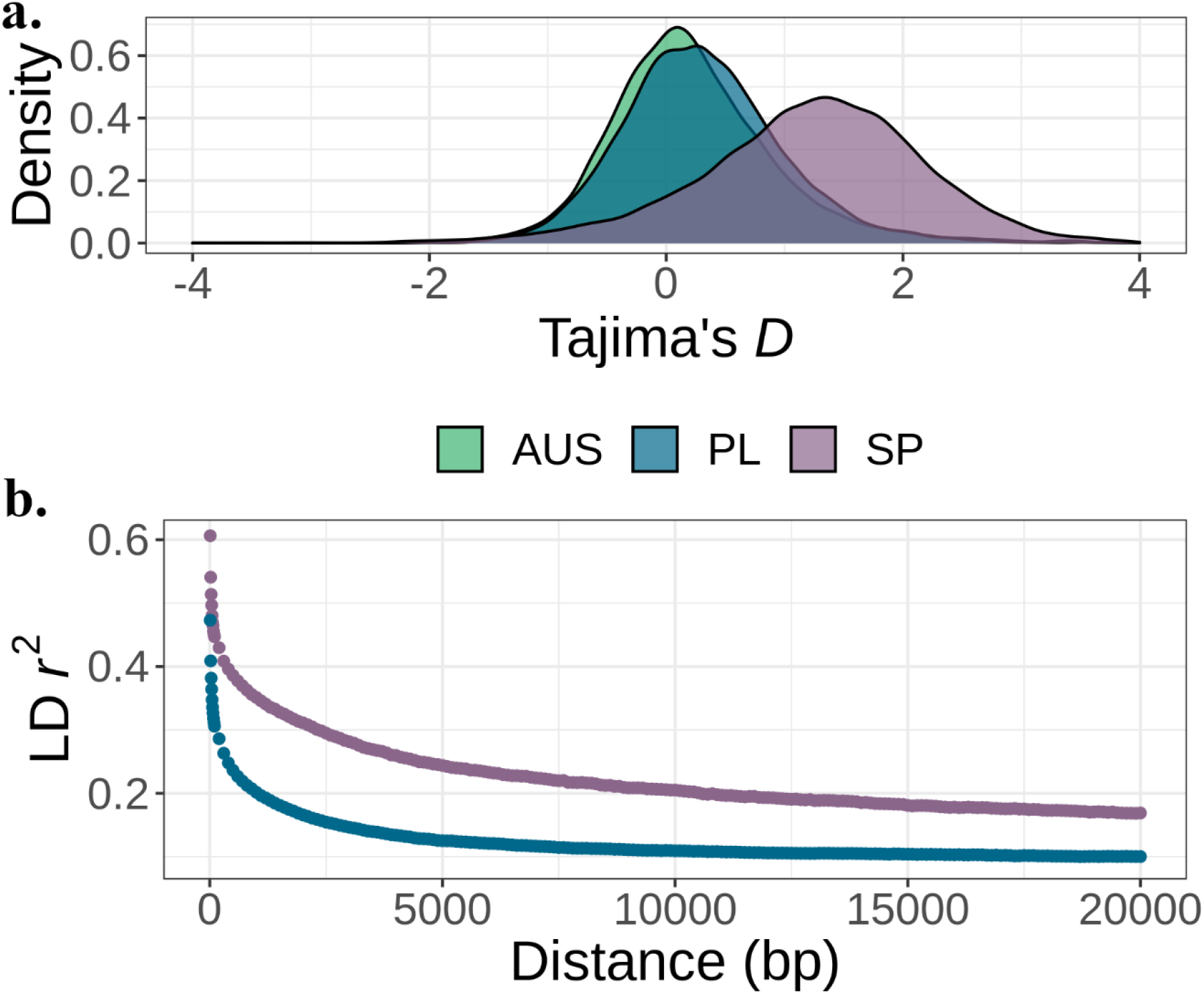
Evidence of a strong bottleneck along the SP genome. **a.** Tajima’s D distribution for AUS, PL and SP calculated along the chromosomes in 10kb non-overlapping windows. **b.** Linkage disequilibrium decay in SP and PL given by SNP pairwise r^2^ as a function of the distance between the SNPs. For comparison, both populations were down-sampled to 12 individuals each.

We calculated Tajima’s D values in 10kb windows for each population (Fig. 2b). The distribution of Tajima’s D values for SP was shifted towards positive values (mean = 1.230, median = 1.286), which was consistent with the inferred demographic history of a strong recent bottleneck in SP. Tajima’s D values for PL and AUS were also mainly positive (mean = 0.313, median 0.265 for PL and mean =0.240, median =0.151 for AUS) but both were significantly lower than in SP (Kolmogorov-Smirnov, KS test *p* < 2.2e-16 in both cases). The two distributions also differed significantly (KS test *p* < 2.2e-16).

Additionally, analysis of linkage disequilibrium (LD) decay further confirmed the stronger bottleneck experienced by the SP population. LD decay was calculated on the subsample of 12 field collected SP individuals to ensure that native LD levels were analyzed (individuals obtained from crosses in the greenhouse were removed). LD was halved within 2.2 kb in SP, which is considerably slower than for an equally sized sample of PL individuals (LD halved within 0.5kb; Fig. 2b).

Demographic modeling indicates that the large and fairly stable effective population sizes along with the persistence of gene flow for quite some time has resulted in a modest population differentiation between PL and AUS, despite their early split. By contrast, a more severe bottleneck and the lack of gene flow led to a stronger differentiation between SP and the other two populations.

### The distribution of fitness effects

To infer the efficiency of negative selection, we estimated the distribution of fitness effects of new mutations (DFE) in both range-edge (SP) and core (PL) populations, taking the demographic history into account, and investigated the range of fitness effects of mutations contributing to population differences in genomic load (Williamson et al. 2005; Boyko et al. 2008). As the AUS population had a smaller sample size, as well as individuals taken from three different local sites, it was excluded. For SP and PL, we used a modified version of the software fit∂a∂i (Kim et al. 2017). We also fit a simplified demographic model that excluded AUS to the 4-fold SFS using ∂a∂i in order to enable DFE inference (Gutenkunst et al. 2009). This model was compatible with the complex model described above but assumed a larger population size in PL to account for migration from AUS (see Methods). The demographic model showed a very good fit with (putatively neutral) SFS at 4-fold degenerate sites of both PL and SP (Fig. S2a-d). Obviously, it was not identical to the demographic model described above, which was fit to the SFS of three populations and allowed migration between demes. In particular, the very recent bottleneck inferred by FASTSIMCOAL in PL was not confirmed. Yet, both models were consistent (Fig. 1, Fig. S3). In particular, the simplified model inferred in SP also corresponded very well to the scenario expected for range-core and-margin populations in an expanding species (Fig. S2a-b). DFEs were modeled as gamma distributions and were estimated based on the nonsynonymous (0-fold) folded SFS in both populations, taking the demographic history fit using ∂a∂i into account. Using the estimated gamma distribution of effects (shape=0.213, scale=552.394, Fig. S2c-d) and the expected site frequency spectrum (SFS) for each s, we predicted, for each frequency bin in a sample the same size of ours, the proportion of variants within four ranges of selection coefficients (Fig. S2e-f, Fig. 3a). The expected strength of s among segregating variants differed between the populations. Neutral and nearly-neutral mutations were predicted to contribute to a greater proportion of variation in the PL population compared to SP, whereas mutations with a stronger s were found to contribute more to variants segregating in SP (Fig. 3a). Additionally, as a robustness check against our assumed non-synonymous mutation rate, we used a multinomial model to predict the DFE by fitting only the observed proportions in the folded 0-fold SFS. In this analysis, the DFE estimate had a vanishingly small variance and was well-described by a point mass at 2*N_anc_*s=1.2 (Fig. S4), indicating that most segregating nonsynonymous mutations in the two populations are likely to be slightly deleterious. Indeed, although the latter model ignores variation too deleterious to show up in the sample, we found that fixing the proportion of strongly deleterious new mutations to 44% provides a good fit to the observed 0-fold SFS in both populations. The 2*N_anc_*s estimate of 1.2 thus also provided a reasonable approximation to the strength of selection against mildly deleterious non-synonymous variants (Fig. S4-d).

**Fig. 3:**
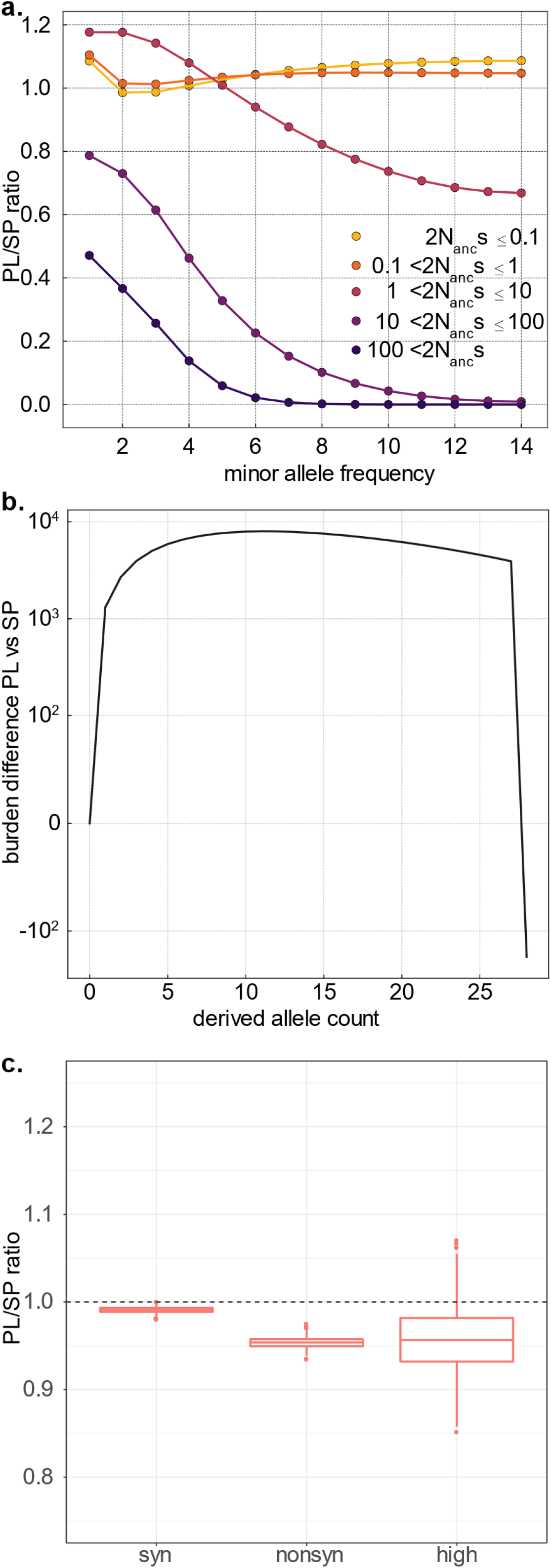
Comparative efficacy of selection and genomic burden in SP and PL. **a.** ratio of PL/SP of the proportion of variants for each s category and each allele frequency bin. Values below 1 indicate that mutations of a given size effect are less abundant in PL than in SP, within each frequency bin. This estimate is based on the joint estimate of the gamma distribution of the DFE using the Poisson optimization and the expected SFS in each category of s. As a proportion of the total number of variants at each count, PL has more slightly neutral and nearly neutral mutations (orange lines) at low frequency and considerably less strongly deleterious mutations (purple lines). **b.** Difference in per-individual cumulative derived allele burden between PL and SP. The cumulative derived allele burden is based on the contribution of deleterious variants depending on their count in the population considering the point mass s estimate of deleterious mutations of −1.2, which was shown to fit the data well. Low frequency mutations contribute more to the burden in PL – negative values indicate that an excess of up to 10 000 deleterious mutations with count 10 or less in the population accumulate in each individual in PL-, whereas fixed mutations (count 28 in the population) play an important role in SP. The net difference, given by the end of the line, is 185. **c.** Comparison of genomic load in PL and SP, for synonymous, non-synonymous and high impact mutations. For each population, the genomic load was calculated as the mean number of non-synonymous corrected by the total number of genotyped sites for each sampled individual. The ratio of mean per individual genomic load of PL vs. SP is given. The distribution was established by bootstrap of the genome (see methods).

### Estimates and Measures of accumulated burden in the range-edge population

To further investigate the effect of the severe bottleneck experienced by the range edge population SP on the deleterious load, we quantified the per-individual burden in each of the two populations. The number of derived non-synonymous mutations per Mbp of each lineage has been shown to be an appropriate proxy for the load of a population, because its expectation is unaffected by demographic events (Simons and Sella 2016).

First, we used the inferred DFE to calculate the expected burden of nonsynonymous derived mutations in each of the two populations under our demographic model. SP and PL differed in the frequency of the variants contributing to the burden (Fig. 3b, Fig. S5d). Irrespective of whether we used the gamma-distributed or point-mass DFE obtained above, modeling of the SFS suggested that low frequency mutations should contribute more to the burden in PL, the core population. We calculated that an excess of about 10,000 slightly deleterious mutations of frequency below 30% were expected in PL, compared to SP. In the latter, instead, we calculated an almost equal expected excess of fixed derived mutations in the range-edge population (SP). Fixed mutations thus played a more important role in the estimated burden of SP individuals. The predicted net difference, however, was comparatively small with an excess burden of 185 mutations per diploid genome in SP, compared to PL under the point-mass DFE (Fig. 3b). A similar burden difference was predicted using the gamma-distributed DFE (Fig. S5d). Although this number remains a crude calculation because it also depends on how we correct for the power to call SNPs in each of the two populations, it clearly indicates that the severity of the bottleneck inferred in the range-margin population SP was not sufficient to allow the accumulation of a large number of deleterious mutations in the relatively short amount of time elapsed since the split between SP and PL. This result could also be illustrated with forward simulations performed under different demographic scenarios (Fig. S6a-b). We also directly measured the accumulated burden of deleterious mutations per individual haploid genome in the range edge and core population by calculating the mean count of derived mutations per haploid genome and corrected by the total number of genotyped sites (see Methods). As expected, the mean per-individual count of derived synonymous mutations did not differ significantly between SP and PL (*p* = 0.121, Tables S5; Fig. S7). There was a shift towards a smaller average number of synonymous mutations per genome in AUS (Fig. S7), which likely reflects a residual effect of the overall lower genomic coverage of AUS individuals.

Thus, AUS individuals also had to be excluded from this analysis. For each of the other two populations, we estimated the mean count of derived non-synonymous mutations (Fig. 3c). The average burden accumulated by SP (range-edge) individuals reached a mean 0.0123 non-synonymous mutation per site (CI: 0.0118 – 0.0127). For the core population, PL, the mean burden was 0.0117 (CI: 0.0113 – 0.0121), which is 4.9% less than in SP. Permuting individuals among populations revealed that the mean difference between the two populations is significantly different from zero (*p* <10^-4^ for SP vs PL). Excluding the regions, which we inferred below as carrying signatures of selective sweeps, did not change the result (Fig. S8). Based on the approximate total of 2M non-synonymous sites per genome, we deduce that there are about 1,200 additional derived non-synonymous mutations per diploid genome in SP individuals, on average, compared to PL. Based on the estimated effect size of deleterious mutations above (point mass 2*N_anc_*s estimated to be 1.2 under the multinomial model fit shown in Fig. S4), this excess would result in a relative difference in the average fitness load of approximately 1,200 * 1.2.10^-6^ = 0.014% between the two populations.

We further used SNPeff (Cingolani et al. 2012) to identify mutations with a high deleterious impact and evaluate whether SP and PL could differ in the number of strongly deleterious mutations. Individuals in SP contained approximately 4.5% more such mutations (0.000164, CI: 0.000148-0.00018) than in PL (0.000156, CI:0.000142-0.000171, Fig. 3c; Table S5). Bootstrap across genomic regions, however, showed that this difference was not significant, with many regions in the genome showing no detectable difference in the number of mutations with high deleterious impact (*p*=0.183, Table S5). This indicates that the bottleneck would have to be either older or more severe to allow detecting a significant reduction of selection efficacy against strongly deleterious mutations. We illustrate this with forward simulation showing that under the demography inferred for SP, the burden will begin to exceed that accumulating in PL only after about 200 000 years (Fig. S6).

### Derived burden predicted to be comparatively stronger in PL for recessive alleles

Recessive mutations with deleterious effects can segregate at higher frequency in a bottlenecked population and thus lead to a genomic load in the population that is higher than predicted by measures of per-individual burden (Balick et al. 2015). Indeed, we observed an excess of heterozygous mutations in SP and PL, especially for non-synonymous and high impact variants, suggesting that homozygotes at these loci are preferentially removed from the population (Fig. S9, KS test *p* < 2.2e-16,). In order to assess whether ignoring recessive deleterious variants led our modeling efforts to underestimate the expected mutational burden in the range-edge population, we estimated the DFE as in the above, with the same demographic model but assuming this time that all derived deleterious variants were fully recessive. The best fit DFE obtained under the model with a fixed mutation rate was a gamma-distributed with shape=0.154 and scale=20396.030 (Fig. S4). In this case, we predicted an excess of approximately 2,799 mutations in PL compared to SP (Fig. S5). Indeed, recessive mutations tended to increase the number of polymorphic variants in PL that contribute to the expected burden, but had little impact on the number of fixed recessive variants, i.e. those that predominantly contribute to the per-individual burden in SP. These results indicate that, in this plant system, the recessive load is unlikely to increase the difference in individual deleterious burden between range-edge and -core populations. Forward simulations show, however, that their cumulative effect on fitness may be different in a population that experienced the decrease in population size more recently than SP did. Indeed, simulations show that there was sufficient time after the bottleneck to purge most of the recessive deleterious mutations that were frequent enough to exist as homozygotes in the SP population (Fig. S6).

### Mild load difference between SP and PL is robust to both DFE estimations and assumption on dominance relationship

Finally, we recognize that our ability to infer the magnitude of strong negative selection and degree of dominance is severely limited by the allele frequencies our sample allows to investigate (Bustamante et al. 2001, Williamson et al. 2004). To investigate the difference in derived allele burden and genomic load between the range-edge and -core populations expected under different DFEs and dominance relationships, using our demographic model (Fig. S2a-b), we calculated the expected values of these quantities for a range of (s, h) pairs (Fig. S10). These burden and load values represent the possible range under different selection scenarios. When selection is codominant, the expected excess burden in SP does not exceed 500 mutations, and the expected excess load does not exceed 0.25%. The greatest excess load in SP is predicted, for completely dominant mutations, in the range [1<2*N_anc_*s<10] and does not exceed 750 deleterious variants. For completely recessive mutations, the greatest excess load in SP is predicted in the range [6<2*N_anc_*s<40] and does not exceed 2%. These results suggest that, under our demographic model, a large difference in load for nonsynonymous mutations is not expected for any DFE and dominance relationship, and that the moderate excess derived allele burden we observe empirically does not necessarily imply an important difference in load.

### SP and PL show similar growth rate in a common garden of the species in the range core

We further investigated whether a significant fitness erosion could be detected at the phenotypic level in the range edge population. We planted six replicate cuttings of 10 genotypes of each of the two populations in the common garden of University of Cologne, at a latitude that is comparable to that experienced in the species core range. The experiment was initiated early autumn and terminated a year later at the end of the growth season. Although individuals of SP had a comparatively smaller rosette diameter after winter, the rosette diameter as well as their accumulated biomass did not differ from that of PL individuals at the end of the growth season (GLM model, *p*=0.26, and *p*=0.28, for the population effects of rosette diameter and accumulated biomass, respectively, Fig. S11), due the comparatively higher growth rate of SP individuals during the growth season (Month and Population interaction *p* < 2.2e-16). Furthermore, none of these fitness measure correlated with the per-individual burden (ρ = −0.111, *p* = 0.66 for weight; ρ = −0.149, *p* = 0.55 for diameter at end of the season), nor with the level of heterozygosity (ρ = 0.243, *p* = 0.34 for diameter at end of the season; ρ = 0.243, *p* = 0.29 for biomass), which was estimated as the inbreeding coefficient *F_IS_* These analyses show that despite its increased per-individual burden and the potential impact of recessive deleterious variants, the cumulative effect of these mutations in the SP population did not result in a detectable decrease in complex fitness component traits such as growth. This observation is in agreement with previous reciprocal transplant experiments involving the same set of *A. lyrata ssp. petraea* populations, which concluded that the SP population is locally adapted (Leinonen et al. 2009). However, it stands in strong contrast with the clear effect of range expansion detected on plant survival and population growth rate in *A. lyrata ssp. lyrata*, which has a mixed mating system (Willi et al. 2018).

### Selective sweeps in the range edge are broader than in the core but equally frequent

We searched for the footprints of selective sweeps within SP and PL – the two populations with the largest sample sizes using the Composite Likelihood Ratio (CLR) test. CLR estimates were computed in windows along the chromosomes with *SweeD* (Pavlidis et al. 2013). Significant deviations from neutral expectation were defined by comparing the observed diversity estimates to neutral diversity estimates simulated under the demographic model obtained above. We used the overlap of outlier CLR and *F_ST_* to identify putative selective sweep regions specific to SP or PL (and thus indicative of local selection). We detected signatures of local sweeps within both populations despite their large differences in recent effective population size. In SP, we identified 1,620 local sweep windows, which grouped in 327 genomic regions of average size 7051bp and they cover 0.17% of the genome (see methods). Within PL, 745 windows, covering 104 genomic regions (average size 4,384bp; 0.87% of the genome), had PL specific signatures for sweep. In both populations, the sweeps were distributed along all the chromosomes (Fig. S12, Table S12). Hence, the rate of adaptive evolution in the SP populations does not seem to have been compromised by the recent bottleneck.

Genes within the genomic regions carrying a population-specific signature of a selective sweep were extracted and tested for functional enrichment (Supplementary Information). In SP, fifteen Gene Ontology (GO) terms were enriched among genes showing signatures of positive selection (significance based on permutation derived *p* threshold of 0.0295). Interestingly, the top three GO terms were related to plant growth in response to environmental stimuli: “cellular response to iron ion”, “response to mechanical stimulus” and “response to hormone”. This observation is in agreement with the higher growth rate displayed by SP individuals in the common garden experiment. In PL, three GO enriched terms were significant (*p* threshold of 0.02137) and they were “intra-Golgi vesicle-mediated transport”, “regulation of anion transport” and “hexose metabolic process” (Table S6). Some of these functions have been associated with abiotic stress reactions in plants (Howell 2013) and may indicate adaptation in PL to the absence of snow cover protection during the cold season.

We further investigated whether specific groups of candidate genes carried signatures of adaptive evolution. Phenotypic differences in flowering time and especially selection related to the photoperiodic pathway, or to development have been shown to contribute to local adaptation in SP (Toivainen et al. 2014; Mattila et al. 2016; Hämälä and Savolainen, 2019), as well as response to abiotic factors such as cold and drought (Vergeer and Kunin 2013; Davey et al. 2018). We thus explored whether specific groups of genes associated with these traits carried signatures of adaptive evolution. We used the *A. thaliana* annotation to identify the *A. lyrata* orthologs of genes involved in these phenotypes. We then tested whether their *F_ST_* estimates tended to be higher than the rest of the annotated genes (Table S7). An excess of high *F_ST_* values was detected for genes involved in development and light (*p* = 0.018 and *p* = 0.036, respectively). Yet, genes related to dormancy, flowering, cold and water conditions did not exhibit significantly higher *F_ST_* values than the control group (Table S7).

### Negative frequency-dependent selection maintained S-locus diversity in the range-edge population

Despite a smaller effective population size in SP, strong negative frequency-dependent selection acting on the self-incompatibility locus effectively maintained or restored S-allele diversity. In SP, 15 S-alleles (allelic richness was equal to 7.6) were detected across 22 individuals, with gene diversity at the S-locus equal to 0.828. These values were only slightly lower than to those observed within the 18 PL individuals (14 S-alleles; allelic richness = 8.1; gene diversity = 0.877) and the 7 AUS individuals (10 S-alleles; allelic richness = 10.0; gene diversity = 0.940) (Table 1; Table S8). High S-allele diversity in SP (while a drastic reduction of the diversity at the S-locus would have been expected if a shift in the mating system had occurred), suggests that individuals are highly outcrossing and thus that the past bottleneck does not seem to have affected the mating system. The S-locus *F_ST_* between SP and either PL or AUS was equal to 0.027 or 0.037, respectively, values much lower than the whole genome (0.231 or 0.234, respectively) as predicted by Schierup et al. (2001).

**Table 1:** S –locus allelic diversity has been maintained within SP. The number of S-alleles for each population sample, as well as the number of individuals is provided. For each population the allelic richness has been calculated according to a rarefaction protocol with *N*=7.

| Population | S – alleles | Allelic Richness | Sample Size |
| --- | --- | --- | --- |
| SP | 15 | 7.6 | 22 |
| PL | 14 | 8.1 | 17 |
| AUS | 10 | 10.0 | 7 |

## Discussion

### Genomic burden detectable in range edge population, but little evidence of impaired fitness

The relationship between population size and selection is a centerpiece of population genetics theory (Kimura et al. 1963). At equilibrium, smaller populations have a higher genomic load that may translate into a lower adaptive potential. These premises formed a viewpoint that population bottlenecks inhibit the removal of deleterious mutations (Kirkpatrick and Jarne 2000; Hamilton 2009; Glémin and Ronfort 2013; Balick et al. 2015). However, it takes time until the equilibrium between gain and loss of mutations is restored in a bottlenecked population, so that population size reduction does not immediately associate with the presence of an increased burden of deleterious mutations (Simons et al. 2014; Do et al. 2015, Fig. S6).

The SP population provides a clear case of a range-edge population likely exposed to a severe bottleneck but with only a mild increase in average burden of deleterious mutations. Demographic modeling estimated that the population progressively decreased to about 4.8% of its initial size, despite the population growth estimated in recent generations. In agreement with previous reports (Mattila et al. 2017; Hämälä et al. 2018), this decrease had pronounced population genetics consequences: a markedly lower level of diversity, a slower LD decay and non-synonymous variants segregating at higher frequency. The genome-wide elevation of Tajima’s *D* further indicates that the population has not yet returned to equilibrium, since it is still depleted in rare alleles relative to common ones. This supports a scenario of colonization in Scandinavia with genetic material from Central European glacial refugia, a history that is common to several plant species (Clauss and Mitchell-Olds 2006; Pyhäjärvi et al. 2007; Ross-Ibarra et al. 2008; Ansell et al. 2010; Schmickl et al. 2010; Pyhäjärvi et al. 2012; Laenen et al. 2018).

Significant mutation load has been associated to post-glacial expansion in several instances, where expansion occurred along with a mating system shift. Individuals of the sister sub-species *A. l. ssp. lyrata* showed a marked increase in phenotypic load at the range edge, particularly in populations that shifted to selfing (Willi et al. 2018). In *Arabis alpina*, individuals sampled in a selfing population of the species Northern European range also appeared to have accumulated a load of deleterious mutations greater than that of populations closer to the range-core (Laenen et al. 2018). Here, we investigated the footprint left by post-glacial range expansion in populations that did not experience a shift in mating system.

To measure the per-individual genomic burden of deleterious variation, we focused on the number of derived nonsynonymous mutations in individual genomes. This metric has the considerable advantage that it is insensitive to variation in population size under neutrality (Simons et al. 2014; Do et al. 2015) and we verified it is not influenced by the presence of selective sweep areas. Other metrics, such as those which use the proportion of variation that is nonsynonymous are confounded by demographic history (Do et al. 2015; Brandvain and Wright 2016; Simons and Sella 2016; Koch and Novembre 2017).

In the range-edge population of *A. lyrata*, prediction based on the estimated DFE indicated that the differences of demographic histories of the two populations had a strong effect on the frequency of the mutations contributing to the per-individual burden. In SP, fixed mutations contributed comparatively more to the individual per-genome burden, whereas in PL, it was sustained by a greater number of low frequency mutations (Fig. 3b, assuming h=0.5). Overall, our model worked well in practice, because it provided a good fit of both the synonymous and non-synonymous SFS of both populations (Fig. S2-S4), and finally predicted an average excess of only 185 non-synonymous mutations per diploid genome in SP.

This prediction ignored the possibility that linkage with adaptive variants could have caused the faster accumulation of a burden by increasing genetic drift in genomic regions linked to selective sweeps (Marsden et al. 2016). We believe, however, that linked selection will not have a strong impact on our predictions. First, differences in per-individual burden obtained after excluding regions carrying sweep signatures were similar (Fig. S8). Second, the increased accumulation of deleterious mutations in the range-edge population is caused by nearly-neutral variants that become effectively neutral in the bottlenecked population, and the rate of fixation of neutral mutations is not expected to be affected by linked selection (Birky and Walsh 1988). Third, linked selection tends to distort allelic distribution in very large samples, because they mostly affect the low and high frequency ends of the spectrum (Cvijović et al. 2018). The effect of linked purifying selection is therefore unlikely to be important in the range of variation we can describe with our sample sizes. We note, however, that the population bottleneck could have been underestimated if we overcorrected for the reduced power to call variants due to the somewhat lower coverage of the range-edge population. This would indeed lead to an underestimation of the burden.

The empirical observation was 6-fold higher than the predicted one (~1200 vs. 184 mutations), which may indicate that we overcorrected for the reduced power to call variants in the range-edge population. We also note that some fraction of the nonsynonymous variants observed in SP are likely to be adaptive and not deleterious. Given the small average fitness effect size we predict for derived non synonymous mutations in our sample, the approximately 6-fold discrepancy between predicted and observed per-individual burden does not alter our conclusion that differences in per-individual burden between the two population is unlikely to have a strong effect on fitness.

This number of deleterious mutations per individual genome, however, remains a crude estimator. First, it underestimates the contribution of recessive deleterious mutations, which may segregate in the population even if they have large effect sizes (Balick et al. 2015). The strong deficit of homozygous large effect mutations within both populations clearly shows that recessive deleterious variants do contribute to the load in these populations. If we assume that all deleterious variants are recessive, however, our analyses showed that recessive deleterious mutations are less likely to contribute to the per-individual burden in the range-edge than in the range-core population and that their effect on the genetic load is limited (Fig. S10b). Second, indirect methods may be more powerful to reveal the extent of load differences between populations. For example, patterns of Neanderthal introgression in the modern human genome, revealed the increased deleterious load of the introgressing genome and its preferential removal in the larger *Homo sapiens* population (Juric et al. 2016). In maize, an outcrossing crop, which experienced two successive drastic bottlenecks during domestication, the variance in gene expression revealed a burden of deleterious regulatory mutations that significantly impaired fitness (Kremling et al. 2018).

A significant burden of deleterious mutations is expected to negatively impact any polygenic fitness trait, such as e.g. growth rate in plants (Leinonen et al. 2009; Debieu et al. 2013; Younginger et al. 2017). Our analysis indicated that the predicted effect of deleterious mutations is around 1.2.10^-6^ and therefore too small to allow the accumulated burden to impair fitness. This result was not substantially changed by considering a wider range of selection and dominance coefficients (Fig. S10). The lack of population difference in growth and survival observed in common gardens within the range-core area of the species both here and in a previous study, further supports the notion that SP individuals do not suffer from a massive deleterious burden (Fig. S11, Leinonen et al. 2009). Our results therefore indicate that, in this plant system, the severity and duration of the bottleneck experienced at the range-edge was not sufficient to allow the emergence of an impactful load of deleterious mutations. Unless selection is strongly recessive, the differential accumulation of load requires 2*N*s to become small in one population but not the other or mutation-selection balance will approximately hold in both. If the bottlenecked population still has a large N in absolute terms the mutations involved in load accumulation will necessarily have very weak fitness effects. This interpretation is illustrated by our forward simulations (Fig. S6). High loads are often observed in simulation studies (Gilbert et al. 2017, 2018), and severe bottlenecks are expected to have an immediate impact on the mutational load, though only from highly recessive mutations (Jarne and Kirkpatrick, 2000). In addition, the expansion load is greatly reduced when species expand along an environmental gradient, because having to adapt to local conditions slows down the pace of expansion (Gilbert et al. 2017). In this sense, the accumulated deleterious burden in SP is more similar to the consequences of the out-of-Africa bottleneck in humans, which has had substantial effects on the SFS of deleterious variation, but no detectable effect on the genetic load (Simons et al. 2014; Do et al. 2015).

### Absence of a bottleneck signature at the self-incompatibility locus

The S-locus diversity, both in terms of allelic richness and heterozygosity, was found to be only marginally lower in SP compared to PL and AUS. Similar levels of S-allele diversity were also reported for 12 Icelandic *A. lyrata* ssp *petraea* populations (Schierup et al. 2008), that share recent history with SP (Pyhäjärvi et al. 2012). This, together with the observation that homozygote genotypes are not more frequent throughout the genome, confirms that SP has maintained a functional self-incompatibility system, despite the historical genetic bottleneck. The persistence of obligate outcrossing in Scandinavian *A. l. ssp. petraea* populations has previously been discussed by Sletvold et al. (2013). Several North American populations of *A. lyrata* ssp. *lyrata*, in contrast, have shifted to predominant selfing at the species distribution edges (Mable et al. 2005; Griffin and Willi 2014). Low inbreeding depression (Willi et al. 2013) along with a reduced diversity of S-alleles (Mable et al. 2017) may have contributed to parallel breakdowns of self-incompatibility in these bottlenecked populations, as predicted by theory (Brom et al. 2020). Accordingly, loss of self-incompatibility has been frequently reported after range expansion or strong genetic bottlenecks [e.g. in *Arabis alpina* (Laenen et al. 2018), *Leavenworthia alabamica* (Busch et al. 2011) or *Capsella rubella* (Guo et al. 2009). Our result illustrates the remarkable power of negative frequency-dependent selection acting on the S-locus at promoting effective resilience against the effect of a bottleneck on allelic diversity. Similar results were found in *L. alabamica*, where a decrease in the size of the population did not associate with reduced S-allele diversity or increased mate limitation (Busch 2005). Even if allelic diversity could have been reduced at the time of bottleneck in Scandinavian populations of *A. lyrata*, theory predicts that negative frequency-dependent selection promotes higher effective migration rates at the S-locus as compared to control loci (Schierup 1998), suggesting that high allelic diversity could have also been restored subsequently by gene flow.

### Adaptive dynamics maintained in SP

Small size populations are also expected to require larger effect mutations to adapt, although these mutations are rare (Hamilton 2009). Whether a population size reduction immediately reduces adaptive evolution is, however, a complex question in the context of range expansion (Gilbert et al. 2017). If populations have to adapt locally at the range edge, the rate of geographical expansion slows down, along with the severity of the expansion bottleneck (Gilbert et al. 2017). A decrease in population size, however, increases the range of beneficial alleles that behave effectively neutrally (Lynch 2007). Searching for signals of selective sweeps in SP, after accounting for its demography, we identified 327 regions that formed outlier for both CLR and *F_ST_* statistics. In fact, the number of genomic regions displaying a signature of positive selection was greater in SP than in PL, a pattern that has also been observed among regions for *A. thaliana* populations collected in Sweden (Huber et al. 2014). However, we cannot exclude that some of the signal detected in SP could also result from the surfing of new alleles towards the range margin, which can mimic signatures of adaptive evolution and create false positive signatures of adaptation (Excoffier et al. 2009). We acknowledge that some of the selective sweep signatures could be caused by background selection. Such cases, however should be rare because theoretical work indicates that genetic signatures of selective sweeps and adaptive divergence are unlikely to be mimicked by background selection (Lynch 2007; Matthey-Doret and Whitlock 2019; Schrider 2020). Adaptive dynamic therefore appear to be maintained in SP. This agrees with basic population genetics theory showing that the fixation probability of deleterious mutation is much more sensitive to changes in population size than that of deleterious alleles (Kimura 1964; Otto and Whitlock 1997).

Functional enrichments among regions displaying signatures of local positive selection, however, indicate the presence of true positive signals. Within those regions, functions involved in the response to stress were enriched, in agreement with a previous study investigating micro-geographical patterns of local adaptation in Norwegian populations connected by gene flow (Hämälä et al. 2018; Hämälä and Savolainen 2019). We also found a significant enrichment in genes involved in light perception, a function enriched in loci differentiating the SP population from a close-by population of lower elevation (Hämälä and Savolainen 2019). Furthermore, the *F_ST_* distribution of genes related to development was significantly shifted towards higher values, a signature indicated of polygenic selection on alleles associated to this function (Foll et al. 2014; Daub et al. 2015; Stephan 2016). Previous work has documented that Scandinavian populations display differences in several traits related to growth and resource allocation, including plant size, inflorescence production and fruit production (Quilot-Turion et al. 2013; Hämälä et al. 2018). Both local and regional reciprocal transplant experiments have revealed local adaptation in this species via life history traits and growth related phenotypes (Leinonen et al. 2009; Hämälä et al. 2018). This shows that adaptive dynamics are ongoing also at smaller geographical scale in this system and is consistent with the broad genomic signals of positive selection we observed. Taken together, our analyses show that range-edge populations of the European *A. l. ssp. petraea* and its associated decrease in population size did not alter adaptive dynamics, presumably thanks to the maintenance of both outcrossing and gene flow (Gilbert et al. 2017; Hämälä and Savolainen 2019).

## Materials and Methods

### Plant Material, Sequencing and Data Preparation

Genomic sequences of *A. l. ssp. petraea* populations of 22 individuals originating from Spiterstulen in Norway (SP; 61.41N, 8.25E), 17 individuals originating from Plech in Germany (PL; 49.65N, 11.45E) and a scattered sample of 7 individuals from Austria (AUS; 47.54N, 15.58E; 47.55N, 15.59E; 47.58N, 16.9E) were used in the analysis (Fig. S1a). Details on the sequencing methodology is given in Supplementary Information.

### Analysis of population structure

Genetic diversity and differentiation along the chromosomes were calculated with PopGenome package (Pfeifer et al. 2014) in the R environment version 3.4.4 (R Core Team 2018). Specifically, we calculated pairwise nucleotide fixation index (*F_ST_*), nucleotide diversity between (d_xy_) and within population (π) in 10kb non overlapping windows for each chromosome with functions F_ST.stats, diver-sity.stats.between and diversity.stats.within, respectively (Wakeley 1996; Hudson and Wayne, 1992). In order to avoid biased *F_ST_* estimates (Cruickshank and Hahn 2014), the windows that had *F_ST_* above 0.8, dxy and π below 0.001 in at least one population comparison, were removed from the analysis. Tajima’s D was calculated with the function neutrality.stats of PopGenome. The linkage disequilibrium for the field collected SP and PL individuals was calculated along the genome with the default functions of PopLDdecay (Zhang et al. 2019) and the values were plotted in R.

Principal component analysis (PCA) of the genomic data was conducted with adegenet package (Jombart 2008) using a dataset including only every 300th SNP to reduce computational load. This reduced dataset of 233,075 SNPs was also used to calculate SNP based *F_IS_* for each population with Hierfstat (Goudet 2005) package function basic.stats (Alexander et al. 2009; Goudet and Jombart 2015). The *F_IS_* value of each gene was estimated based on the median *F_IS_* value of its SNPs, for SP and PL.

For the admixture analysis (Alexander et al. 2009) bed files were generated with PLINK (Purcell et al. 2007), which were then analyzed for clusters K=1 till K=5, with 10 iterations of cross-validation each. The clusters were normalized across runs using CLUMPAK (Kopelman et al. 2015) and subsequently they were plotted in R.

### Demography simulations

To study the demographic history of these populations, we conducted site frequency spectra (SFS) based coalescent simulations with fastsimcoal2 v2.6.0.3 (Excoffier et al. 2013). Folded 3D SFS, comprising of SP, PL and AUS individuals, was estimated from 4-fold sites with ANGSD v0.917 (Korneliussen et al. 2014), using the same filtering steps as with variant calls. We first considered models with all possible divergence orders (see Table S2), and then compared models with five different migration scenarios, guided by previous work on the SP and PL populations (Mattila et al. 2017, Hämälä et al. 2018): no migration, current migration between PL and AUS, historic migration between PL and AUS, and historic migration between all three populations (Table S3). Each model was repeated 50 times and ones with the highest likelihoods used for model selection was based on Akaike information criterion (AIC) scores. Confidence intervals were estimated by fitting the supported model to 100 nonparametric bootstrap SFS. We used these models to define effective populations sizes (*N_e_*), divergence times (*T*) and migration rates (*m*). To evaluate how the estimated demography influences measures of positive selection, we used the *N_e_, T* and *m* parameters in combination with recombination rate estimates derived from an *A. lyrata* linkage map (Hämälä et al. 2017) to generate 10,000 10 kb fragments with *ms* (Hudson 2002). These data were then used to define neutral expectations for analysis of positive selection.

Additionally, we used the program smc++ (Terhorst et al. 2017) to infer the population size and split time in the PL and SP populations. For this, we first transformed the filtered vcf files for SP and PL using the vcf2smc command. We then inferred marginal estimates for each population using the estimate function and finally estimated the joint demography using the split command. The program was run under standard settings with the addition of the changed time points using --timepoints 1 1e6.

### Estimating the distribution of fitness effects of fixed and segregating variants

For analyzing the strength of selection, vcf files were re-filtered for each population separately, as described in the section “data preparation”. This was done to retain the largest possible number of informative positions in each of the two populations. Sites with data for more than 80% of the individuals were randomly down sampled so that each position had the same number of alleles. Because the SP and PL populations differed in the number of individuals sampled, individuals in the SP population were further randomly down-sampled at each position to give the same number of alleles in both populations. The folded site frequency spectrum was determined for each population.

A modified version of the program fit∂a∂i (Kim et al. 2017) was used to estimate the distribution of fitness effects. We describe below each step of the estimation procedure. The program fit∂a∂i is an extension to the ∂a∂i program (Gutenkunst et al. 2009), which infers demographic history using a Poisson random field model for the site frequency spectrum. The Poisson random field model assumes free recombination among sites and provides a likelihood, based on classical diffusion models in population genetics, for the observed sample allele frequencies given a demographic model and strength of selection (Sawyer and Hartl 1992, Ragsdale et al. 2018). Because we only estimate the DFE using variation in PL and SP, we first fit a simplified demographic model for these populations only using ∂a∂i (Fig. S2a-b). The simplified demographic model was inferred by maximizing the composite likelihood of the folded SFS at 4-fold degenerate sites in PL and SP using the “L-BFGS-B” method and basinhopping algorithm implemented in scipy. These models provided a good fit of the predicted neutral SFS to the data (Fig. S2c-d). They were compatible with the complex 3-population model, but assumed a larger ancestral population size to account for migration from AUS. This model also indicated that the increase in population size following the last bottleneck may have been underestimated in SP (Fig. 3c). We also confirmed this expansion in SP by inferring the population size and split time in the SP and PL populations (Fig. S3) using smc++ (Terhorst et al. 2017).

After estimating the demographic parameters of SP and PL, we proceeded to the second step of our analysis and used the 0-fold SFS to fit the DFE by estimating the shape and scale parameter of a gamma distribution of selection coefficients, taking the demographic model of each population into account. The analysis was performed assuming that deleterious variants were either all co-dominant (h=0.5) or all recessive (h=0). For this, we also estimated the 4-fold-population-scaled mutation rate theta, which reached 24,000 for PL. This rate was multiplied by 2.76 to get the 0-fold mutation rate, i.e. the non-synonymous mutation rate, for PL. In all instances, the theta used for the SP population had to be constrained to thetaPL*0.74, to account for the difference in number of sites retained in each population after all sequence quality filters (see mapping pipeline described in Suppl. Information text). To estimate the DFE from the data, we used both a Poisson model (including the population scaled mutation rate, theta) and a multinomial model (without using theta) and compared the likelihood of the data along the parameter space (Fig. S4). The primary practical difference between these models is that the multinomial model only fits the DFE for variation sufficiently weakly selected to be observed in the sample. Indeed, the multinomial model only fits the proportion of alleles observed at different frequencies (the “shape” of the SFS). In contrast to the Poisson distribution, it does not consider the decrease in per-site reduction in variation compared to 4-fold sites. Strongly deleterious variation will largely be absent from our moderate sample size and therefore does not affect the shape of the SFS. After the DFE for observed variation was fit using the multinomial approach, we also estimated the fraction of strongly deleterious mutations by examining the ratio of the observed SFS to that under the multinomial DFE using the theta calculated for 0-fold sites. This ratio gives an estimate of the fraction of mutations that are sufficiently weakly selected to be observed in the sample.

The DFE describes the distribution of fitness effects of new mutations arising in a population, and as such is independent of the demographic history. It was therefore assumed to be the same in both populations. Therefore, although fit∂a∂i includes a function for finding the maximum likelihood values for DFE parameters, we had to implement a different function to fit parameters using the composite likelihood of the SFS in both populations. We calculated the likelihood using corresponding ∂a∂i functions and determined maximum likelihood parameters using Sequential Least Squares Programming as implemented in scipy. In practice, we found that the method worked well because it converged on shape parameters that allowed a good fit to the observed data (Fig. S4). The gamma DFE fit using the multinomial method converged on a point mass at a single selection coefficient, with very low variance, that provided the best fit to the observed data, when h=0.5. For h=0, the best fit yielded a gamma distribution with shape and scale parameters but no unique point estimate.

Having determined the DFE and the demographic parameters of the two populations, we proceeded to the third step of our analysis, which predicts the properties of genetic variation in the two populations. These properties follow simply from the DFE and demographic histories under the standard diffusion model. For this, we calculated the distribution of selection coefficients for variants in each count of the SFS. We first calculated the expected SFS for each selection coefficient under the demographic model using ∂a∂i functions. Then, we calculated the expected distribution of s using the python function gamma.cdf with the shape and scale parameter calculated for the joint estimate of the DFE under the Poisson model. Finally, we inferred the distribution of selection coefficients in each count of the SFS by applying Bayes’ rule. All details are given in the annotated code file provided as supplementary information.

### Predicted and observed per-individual genomic burden

In addition to predicting the distribution of selection coefficients for different frequency alleles in our sample, we also predicted the difference in derived allele burden using the expected SFS in both populations. For h=0.5, we calculated the expected derived allele burden using both between the populations by first calculating the expected burden differences using both the DFE estimated using the Poisson likelihood and that using the multinomial. Since both were nearly identical, we focus our analysis on the point mass DFE estimated using the multinomial likelihood. For each entry in the SFS we then calculated the difference in the expected count between PL and SP, weighted by their frequency in the sample to account for their probability of being present in an individual genome. Crucially, we also counted alleles that were fixed in one population but not the other. The cumulative difference over all frequencies gives the overall expected difference in the burden of derived deleterious mutations. Assuming all variants are co-dominant (h=0.5), the multinomial model converged on a single point mass for s, which describes the average s effect of deleterious mutations observed in our sample. The multinomial model, however, did not converge on a single point mass for s when variants were assumed to be recessive (h=0). We therefore also estimated the burden using the expected joint SFS under the Poisson model when assuming variants were recessive (h=0) (see annotated code file provided as supplementary information). To illustrate the load dynamics over time, we also used PReFerSim, a forward simulation program that uses the Poisson Random Field model to monitor genetic variation over time under specified demographic scenarios, dominance levels and DFE distributions (Ortega-Del Vecchyo et al. 2016). Using the demographic model inferred from the data (Fig. S2ab), we simulated over 4.10^9^ independent mutations, assuming best fit DFE estimated for h=0.5 (Point Estimate or Gamma distributed) and h=0 (Gamma distribution). We monitored the mean per-individual load in each generation by computing the weighted sum of s of all segregating and fixed alleles. We performed the simulations under three demographic model: the demography of PL (as in Fig. S2a), the demography of SP (as in Fig. S2b) and a third demographic scenario, which was identical to the scenario in SP, with the exception that the bottleneck was extended for 300 000 years, before the population returned to its ancestral size. Simulations were run from 700 000 years in the past into 300 000 years into the future.

To investigate the dependence of the difference in derived allele burden between the populations on the particular fit DFEs, we also calculated expected differences for a grid of s and h values (Fig. S10). Because the derived allele burden and genetic load are additive, these results represent the range of possible burden and load differences for all possible DFEs.

To compare theoretical predictions to the sampled genetic variation from SP and PL, we used the number of derived non-synonymous mutations per individual to quantify the observed mean genomic burden in each population (Simons and Sella, 2016). We used SNPeff (Cingolani et al. 2012) to annotate synonymous and non-synonymous sites, as well as sites with different level of high putative impact on the protein, whose ancestral state inference was done comparing to *A.thaliana* and *C.rubella* (see Supplementary Material). Then we counted their respective numbers per individual, with weight of +1 for each instance of homozygous state of the derived allele and as +0.5 for the heterozygous sites. We divided the counts by the total number of genotyped sites, in order to correct for differences in genome mapping between the individuals. The genomic load of each population was calculated as the mean of the weighted number of non-synonymous sites of individual samples. The synonymous sites were used to confirm the robustness of the analysis, as they are expected to not differ among the populations. The confidence intervals for each population, were estimated by bootstrapping with replacement of 1Mbp windows to simulate each time a whole genome (207 1Mb regions). Significance of the mean load difference between SP and PL was estimated following Simon and Sella (2016). Briefly we bootstrapped 16 1Mbp-windows of the genome with replacement and selected two random samples from the union of the two populations to create two groups of size equal to the original populations. This generated a random distribution of expected variance in the mean derived mutation counts. We used the quantile of this distribution to determine the *p* value. Note that we verified that these estimates of per-individual burdens do not change if the regions carrying sweep signatures are removed from the analysis.

### Comparative analysis of growth rate and biomass accumulation in a common garden experiment

We propagated clonally 10 genotypes from SP and 10 from PL to study growth in a common garden setting. The experiment was initiated in September 2017 and ended August 2018 and took place at the garden of the University of Cologne. Throughout the growing season (March to August) we scored monthly diameter size, in millimeters, as a proxy for vegetative growth. At the end of the growing season, we harvested the above ground material to estimate the dry to fresh weight ratio of the plants as a proxy for the plants’ biomass. The phenotypic data are provided in Table S11. Differences between the two populations were tested in R with linear mixed models using the library lme4 (Bates et al. 2015). The model included population and month of the measurement taken as fixed effects. The genotype and replicate number were included as random effects in order to correct for pseudoreplication resulting from sampling the same individuals multiple times throughout the experiment. Significance levels were estimated with a type-II likelihood-ratio-test using the function Anova, from car library (Fox and Weisberg 2019). We estimated the per individual heterozygosity level (inbreeding coefficient F) for the derived sites, using vcftools. The phenotypes of the clonal plants were averaged per genotype and correlated to F and genomic load using spearman’s rank correlation (ρ).

### Scan for selective sweeps

Areas influenced by selective sweeps were inferred by estimating composite likelihood ratios with SweeD v4.0 (Pavlidis et al. 2013). The analysis was done in 2 kb grid sizes for the SP and PL samples. As a bottleneck can easily bias CLR estimates (Jensen et al. 2007), we used data simulated under the best supported demographic model to define limits to neutral variation among the observed estimates. Estimates exceeding the 99^th^ percentile of neutral CLR values were considered putatively adaptive. We combined significant grid points within 10 kb regions to create outlier windows. Grid points that had no other outliers within 10 kb distance were removed from the analysis. Next, we examined the sweep regions in combination with regions showing elevated differentiation to find areas targeted by strong selection after the populations diverged. As with CLR, windows with *F_ST_* values above the 99^th^ percentile of their distribution were considered outliers. Genes from the regions showing higher than neutral differentiation with both CLR and *F_ST_* were extracted. Gene Ontology enrichment analysis was performed in R with the topGO package (Subramanian et al. 2005; Alexa and Rahnenfuhrer 2016). Significance of the enrichment was evaluated with Fisher’s exact test. Significance threshold was evaluated by permutating the sample’s population identity 1,000 times.

### Identification of S alleles

We genotyped individuals at the self-incompatibility locus (S-locus) with a genotyping pipeline (Genete et al. 2019) using raw Illumina reads from each individual and a database of all available sequences of *SRK* (the self-incompatibility gene expressed in the pistil) from *A. lyrata, A. halleri* and *Capsella grandiflora* (source: GenBank and unpublished sequences). Briefly, this pipeline uses Bowtie2 to align raw reads against each reference sequence from the database and produces summary statistics with Samtools (v1.4) allowing to identify alleles at the S-locus (S-alleles). Coverage statistics allow to reliably identify homozygote versus heterozygote individuals at the S-locus. Genotype data was used to compute population genetic statistics using Fstat (Goudet 1995): number of alleles per sample, sample allelic richness (a standardized estimate of the number of alleles taking into account differences in sample sizes among populations, computed after the rarefaction method described in El Mousadik and Petit 1996, gene diversity (expected heterozygosity under Hardy-Weinberg hypotheses), and *F_ST_* (unbiased estimate of the among population fixation index).

### Identification of gene functional groups

*F_ST_*, dxy and π were estimated for all genes according to the *A. lyrata* gene annotation version 1.0.37 with PopGenome and as described above for the genomic windows. Genes that had functions involved in light, cold, flowering, plant development and dormancy were determined based on the gene ontology of the sister species *A. thaliana*. To explore whether the aforementioned groups of genes had genetic differentiation estimates that were significantly different from the genome-wide background, we performed a non-parametric, two sample Kolmogorov Smirnov test (Marsaglia et al. 2003) between the gene group of interest and the rest of the genomic genes identified in *A. thaliana* and belong in a GO group (ks.test function in R).

## Data Availability

All sequence data are available in either NCBI Short Read Archive (SRA; https://www.ncbi.nlm.nih.gov/sra) or in the European Nucleotide Archive (ENA; https://www.ebi.ac.uk/ena) with accession codes: SAMN06141173-SAMN06141198 (SRA; Mattila et al. 2017), SRP144592 (SRA; Hämälä et al. 2018), PRJEB34247(ENA; Marburger et al. 2019), and PRJEB33206 (ENA; whole genome sequences generated for this project and the rest of PL sequences).

## Acknowledgements

We thank M. Nothnagel, B. Wieters, and G. Schmitz for insightful discussions and comments on the results; Diego Ortega del Vecchyo for help with PReFerSim, M.Nordborg and P.Novikova for PL sequences; B. Laenen for help with the DFE data preparation; V. Kovacova for help with inference of derived and ancestral alleles; K. Bell for lab assistance; students and gardeners at the university of Cologne for assisting with the field experiment; and Cologne Center for Genomics (Now West Germany Center) for sequencing of SP samples. This project was funded by ERC projects #648617 Adaptoscope, #648321 (Novel) and Horizon 2020 research and innovation programme [grant number ERC-StG 679056 HOTSPOT], via a grant to L.Y.. The authors also thank the Région Hauts-de-France, the Ministère de l’Enseignement Supérieur et de la Recherche (CPER Climibio), the Biocenter Oulu and the European Fund for Regional Economic Development for their financial support.

