## Supplemental Material for "Maintenance of adaptive dynamics and no detectable load in a range-edge out-crossing plant population"

**Supplementary information (Takou et al. 2020).**

**Collection of whole genome sequence data**

Lab generated seeds of SP plants and field collected seeds of PL were grown in greenhouse conditions. We sequenced 11 PL and 10 unrelated SP individuals. The rest of the sequences were obtained from previously published data. We obtained 6 PL sequences and 5 sequences of field collected SP individuals from Mattila et al. (2017) and 7 sequences of field collected SP from Hämälä et al. (2018). Seven sequences of diploid AUS individuals were provided by Marburger et al. (2019). In total, we used 22 SP, 17 PL and 7 AUS whole genome sequences in the analysis.

Read quality was checked with FastQC and the last two bases of the sequences were removed with cutadapt v1.14 (Martin 2011). Reads were mapped against the reference genome of *Arabidopsis lyrata* (Hu et al. 2011) Ensembl version 1.0 with bwa mem, options -M (Li and Durbin 2009). Only the reads mapped against the main eight chromosomes were kept in the analysis. Samtools version 1.5 (Li 2011) was used for quality filtering (view -f 3 -q30 -F 264) and remove of PCR duplicates (rmdup). Indels were realigned with Genome Analysis Toolkit (McKenna et al. 2010) version nightly-2017-12-11-1.

Single nucleotide polymorphisms (SNPs) were called with samtools mpileup, with options -E -q 30, and bcftools call (Li 2011) with options -p 0.01. Since repeat regions align poorly, they were flagged using the script generate_masked_ranges.py (https://gist.github.com/danielecook/cfaa5c359d99bcad3200 ) and subsequently were removed with bedtools subtract, version 2.25.0 (Quinlan and Hall 2010). Sites where all individuals are heterozygous for one population likely result from excessive paralogous mapping and thus were removed with a custom python script. Indels and sites that had more than two alleles, coverage less than 10, genotype quality less than 20 or quality less than 30 were filtered out with VCFtools version 0.1.5 (Danecek et al. 2011). Also, the sites that had more than 80% of the individual data of a population missing were removed. In the end, 1,878,003 SNPs were used for the downstream analysis, and the mean depth of the individuals ranged from 11.7 to 40.6 (Table S10). Compared to PL, less sites were retained after filtering in SP (1:0.74), a difference that had to be taken into account in the modeling of the expected counts of derived mutations.

**Determining ancestral and derived SNPs**

For estimating the distribution of fitness effect, an unfolded frequency spectrum had to be generated and the ancestral state of each SNP had to be determined. For this, pairwise alignments were first generated with LASTZ v1.04 (Harris 2007), (astz 32 --ambiguous=n –notransition –step=25 –nogapped) and one combined multiple alignment file for the four species was generated using mugsy v1r2.3 (Angiuoli and Salzberg 2011) with default settings. To infer ancestral state genomic information of each SNP of the *A. lyrata* was combined with base information of *A. thaliana* and *C.*

*rubella* using custom perl scripts. Ancestral state of the SNPs was inferred using the program est-sfs v2.03 (Keightley and Jackson, 2018) using the Jukes-Cantor model. At probabilities between 1 – 0.9 the major allele was assumed to be ancestral and at probabilities between 0.1 and 0 the minor allele was assumed to be ancestral. 4-fold and 0-fold positions were identified using the NewAnnotateRef.py script (https://github.com/fabbyrob/science/blob/master/pileup_analyzers/NewAnnotateRef.py) and uSFS were extracted for these positions using custom R scripts.

**Table S1**: Summary of genome wide statistics calculated along 10kb windows in each population. The mean nucleotide diversity (π), Tajima's D, $F_{IS}$, and differentiation (pairwise $F_{ST}$ above the diagonal and dxy values below the diagonal and in bold) between the 3 populations. Finally, the ratio of synonymous to non-synonymous derived alleles (Pn/Ps) are given. Whenever it is applicable, the 75$^{th}$

| **Population** | **Watterson's θ** | **Π** | **Pn/Ps** | **Differentiation (**Fst , **dxy)** | | | **TajD** | **F$_{IS}$** |
|---|---|---|---|---|---|---|---|---|
| | | | | **SP** | **PL** | **AUS** | | |
| SP | 0.00512 (0.00682) | 0.0055 (0.007) | 0.5028 | - | 0.231 (0.349) | 0.234 (0.367) | 1.23 (1.85) | -0.193 (0.130) |
| PL | 0.0093 (0.0121) | 0.0081 (0.011) | 0.4826 | **0.0094 (0.013)** | - | 0.079 (0.129) | 0.31 (0.70) | -0.04 (0.17) |
| AUS | 0.00768 (0.0102) | 0.0067 (0.0096) | 0.4612 | **0.0083 (0.012)** | **0.0079 (0.011)** | - | 0.24 (0.59) | - |

percentile of the distribution is given between parenthesis.

**Table S2:** Selection criteria for the population divergence models tested. The three possible trees were compared based on the Aikake Information Criterion (AIC). The best fitting model is shown in bold.

| **Split model** | **Log Likelihood** | **AIC** | **Akaike weight** |
|---|---|---|---|
| **(AUS, (PL, SP))** | **-1438975.068** | **2877964** | **1** |
| (PL, (AUS, SP)) | -1439477.708 | 2878969 | ~0 |
| (SP, (AUS, PL)) | -1439527.678 | 2878809 | ~0 |

**Table S3**: Selection criteria for different migration scenarios. Selection of the best fit model was done based on the Aikake Information Criterion (AIC). The best fitting model is shown in bold.

| **Migration model** | **Log likelihood** | **AIC** | **Akaike weight** |
|---|---|---|---|
| No migration | -1438975.068 | 2877964 | ~0 |
| Current migration between PL and AUS | -1439112.311 | 2878247 | ~0 |
| Historic migration between PL and AUS | -1438901.002 | 2877824 | ~0 |

| | | | |
|---|---|---|---|
| **Historic migration between all populations** | **-1438455.012** | **2876936** | **1** |

**Table S4**: Demographic parameters estimated for the complete data set are compared against two downsampled data sets. SP and PL datasets were reduced to 2/3 and 1/3 of their original sizes.

| **Parameter** | **3/3** | **2/3** | **1/3** |
|---|---|---|---|
| NSP | 40,886 | 43,607 | 46,809 |
| NSP-H | 35,479 | 30,198 | 14,935 |
| NPL | 11,190 | 9,041 | 2,606 |
| NPL-H | 127,100 | 111,881 | 59,107 |
| NAUS | 219,078 | 187,331 | 181,429 |
| NANC_SP-PL | 206,610 | 233,398 | 197,094 |
| NANC_PL-AUS | 839,169 | 863,225 | 838,472 |
| TSP-H | 4,421 | 3,390 | 4,134 |
| TPL-H | 143 | 312 | 676 |
| TISOL_PL-AUS | 22,308 | 27,453 | 4,912 |
| TISOL_SP-PL | 61,148 | 59,642 | 20,622 |
| TSP-PL | 74,043 | 65,341 | 33,507 |
| TPL-AUS | 292,210 | 244,191 | 335,141 |

$N$ = effective number of diploids ($N_e$)

$T$ = time in number of generations

$M$ = population migration rate $4N_em$

$N_{\text{SP-H}}$ and $N_{\text{PL-H}}$ denote historic population sizes after population split, and $T_{\text{SP-H}}$ and $T_{\text{PL-H}}$ are times since population sizes changed to current estimates

$T_{\text{ISOL}}$ are times since migration ended

**Table S5**: Genomic load per population for synonymous, nonsynonymous and high impact mutations as defined by SNPeff. Confidence intervals are provided within the parenthesis (see methods). The excess of mutations in SP was estimated by multiplying the genomic load difference by the number on non-synonymous sites in the *A. lyrata* genome as estimated by SNPeff.

| **Type of Mutations** | **Total Number of Derived Positions** | **Genomic load in SP** | **Genomic load in PL** | **Genomic load Ratio of SP to PL** | **P-value** | **Excess of mutations in SP** |
|---|---|---|---|---|---|---|
| synonymous | 125228 | 0.0245 (0.0237-0.0252) | 0.0243 (0.0235-0.0251) | 1.008 | 0.121 | 400 |
| non-synonymous | 77781 | 0.0123 (0.0118-0.0127) | 0.0117 (0.0113-0.0121) | 1.049 | 0.00099 | 1,200 |
| high impact | 1323 | 0.000164 (0.000148-0.0001800) | 0.000156 (0.000142-0.000171) | 1.099 | 0.183 | 16 |

**Table S6:** Gene Ontology (GO) categories that are significantly enriched for genes located within areas with signature of selective sweep in SP or PL. The GO ID number, the description of the term and the p value of the Fisher's exact test (topGO) are provided. The p value threshold for determining the significance of the enrichment analysis was set by randomly picking genes to belong in area with signature of selective sweep, as described in the material and methods. Non significant GO.IDs are not shown.

| **GO.ID** | **Term** | ***p*** | **Population** |
|---|---|---|---|
| GO:0071281 | cellular response to iron ion | 0.0043 | SP |
| GO:0009612 | response to mechanical stimulus | 0.0066 | SP |
| GO:0009725 | response to hormone | 0.0080 | SP |
| GO:0009743 | response to carbohydrate | 0.0086 | SP |

| | | | |
|---|---|---|---|
| GO:1901699 | cellular response to nitrogen compound | 0.0094 | SP |
| GO:0042592 | homeostatic process | 0.0150 | SP |
| GO:0019725 | cellular homeostasis | 0.0162 | SP |
| GO:0090304 | nucleic acid metabolic process | 0.0165 | SP |
| GO:0009581 | detection of external stimulus | 0.0200 | SP |
| GO:0046483 | heterocycle metabolic process | 0.0203 | SP |
| GO:0016070 | RNA metabolic process | 0.0246 | SP |
| GO:0042991 | transcription factor import into nucleus | 0.024 | SP |
| GO:0006338 | chromatin remodelling | 0.0247 | SP |
| GO:1901360 | organic cyclic compound metabolic processes | 0.0260 | SP |
| GO:0050801 | ion homeostasis | 0.0261 | SP |
| GO:0006891 | intra-Golgi vesicle-mediated transport | 0.0024 | PL |
| GO:0044070 | regulation of anion transport | 0.0063 | PL |
| GO:0019318 | hexose metabolic process | 0.0141 | PL |

**Table S7:** Comparative analysis of $F_{ST}$ distributions for genes grouped based on candidate adaptive traits in the populations, or based on functions related to environmental differences between SP and PL, compared to the rest of the genome. The adjusted p values for multiple KS tests are provided.

| **Gene Group** | **Control Group** | **Adjusted *p*** |
|---|---|---|
| Flower | Athaliana GO annotated | 0.235 |

| | | |
|---|---|---|
| Cold | Athaliana GO annotated | 0.331 |
| Water | Athaliana GO annotated | 1 |
| Light | Athaliana GO annotated | 0.0364 |
| Development | Athaliana GO annotated | 0.0183 |
| Dormancy | Athaliana GO annotated | 0.899 |

**Table S8:** List of genotypes at the S-locus obtained by applying the NGSgenotyp pipeline to raw Illumina reads data. The following unpublished *A. lyrata* S-allele sequences were obtained: AlSRKnew1 (close to AhSRK07); AlSRKnew2 (close to CgSRK57); AlSRKnew3 (new allelic lineage); AlSRKnew4 (close to AhSRK47); AlSRKnew5 (new allelic lineage); AlSRKnew6 (close to AhSRK38); AlSRKnew7 (close to AhSRK13); AlSRKnew8 (close to AhSRK41); AlSRKnew9 (close to CgSRK59); AlSRKnew10 (close to AhSRK20); AlSRKnew11 (close to CgSRK20); AlSRKnew12 (close to AhSRK15).

| Individual | *SRK* allele 1 | *SRK* allele 2 |
|---|---|---|
| *Spiterstulen* | | |
| SP 1 | AlSRK18 | AlSRK01 |
| SP 2 | AlSRK16 | AlSRKnew9 |
| SP 3 | AlSRKnew10 | AlSRK01 |
| SP 4 | AlSRK01 | AlSRK34 |
| SP 5 | AlSRK42 | AlSRK16 |
| SP 6 | AlSRK01 | AlSRK15 |
| SP 7 | AlSRK05 | AlSRKnew11 |
| SP 21 | AlSRK37 | AlSRK01 |
| SP 154 | AlSRK01 | AlSRK01 |
| SP 164 | AlSRK01 | AlSRK01 |
| SP 3 | AlSRK01 | AlSRK16 |
| SP 76 | AlSRK01 | AlSRKnew10 |
| SP 70535 | AlSRK37 | AlSRK42 |

| | | |
|---|---|---|
| SP 70536 | AlSRK37 | AlSRK01 |
| SP 70537 | AlSRK10 | AlSRK01 |
| SP 70538 | AlSRK37 | AlSRKnew9 |
| SP 70539 | AlSRK15 | AlSRK21 |
| SP 70540 | AlSRK42 | AlSRK35 |
| SP 70541 | AlSRK01 | AlSRK37 |
| SP 70542 | AlSRK18 | AlSRK42 |
| SP 70543 | AlSRK01 | AlSRK01 |
| SP 70544 | AlSRK38 | AlSRK01 |
| SP 70545 | AlSRK01 | AlSRK42 |
| *Plech* | | |
| a1 | AlSRK18 | AlSRK01 |
| 2a | AlSRK18 | AlSRK01 |
| 3a | AlSRK01 | AlSRK34 |
| 4a | AlSRK39 | AlSRKnew1 |
| 5a | AlSRK18 | AlSRK18 |
| 6a | AlSRK34 | AlSRKnew2 |
| 7a | AlSRKnew6 | AlSRKnew3 |
| 8a | AlSRKnew4 | AlSRK01 |
| 9a | AlSRKnew5 | AlSRK18 |
| 10a | AlSRK01 | AlSRKnew6 |
| 11a | AlSRK34 | AlSRK34 |
| PL S2 | AlSRK01 | AlSRK01 |
| PL S4 | AlSRK28 | AlSRKnew4 |
| PL S5 | AlSRK01 | AlSRKnew1 |
| PL S6 | AlSRK27 | AlSRK01 |
| PL S7 | AlSRKnew7 | AlSRKnew5 |
| PL S8 | AlSRK01 | AlSRK01 |
| PL 80936 | AlSRKnew8 | AlSRK34 |
| *Austria* | | |

| | | |
|---|---|---|
| AUS PEQ6 | AlSRK01 | AlSRK13 |
| AUS PEQ9 | AlSRK08 | AlSRKnew12 |
| AUS PER08 | AlSRK01 | AlSRK38 |
| AUS PER11 | AlSRK01 | AlSRK15 |
| AUS VLH2 | AlSRK17 | AlSRK17 |
| AUS VLH5 | AlSRK46 | AlSRK19 |
| AUS VLH6 | AlSRK17 | AlSRK10 |

**Table S9:** Summary of distribution of distances (in bp) between two consecutive SNPs with $F_{IS}$ value equal to -1 in the PL, SP collected from the field and SP amplified in the greenhouse individuals.

| | **PL** | **SP collected in the field** | **SP amplified in the greenhouse** |
|---|---|---|---|
| **Min** | 322 | 301 | 300 |
| **1st Quartile** | 574,658 | 41,744 | 28,271 |
| **Median** | 1,528,536 | 172,937 | 129,204 |
| **Mean** | 2,120,124 | 324,666 | 257,872 |
| **3rd Quartile** | 3,604,405 | 417,027 | 314,092 |
| **Max** | 10,011,347 | 3,354,800 | 3,581,874 |

**Table S10:** Mean depth of each sample in the filtered dataset.

| Sample Name | Average Depth |
|---|---|
| *Spiterstulen* | |
| SP 70535 | 18.39 |
| SP 70536 | 16.47 |
| SP 70537 | 15.41 |
| SP 70538 | 15.74 |
| SP 70540 | 16.59 |
| SP 70541 | 16.67 |
| SP 70542 | 16.42 |
| SP 70543 | 17.78 |
| SP 70544 | 16.90 |
| SP 70545 | 15.32 |
| SP 154 | 18.34 |
| SP 164 | 34.48 |
| SP 1 | 24.03 |
| SP 21 | 23.98 |
| SP 2 | 21.36 |
| SP 3 | 29.51 |
| SP 4 | 32.75 |
| SP 5 | 26.98 |
| SP 6 | 12.43 |
| SP 76 | 35.27 |
| SP 7 | 22.62 |
| *Plech* | |
| PL 1a | 29.54 |
| PL 2a | 14.03 |

| | |
|---|---|
| PL 3a | 24.9 |
| PL 4a | 30.66 |
| PL 5a | 15.02 |
| PL 6a | 27.20 |
| PL 7a | 29.38 |
| PL 8a | 20.84 |
| PL 9a | 20.96 |
| PL 10 | 22.95 |
| PL 80936 | 27.22 |
| PL S2 | 31.48 |
| PL S4 | 30.59 |
| PL S5 | 36.41 |
| PL S6 | 38.32 |
| PL S7 | 34.99 |
| PL S8 | 40.61 |
| *Austria* | |
| AUS PEQ6 | 11.70 |
| AUS PEQ9 | 12.17 |
| AUS PER11 | 10.22 |
| AUS PER8 | 18.76 |
| AUS VLH2 | 14.67 |
| AUS VLH5 | 22.88 |
| AUS VLH6 | 21.58 |

Table S11: Excel table reporting the fitness measures in the common garden experiment.

Table S12: Excel table listing regions in the genome carrying a sweep signature in range-edge population SP and range-core population PL.

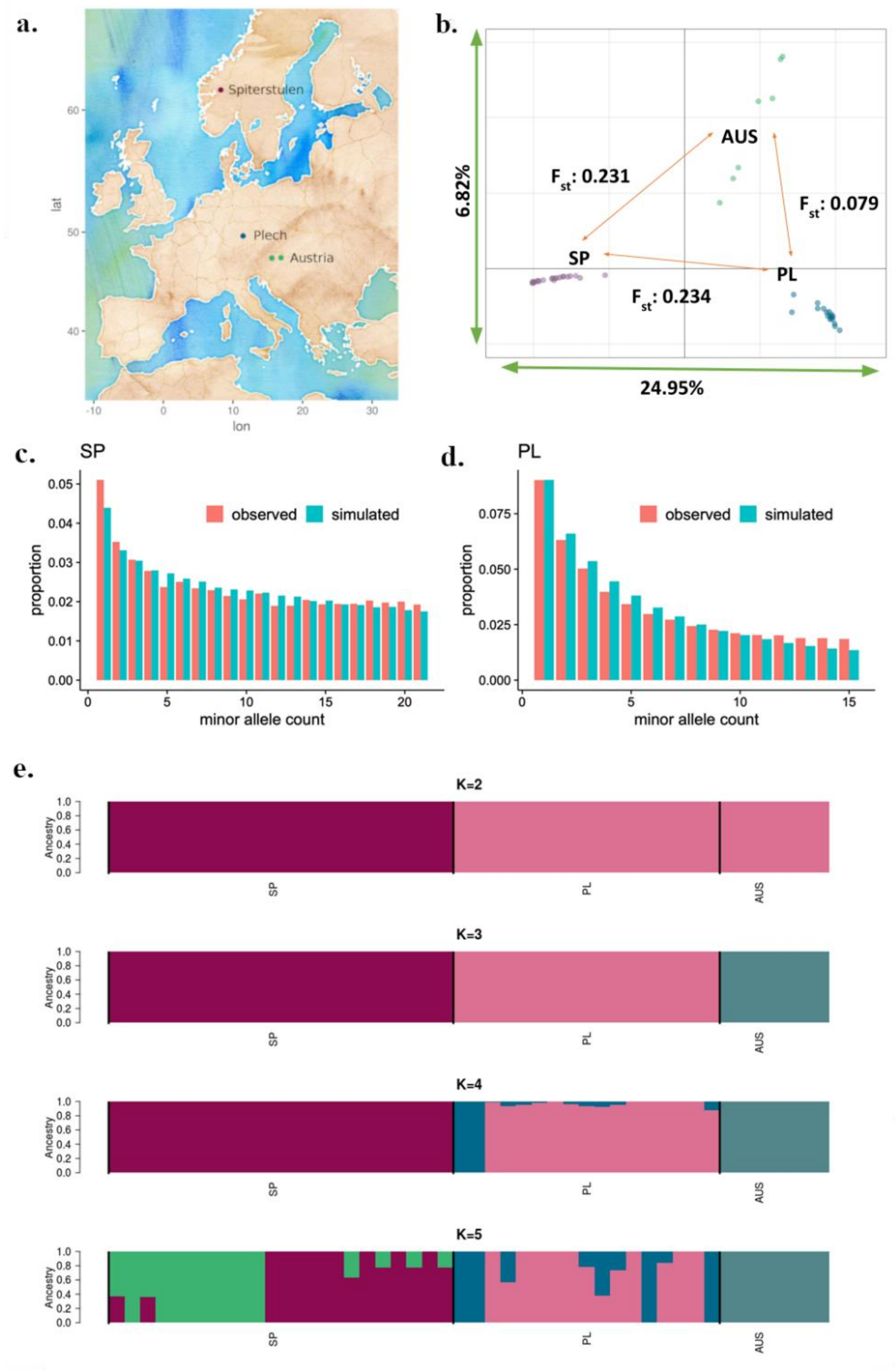


**Fig S1**: Population differentiation of 3 *A. lyrata* populations **a.** Geographical distribution of the Spiterstulen (SP), Plech (PL) and Austrian (AUS) populations. **b**. Principal Components analysis of SP, PL and AUS. The first Principal Component (PC) explains 24.95% of the sample variation and the second PC explains 6.82%. Within the PCA plot the $F_{ST}$ values between all the population pairs are given. **c-d.** Observed and estimated site frequency spectra of SP and PL used for the *fastsimcoal* analysis. **e.** Admixture analysis results for all samples. From top to bottom, the clustering in 2, 3, 4, or 5 clusters is shown. The samples are arranged in the same order in all five panels, with SP samples shown first, then PL and lastly AUS. According to the cross-validation error the most probable clustering is the K=2, and second best the K=3.

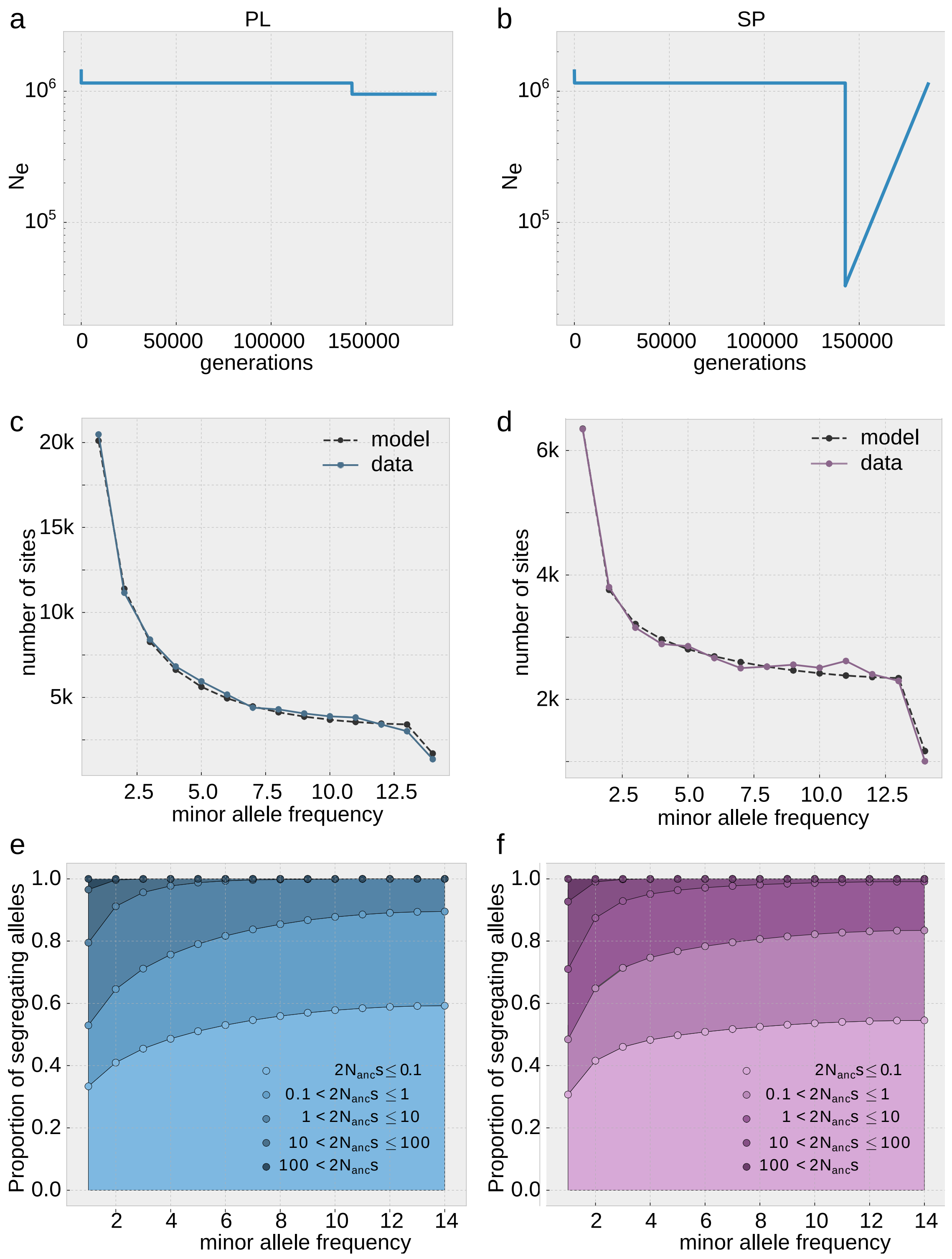


**Fig S2:** Simplified and re-fit demographic model used to assess the gamma distribution of the distribution of fitness effects for (a) PL and (b) SP. The simplified model in PL is a step wise population change and in PL shows a strong bottleneck following population expansion. (c) and (d) show the site frequency spectrum for synonymous sites. The solid line shows the data, the dashed line the estimate based on the model. The distribution of selection coefficients for variants in each category of the SFS for PL (e) and SP (f) based on the joint gamma distribution using the Poisson optimization and the expected SFS in each category of s estimated (for comparison e-f are also shown in Fig. S6).

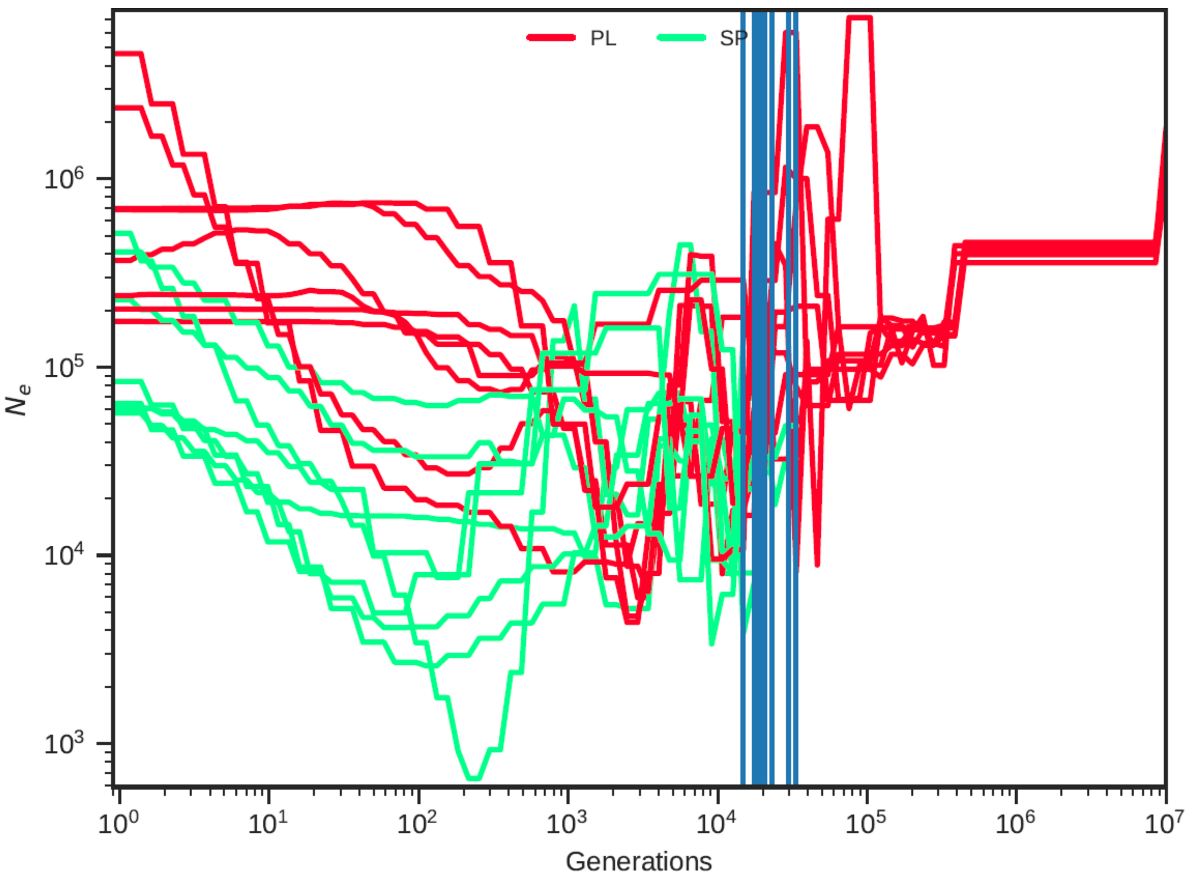


**Fig S3**: Estimated population size over time in PL and SP based on smc++. Each scaffold was estimated independently and is indicated by one line. Blue vertical lines show the estimated split time between the populations.

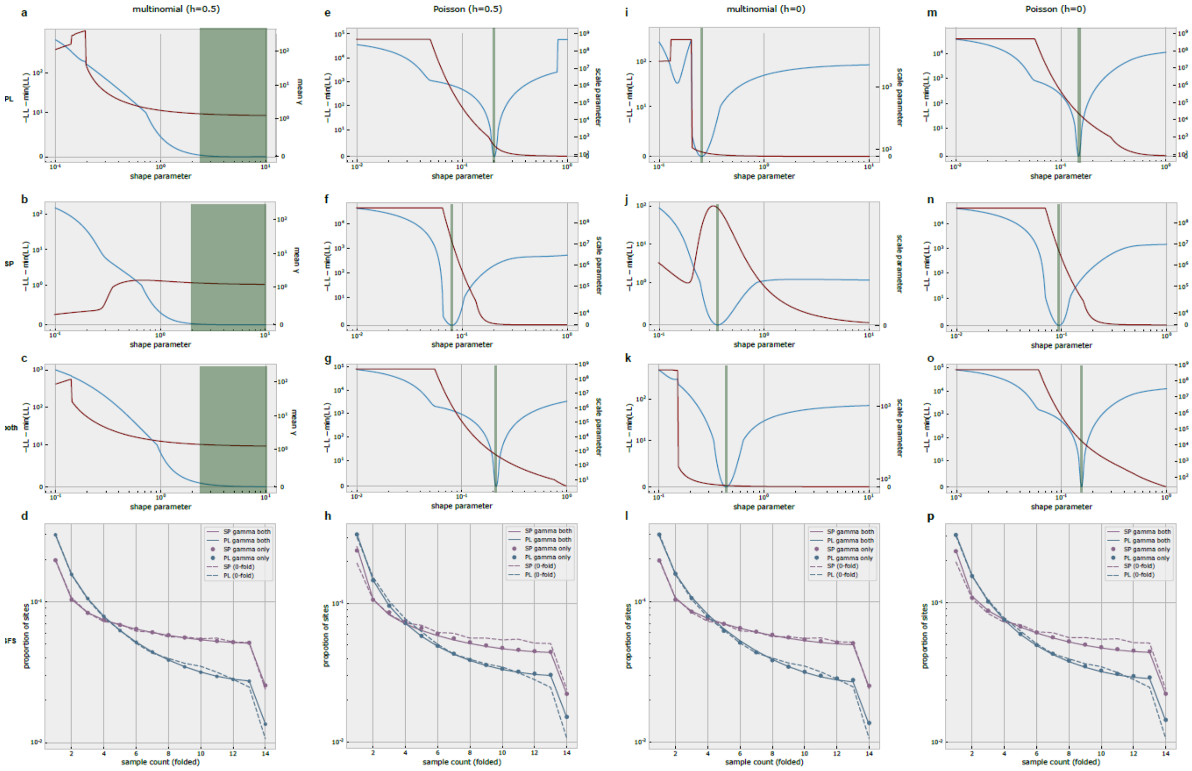


Fig S4: The shape and scale parameter of the gamma distribution were estimated assuming co-dominance h=0.5 (a-h) or complete recessivity h=0 (i-p), by fitting the 0-fold SFS with either a multinomial model that fits the SFS for mutations observed *in* the sample (a-d, i-l) or a Poisson model, which includes the value of theta estimated from the 4-fold SFS (e-f, m-p). Estimations were conducted by fitting either the 0-fold SFS of each single population (PL: first row, SP: second row) or the 0-fold SFS of both populations (third row). The shape parameter is plotted against the likelihood (blue line, left axis) as well as against the mean gamma (shape * scale) or the scale parameter (red line, right axis). This is because when the model converges on a point mass estimate, it is more informative to plot the mean gamma. Values on the red curve that provide the best fit to the data are those where the blue curve reaches its minimum (green shading). The last row shows the site frequency spectrum for the data (dashed lines) as well as under the optimized gamma estimated either for both populations (solid lines) or separately for each population (dots). Note that solid lines and dots overlap almost perfectly indicating that the two approaches are equally efficient.

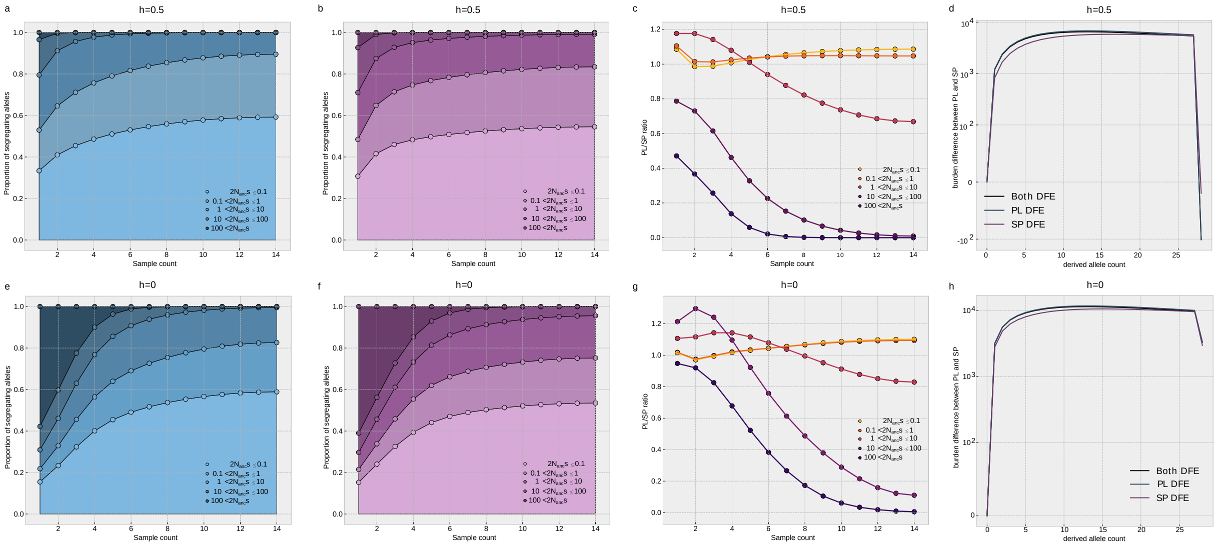


**Fig S5: Comparative efficacy of selection and genomic burden in SP and PL assuming all deleterious variants are co-dominant (h=0.5, a-d) or recessive (h=0, e-h).**

Distribution of selection coefficients for deleterious variants of each size category in each frequency bin of the SFS for PL (a,e) and SP (b,f) based on the joint gamma distribution and the expected SFS in each category of s estimated in fit∂a∂i assuming h=0.5 (top) or h=0 (bottom). Graphs c and g show the ratio of PL/SP of the proportion of variants for each s category and each allele frequency bin. Values below 1 indicate that mutations of a given size effect are less abundant in PL than in SP, within each frequency bin. This estimate is based on the joint estimate of the gamma distribution of the DFE using the Poisson optimization and the expected SFS in each category of s. As a proportion of the total number of variants at each count, PL has more slightly neutral and nearly neutral mutations (orange lines) at low frequency and considerably less strongly deleterious mutations (purple lines). For h=0, the proportion of strongly deleterious variants in PL has increased at low frequency, but does not change much at high frequency. Graphs d and h display the difference in per-individual cumulative derived allele burden between PL and SP. Low frequency mutations contribute more to the burden in PL – negative values indicate that an excess of up to 10 000 deleterious mutations with count 10 or less in the population accumulate in each individual in PL-, whereas fixed mutations (count 28 in the population) play an important role in SP. The net difference, given by the end of the line, based on the DFE from the joint gamma estimate using the Poisson model is -103 for h=0.5 (d) and +2799 for h=0 (h). Note that (a) and (b) are also shown in Fig. S2.

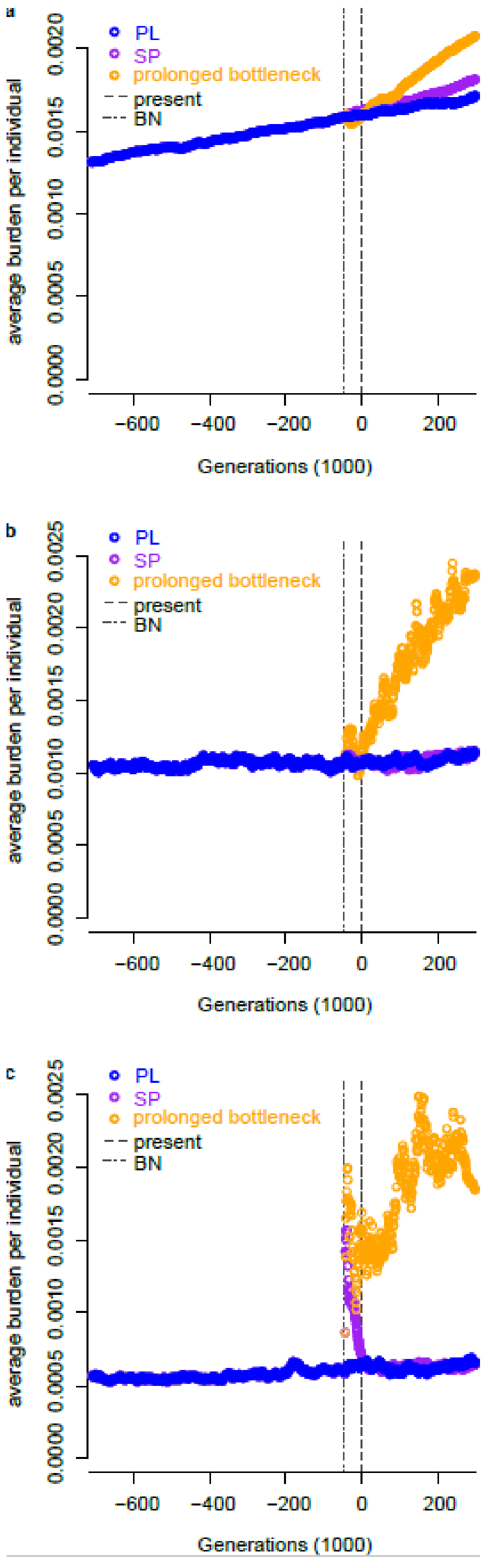


Fig. S6: We used PRefeRSim (Ortega-Del Vecchyo et al. 2016) to run forward simulation under three demographic models. The program simulates independent mutations under any given demography and DFE. The model inferred for PL (blue), the model inferred for SP (purple) and a model assuming the demography of SP, but with a bottleneck extended to 300000 years. The simulations were run assuming h=0.5 and either the point estimate DFE (a) or the gamma distributed DFE (b), or assuming that h=0 and using the corresponding gamma distributed DFE(c). Values for the Point mass or Shape and scale parameters were the same as reported in the results section. Forward simulations were run with a burn-in of 2 million generations and scored for 600 000 years before present and 300 000 years after present. In total, approximately $4.10^{9}$ unlinked mutations were simulated for each parameter set. We see that the bottleneck in SP is too recent to allow the accumulation of a significant deleterious burden of co-dominant mutations but old enough so that most recessive deleterious variants were purged. We also observe that a prolonged bottleneck (300 000 years) would lead to an increased recessive burden, but would have no effect on the burden due to co-dominant recessive mutations.

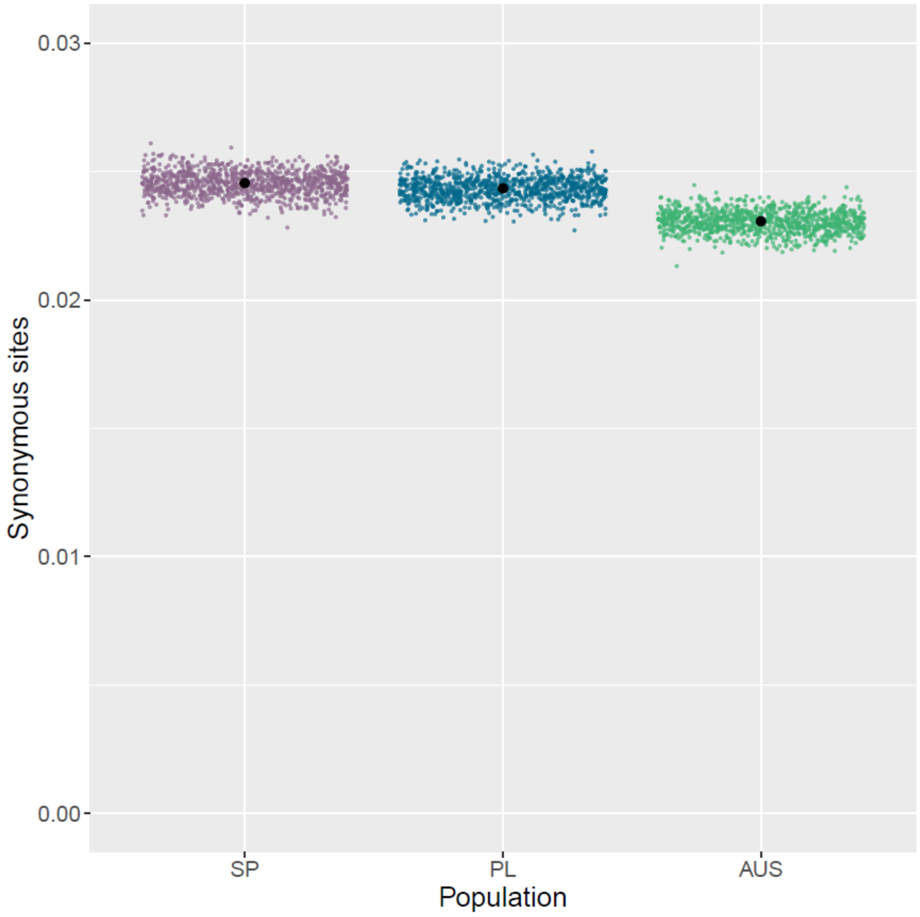


**Fig S7:** The number of synonymous sites corrected by the total number of genotyped sites per sample for each population. For each population, the mean obtained during each bootstrap iteration is shown in color and the original mean is marked in black. The expectation is that all three populations should show no differences among the number of accumulated synonymous sites. The discrepancy noted between AUS and the other populations is the result of lower genome wide coverage, which lessened the power to detect derived mutations in this population.

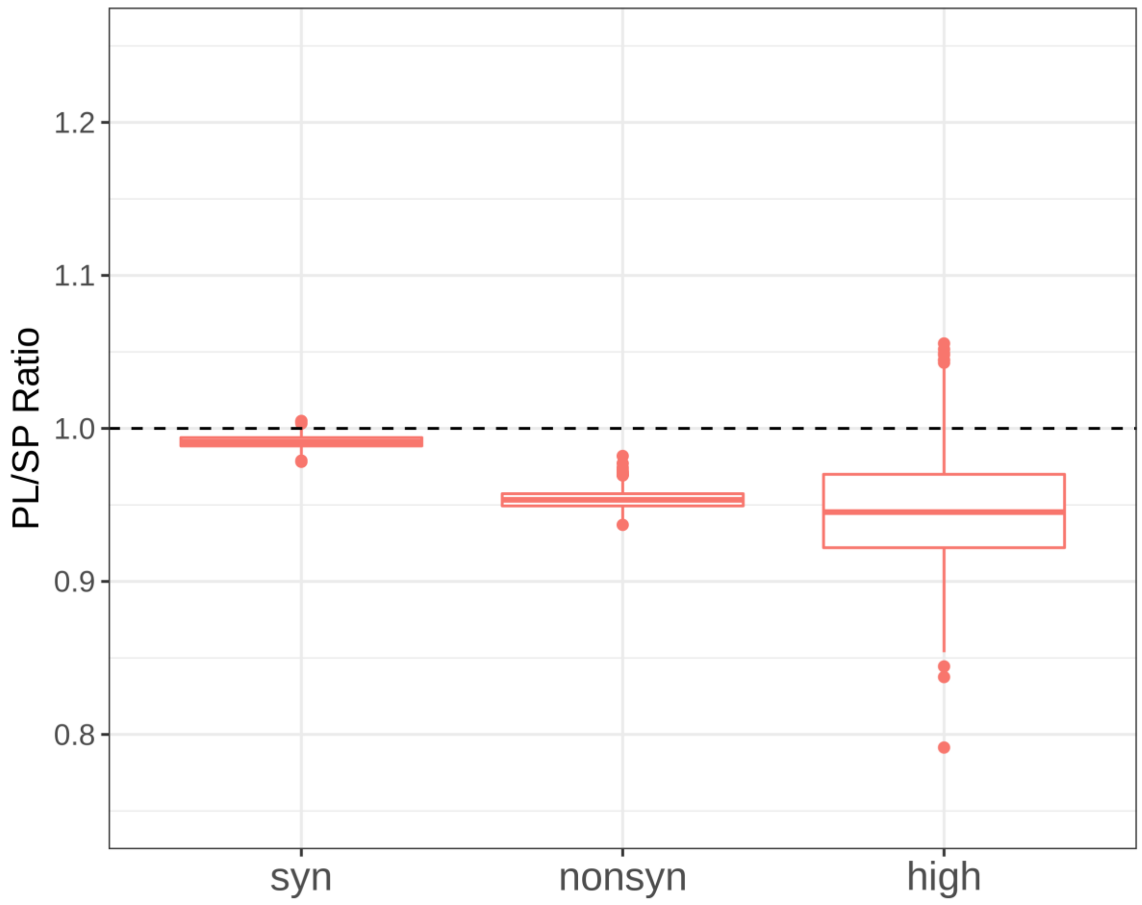


**Fig S8:** Comparison of genomic load in PL and, for synonymous, non-synonymous and high impact mutations, when the areas with signatures of selective sweep have been removed. The values per category were not altered drastically compared to Fig 3c, which includes all the derived sites. For each population, the genomic load was calculated as the mean number of non-synonymous corrected by the total number of genotyped sites for each sampled individual. The ratio of mean per individual genomic load of PL vs SP is given, for 1000 permutations.

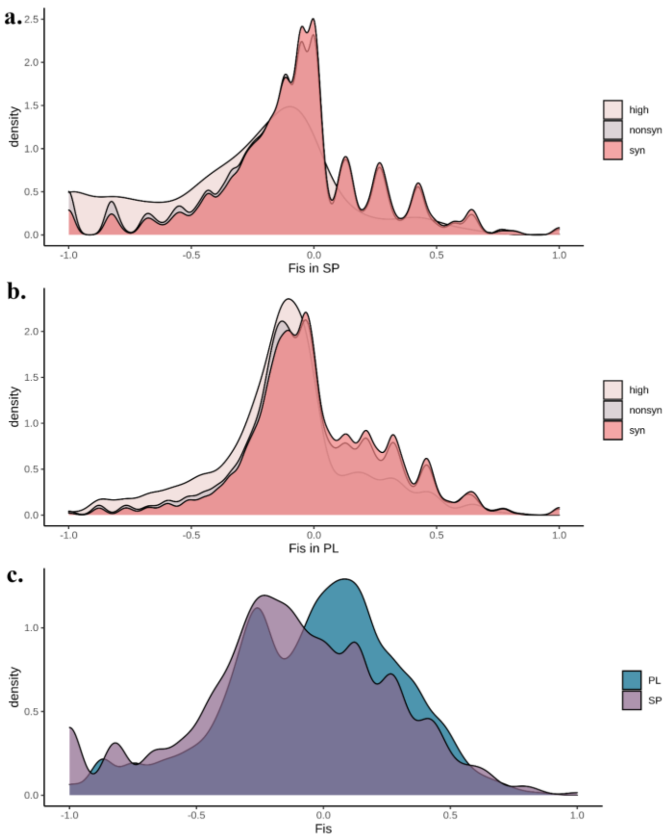


**Fig S9: FIS distribution of SP and PL.** To evaluate whether recessive deleterious mutations may contribute to the genomic load in SP and PL, we tested whether the $F_{IS}$ distribution of non-synonymous mutations (grey) showed a departure from Hardy Weinberg expectations indicative of a selective removal of individuals that were homozygote for deleterious variants. We found that both in range edge (a) and core (b) populations, the $F_{IS}$ distribution of non-synonymous mutations (grey) was significantly shifted towards lower values, compared to the $F_{IS}$ distribution of synonymous mutations (dark pink), revealing an excess of heterozygous non-synonymous mutations (a--b, KS test $p < 2.2e\text{-}16$). This effect was even more pronounced when comparing $F_{IS}$ for high impact variants (light pink) to $F_{IS}$ for synonymous mutations (dark pink, KS test $p < 2.2e\text{-}16$). This pattern suggests that, in both populations, offsprings homozygous for deleterious alleles tend to be removed by selection. (c) Compared to PL (blue), however, the $F_{IS}$ distribution of all variants in SP (purple) was shifted towards negative values (Fig. S9c, $p < 2.2e\text{-}16$). It therefore suggests that the preferential removal of recessive homozygous might be more important in SP.

**Robustness of $F_{IS}$ estimates to possible mapping errors**: We cannot fully rule out that this effect is not due to mapping inaccuracies (see below), but it was independent from coverage thresholds or SNP density. We verified that this result was not influenced by unanticipated mapping biases by using the mean read coverage of each population and the distribution of genic coverage to set depth read filters. Then, for each filter, we correlated the new gene $F_{IS}$ values with the median gene depth. Spearman's rho was in the range of -0.125 to -0.145 for SP and -0.126 to -0.164 for PL. Filtering stringency did not modify the correlation, indicating that the $F_{IS}$ bias that we specifically observe in SP is independent of read depth. In addition, we observed that SNPs with $F_{IS} = -1$ were not clustered in the genome, as would be expected from paralogous mapping. In contrast, the distribution of the physical distance between 2 consecutive such SNPs was significantly shifted towards higher values than two consecutive SNPs with any $F_{IS}$ value (KS test for each population $p < 2.2e\text{-}16$).

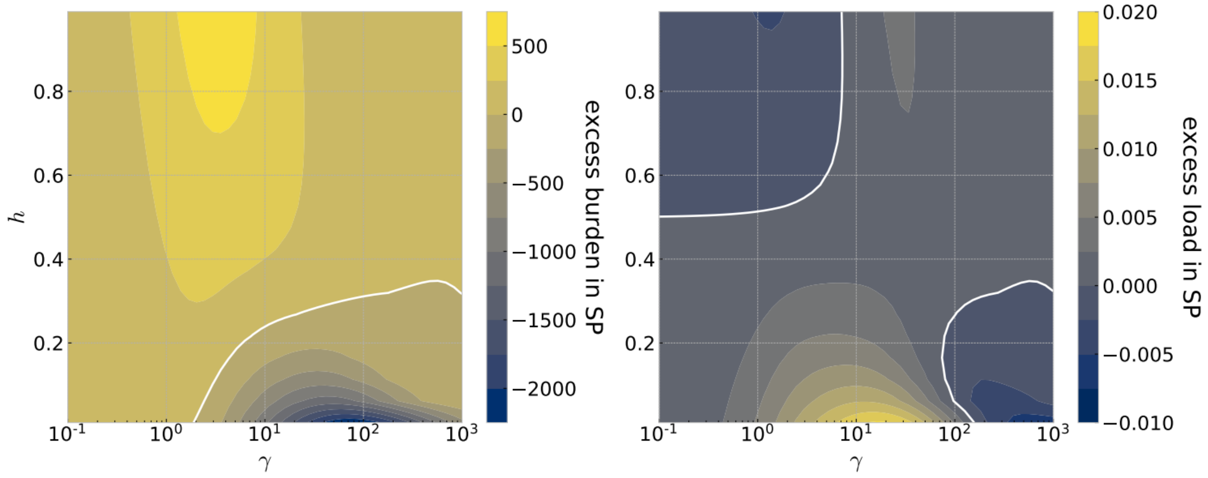


**Fig S10:** Expected excess derived allele burden (maximum 700 mutations) and genetic load (maximum 2%) in SP relative to PL for a range of selection and dominance coefficients. Expected derived allele burdens (left) and genetic loads (right) for nonsynonymous mutations were calculated from allele frequency spectra computed using ∂a∂i for the demographic models shown in Fig. S2a-b. The genetic load was approximated as $L=\sum 2s(h \times x_1+x_2)$ where $x_1$ is the expected number of heterozygous genotypes and $x_2$ is the expected number of homozygous derived genotypes. γ is twice the product of the effective population size of the ancestral population and the selection coefficient ($2*N_{anc}*s$). The population scaled rate of nonsynonymous mutations in the ancestral population is 66,379.

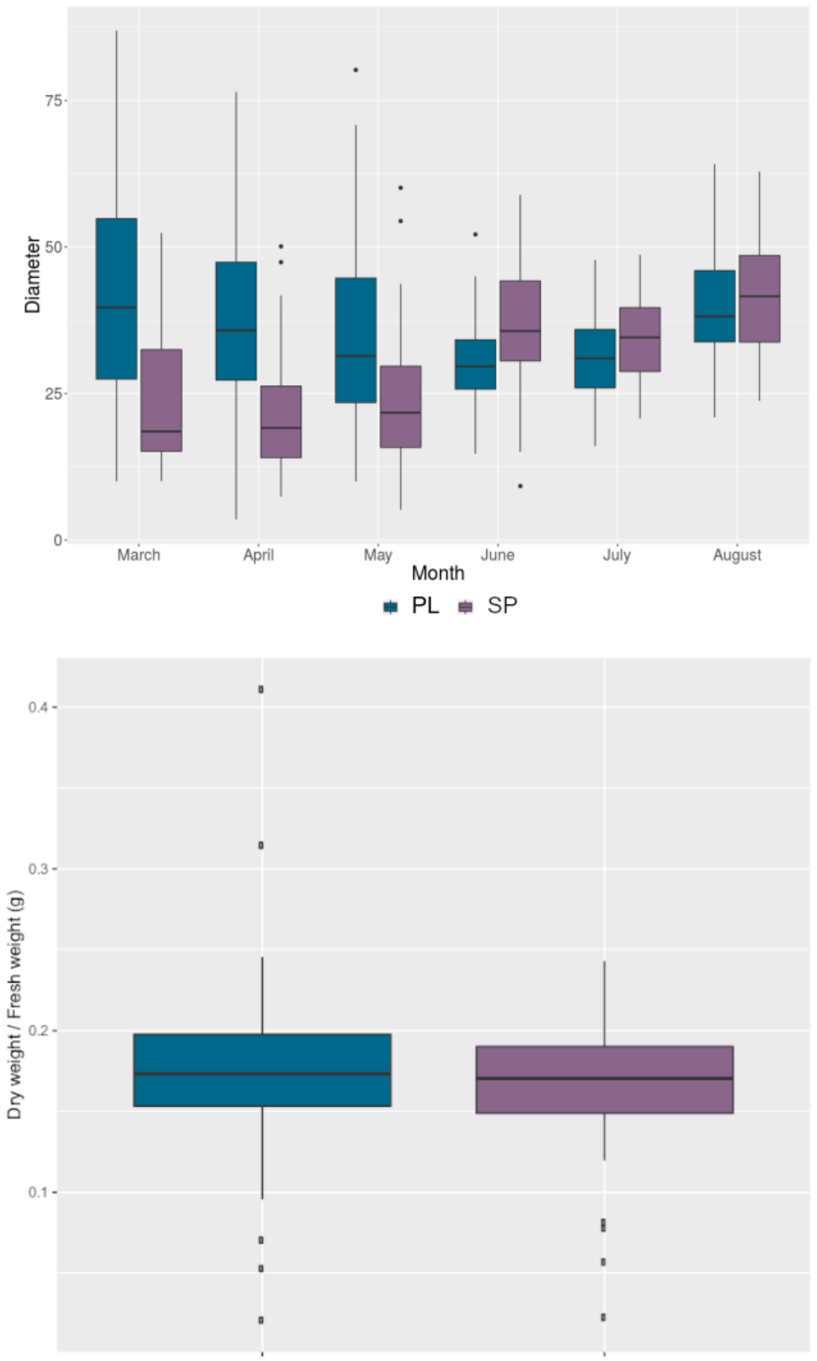


**Fig S11:** SP and PL show similar growth rate in a common garden experiment performed in the range core of the species. Six replicates of 10 genotypes per population were grown for one year in common garden setting in Cologne, which has climate representative of the core range of the species distribution. Top figure shows the diameter size (mm) per population for each month of the growing season in Cologne. Population and Month had a significant interaction ($p$ <2.23-16). The overall population effect was significant ($p$ = 0.01403), even though SP and PL did not differ at the end of the growing season (August $p$ =0.265). Bottom figure, shows the biomass of the plants at the end of the experiment as the dry to fresh weight ratio. The populations did not differ significantly ($p$ = 0.2873).

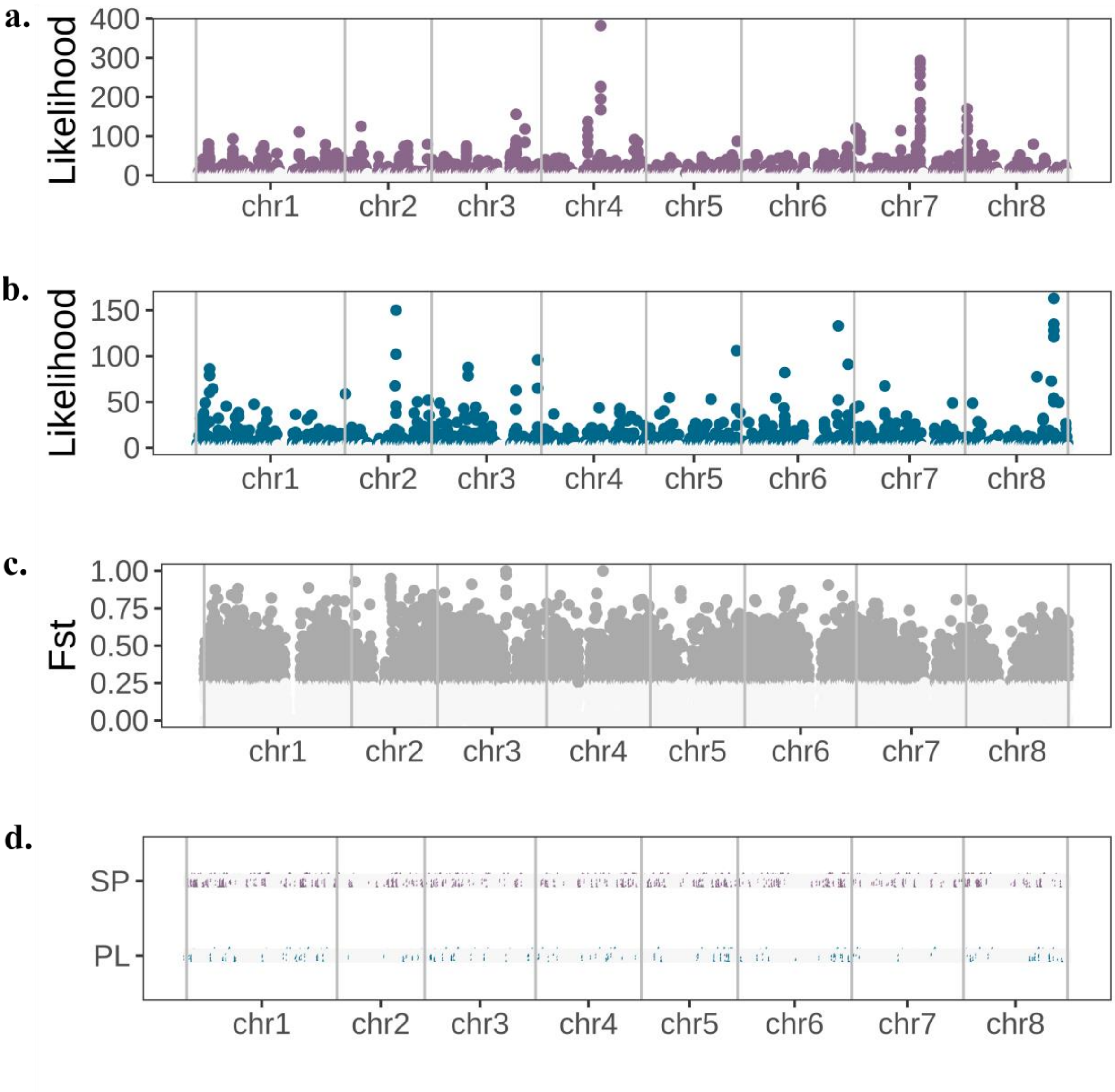


**Fig S12:** The genomic distribution of selective sweep signatures in SP and PL. The Composite Likelihood Ratio (CLR) of a sweep along the **a.** SP and **b.** PL genomes. The outlier loci are colored in dark purple for SP and blue for PL. **c.** $F_{ST}$ outliers (5$^{th}$ percentile) between SP and PL. d. The position of local sweep areas in SP and PL, combining high CLR and high $F_{ST}$. The exact location and size of the areas is provided in Table S12.

**Supplementary Code**

```
######################################################################
# estimate distribution of fitness effects and burden for PL and SP
######################################################################

# python 2.7.16

# move to correct folder
import os
path="PATH"
os.chdir(path)

# import functions
import numpy as np
import sys
import pickle
import scipy
import matplotlib.pyplot as plt
plt.rcParams["figure.facecolor"] = "w"
plt.rcParams["font.size"] = 18
plt.style.use('bmh')
np.seterr(all='ignore')

# import dadi functions
import dadi
from dadi import logging
logging.basicConfig()
```

In [3]:

```
# import fitdadi function that we are using
# the following code has been taken directly from https://bitbucket.org/gut
enkunstlab/fitdadi/src/master/Selection.py
# we have added some modifications that are indicated by ### at the end of
the line


import os
import sys
import operator
import numpy
from tqdm import tqdm_notebook
from numpy import logical_and, logical_not
from scipy.special import gammaln
import scipy.stats.distributions
import scipy.integrate
import scipy.optimize
from dadi import Numerics, Inference, Misc
from dadi.Spectrum_mod import Spectrum
```

```
class spectra:
    def __init__(self, params, ns, demo_sel_func, pts=500, pts_l=None, ###
added pts=500
                 Npts=500, n=20170., int_breaks=None, ### added n=20170
                 int_bounds=(1e-4, 1000.), mp=False, echo=False, cpus=None)
:
        """
        params: optimized demographic parameters, don't include gamma
        here
        demo_sel_func: dadi demographic function with selection. gamma
        must be the last argument.
        ns: sample sizes
        Npts: number of grid points over which to integrate
        steps: use this to set break points for spacing out the
        intervals
        mp: True if you want to use multiple cores (utilizes
        multiprocessing) if using the mp option you must also specify
        # of cpus, otherwise this will just use nthreads-1 on your
        machine
        """

        self.ns = ns
        self.spectra = []

        #create a vector of gammas that are log-spaced over sequential
        #intervals or log-spaced over a single interval.
        if not (int_breaks is None):
            numbreaks = len(int_breaks)
            stepint = Npts/(numbreaks-1)
            self.gammas = []
            for i in reversed(range(0,numbreaks-1)):
                self.gammas = numpy.append(
                    self.gammas, -numpy.logspace(numpy.log10(int_breaks[i+1
]),
                                                 numpy.log10(int_breaks[i])
,
                                                 stepint))
        else:
            self.gammas = -numpy.logspace(numpy.log10(int_bounds[1]),
                                          numpy.log10(int_bounds[0]), Npts)
        if pts_l == None: ### added if statement here if pts_l not given
            self.pts = pts
            self.pts_l = [self.pts,self.pts+(self.pts/5), \
                          self.pts+(self.pts/5)*2]
        else:
            self.pts_l = pts_l
        func_ex = Numerics.make_extrap_func(demo_sel_func)
```

```python
        self.params = tuple(params)
        if not mp: #for running with a single thread
            for ii,gamma in enumerate(self.gammas):
                self.spectra.append(func_ex(tuple(params)+(gamma,), self.ns
,
                                            self.pts_l))
                if echo:
                    print('{0}: {1}'.format(ii, gamma))
        else: #for running with with multiple cores
            import multiprocessing
            if cpus is None:
                cpus = multiprocessing.cpu_count() - 1
            print(cpus)
            def worker_sfs(in_queue, outlist, popn_func_ex, params, ns,
                           pts_l):
                """
                Worker function -- used to generate SFSes for
                single values of gamma.
                """
                while True:
                    item = in_queue.get()
                    if item == None:
                        return
                    ii, gamma = item
                    sfs = popn_func_ex(tuple(params)+(gamma,), ns, pts_l)
                    print('{0}: {1}'.format(ii, gamma))
                    result = (gamma, sfs)
                    outlist.append(result)
            manager = multiprocessing.Manager()
            results = manager.list()
            work = manager.Queue(cpus)
            pool = []
            for i in range(cpus):
                p = multiprocessing.Process(target=worker_sfs,
                                            args=(work, results, func_ex,
                                            params, self.ns, self.pts_l))
                p.start()
                pool.append(p)
            for ii,gamma in enumerate(tqdm_notebook(self.gammas)): ### adde
d tqdm_notebook()
                work.put((ii, gamma))
            for jj in tqdm_notebook(range(cpus)): ### added tqdm_notebook()
                work.put(None)
            for p in pool:
                p.join()
            reslist = []
            for line in results:
                reslist.append(line)
            reslist.sort(key = operator.itemgetter(0))
```

```
            for gamma, sfs in reslist:
                self.spectra.append(sfs)

        #self.neu_spec = demo_sel_func(params+(0,), self.ns, self.pts)
        self.neu_spec = func_ex(tuple(params)+(0,), self.ns, self.pts_l)
        self.extrap_x = self.spectra[0].extrap_x
        self.spectra = numpy.array(self.spectra)

    def integrate(self, params, sel_dist, theta): ### this is called integr
ate_old in Selection.py
        """
        integration without re-normalizing the DFE. This assumes the
        portion of the DFE that is not integrated is not seen in your
        sample.
        """
        #need to include tuple() here to make this function play nice
        #with numpy arrays
        sel_args = (self.gammas,) + tuple(params)
        #compute weights for each fs
        weights = sel_dist(*sel_args)

        #compute weight for the effectively neutral portion. not using
        #CDF function because I want this to be able to compute weight
        #for arbitrary mass functions
        weight_neu, err_neu = scipy.integrate.quad(sel_dist, self.gammas[-1
],
                                                   0, args=tuple(params))

        #function's adaptable for demographic models from 1-3 populations
        pops = len(self.neu_spec.shape)
        if pops == 1:
            integrated = self.neu_spec*weight_neu + Numerics.trapz(
                weights[:,numpy.newaxis]*self.spectra, self.gammas, axis=0)
        elif pops == 2:
            integrated = self.neu_spec*weight_neu + Numerics.trapz(
                weights[:,numpy.newaxis,numpy.newaxis]*self.spectra,
                self.gammas, axis=0)
        elif pops == 3:
            integrated = self.neu_spec*weight_neu + Numerics.trapz(
                weights[:,numpy.newaxis,numpy.newaxis,numpy.newaxis]*self.s
pectra,
                self.gammas, axis=0)
        else:
            raise IndexError("Must have one to three populations")

        integrated_fs = Spectrum(integrated, extrap_x=self.extrap_x)

        #no normalization, allow lethal mutations to fall out
        return integrated_fs * theta
```

```python
#define a bunch of default distributions just to make everything easier
def gamma_dist(mgamma, alpha, beta):
    """
    x, shape, scale
    """
    return scipy.stats.distributions.gamma.pdf(-mgamma, alpha, scale=beta)
```

In [4]:

```python
# starting out with the initial demographic parameters
# these have been estimated using fastsimcoal2
Nanc = 2846301.0/2
Nbot = 1708210.0/2
SPN2 = 13218.0/2
PLN2 = 898319.0/2
SPN1 = 13218.0/2
PLN1 = 101220.0/2
TPLSP = 26902.0
TPLAUS = 372701.0

# Migration rates
M_SPPL = 0.415 # SP to PL scaled migration rate
M_PLSP = 0.036 # PL to SP scaled migration rate
T_ISOL = 24726

# Durations of demographic periods
TB = (TPLAUS-TPLSP)/(2*Nanc) # bottleneck duration following (SP-PL)/AUS sp
lit
TF = (TPLSP-T_ISOL)/(2*Nanc) # duration from split until migration stops
TI = T_ISOL/(2*Nanc)         # duration of exponential decline until presen
t

nuB = Nbot/Nanc # scaled population size during ancestral bottleneck
SPnuF2 = SPN2/Nanc # scaled population size of SP after split with PL
PLnuF2 = PLN2/Nanc # scaled population size of PL after split with SP
SPnuF1 = SPN1/Nanc # scaled final population size of SP
PLnuF1 = PLN1/Nanc # scaled final population size of PL

# a different number of sites are retained after all filtering steps
# ratio of all observable sites SP/PL
SP_PL_theta_ratio = 0.7402746
```

In [5]:

```python
# neutral models for demographic history
# functions estimate the neutral SFS based on the demographic parameters
# dadi functions within to incorporate population size change and split of
populations
#
# single SFS returned
```

```
def three_epoch_growth_neut(params, pts):
    nuF2, nuF1 = params
    xx = dadi.Numerics.default_grid(pts)
    phi = dadi.PhiManip.phi_1D(xx)
    phi = dadi.Integration.one_pop(phi, xx, TB, nuB)
    phi = dadi.Integration.one_pop(phi, xx, TF, params[0])
    # add decrease in population size
    nu_func = lambda t: nuF2*(nuF1/nuF2)**(t/TI)
    phi = dadi.Integration.one_pop(phi, xx, TI, nu_func)
    fs = dadi.Spectrum.from_phi(phi, (28,), (xx,))
    return fs

# SFS for both populations returned
def three_epoch_growth_neut_SP_opt(params, pts):
    nuB, PLnu, SPnuF2, SPnuF1, pi = params
    xx = dadi.Numerics.default_grid(1000)
    T1 = (TB + TF + TI)*pi
    T2 = (TB + TF + TI)*(1-pi)
    phi = dadi.PhiManip.phi_1D(xx)
    phi_2 = dadi.Integration.one_pop(phi, xx, T1, nuB)
    phi_3_PL = dadi.Integration.one_pop(phi_2, xx, T2, PLnu)
    # add decrease in population size
    nu_funcSP = lambda t: SPnuF2*(SPnuF1/SPnuF2)**(t/T2)
    phi_4_SP = dadi.Integration.one_pop(phi_2, xx, T2, nu_funcSP)
    fs_PL = dadi.Spectrum.from_phi(phi_3_PL, (28,), (xx,))
    fs_SP = dadi.Spectrum.from_phi(phi_4_SP, (28,), (xx,))
    return fs_PL, fs_SP
```

In [6]:

```
# read in SFS data
# using dadi function
# fold SFS if unfolded

data_SP_0fold = dadi.Spectrum.from_file('input_files/input_SP_0fold_unfolde
d.sfs').fold()
data_PL_0fold = dadi.Spectrum.from_file('input_files/input_PL_0fold_unfolde
d.sfs').fold()
data_SP_4fold = dadi.Spectrum.from_file('input_files/input_SP_4fold_unfolde
d.sfs').fold()
data_PL_4fold = dadi.Spectrum.from_file('input_files/input_PL_4fold_unfolde
d.sfs').fold()
```

```
/home/ksteige/anaconda2/lib/python2.7/site-packages/dadi/Numerics.py:138: F
utureWarning: Using a non-tuple sequence for multidimensional indexing is d
eprecated; use `arr[tuple(seq)]` instead of `arr[seq]`. In the future this
will be interpreted as an array index, `arr[np.array(seq)]`, which will res
ult either in an error or a different result.
  return arr[reverse_slice]
```

In [7]:

```
# use a simplified demographic model for SP and PL population for downstrea
m analyses
# optimize demographic parameters for PL and SP population based on initial
parameters estimated
# compare SFS predicted by the model to the data
#
# inferrence by maximizing the composite likelihood of the folded SFS
# using the L-BFGS-B method and basinhopping in the scipy package

def obj_funct_SP_opt(params):
    demo_params = np.concatenate([10**np.array([params[0], params[1], param
s[2], params[3]]), [params[4]]])
    sfs_PL, sfs_SP = three_epoch_growth_neut_SP_opt(demo_params, 400)
    sfs_PL_fold = sfs_PL.fold()*10**params[5]
    sfs_SP_fold = sfs_SP.fold()*10**params[5]*SP_PL_theta_ratio
    result = -Inference.ll(sfs_PL_fold, data_PL_4fold) - Inference.ll(sfs_S
P_fold, data_SP_4fold)
    return result


start_sfs_PL, _ = three_epoch_growth_neut_SP_opt([nuB, PLnuF2, SPnuF2, SPnu
F1, (TB)/(TF+TI+TB)], 1000)
start_theta = dadi.Inference.optimal_sfs_scaling(start_sfs_PL, data_PL_4fol
d)
scipy.optimize.basinhopping(obj_funct_SP_opt, [np.log10(nuB), np.log10(PLnu
F2), np.log10(SPnuF2), np.log10(SPnuF2), 0.1, np.log10(start_theta)],
                            T=100, stepsize=5, interval=5, disp=True, niter
=10,
                            minimizer_kwargs={"method":"L-BFGS-B",
                                              "bounds":[(-1, 1), (-3, 1), (
-4, 3), (-4, 3), (0.001, 0.999), (4, np.log10(5e5))],
                                              "options":{"disp":False,"maxc
or":20}})
```

```
basinhopping step 0: f 273.771
basinhopping step 1: f 273.771 trial_f 273.771 accepted 1  lowest_f 273.771
basinhopping step 2: f 273.771 trial_f 474.333 accepted 0  lowest_f 273.771
basinhopping step 3: f 273.771 trial_f 273.771 accepted 1  lowest_f 273.771
basinhopping step 4: f 273.771 trial_f 417.761 accepted 0  lowest_f 273.771
adaptive stepsize: acceptance rate 0.400000 target 0.500000 new stepsize 4.
5 old stepsize 5
basinhopping step 5: f 273.771 trial_f 273.771 accepted 1  lowest_f 273.771
basinhopping step 6: f 273.771 trial_f 9419.09 accepted 0  lowest_f 273.771
basinhopping step 7: f 273.771 trial_f 820.515 accepted 0  lowest_f 273.771
basinhopping step 8: f 273.771 trial_f 273.771 accepted 1  lowest_f 273.771
basinhopping step 9: f 273.771 trial_f 417.761 accepted 0  lowest_f 273.771
adaptive stepsize: acceptance rate 0.400000 target 0.500000 new stepsize 4.
05 old stepsize 4.5
```

```
basinhopping step 10: f 273.771 trial_f 273.771 accepted 1  lowest_f 273.77
1
```

Out[7]:

```
                        fun: 273.77143702520334
 lowest_optimization_result:       fun: 273.77143702520334
 hess_inv: <6x6 LbfgsInvHessProduct with dtype=float64>
      jac: array([ 0.00463842,  0.00600267, -0.00045475,  0.0007276 , -0.00
236469,
        0.00636646])
  message: 'CONVERGENCE: REL_REDUCTION_OF_F_<=_FACTR*EPSMCH'
     nfev: 385
      nit: 45
   status: 0
  success: True
        x: array([-0.09028819, -0.17579905, -1.63579769, -0.09814862,  0.76
539362,
        4.38111948])
                    message: ['requested number of basinhopping iterations
completed successfully']
      minimization_failures: 0
                       nfev: 7679
                        nit: 10
                          x: array([-0.09028819, -0.17579905, -1.63579769,
-0.09814862,  0.76539362,
        4.38111948])
```

In [8]:

```
# demograhic parameters estimated by the optimization
# we optimized parameters for the population size, timing of the split and
theta

params = [-0.09028819, -0.17579905, -1.63579769, -0.09814862,  0.76539362,
4.38111948]
nuB_opt, PLnu_opt, SPnuF2_opt, SPnuF1_opt, _, theta_PL_opt = 10**np.array(p
arams)
opt = params[4]
nuB_opt, PLnu_opt, SPnuF2_opt, SPnuF1_opt, opt, theta_PL_opt
```

Out[8]:

```
(0.8122913153879385,
 0.667115375080474,
 0.02313142084068101,
 0.7977216520696031,
 0.76539362,
 24050.24362147892)
```

In [9]:

```
# get population size N over time T for the simplified model
# estimates for PL and SP

T1 = (TB+TF+TI)*opt
```

```python
T2 = (TB+TF+TI)*(1-opt)
T_set = np.arange(1+round(Nanc*TB)+round(Nanc*TF)+round(Nanc*TI))
N_set_SP = [Nanc] + [Nanc*nuB_opt]*int(round(Nanc*T1+1)) + [Nanc*SPnuF2_opt
*(SPnuF1_opt/SPnuF2_opt)**(t/round(Nanc*T2)) for t in np.arange(1,1+round(N
anc*T2))]
N_set_PL = [Nanc] + [Nanc*nuB_opt]*int(round(Nanc*T1+1)) + [Nanc*PLnu_opt]*
int(round(Nanc*T2))
#N_set_SP = [Nanc] + [Nanc*nuB_opt]*142632 + [Nanc*SPnuF2_opt*(SPnuF1_opt/S
PnuF2_opt)**(t/43719.0) for t in np.arange(1,1+(43719.0))]
#N_set_PL = [Nanc] + [Nanc*nuB_opt]*142632 + [Nanc*PLnu_opt]*43719

Ne_min = np.min([np.min(N_set_SP), np.min(N_set_PL)])
Ne_max = np.max([np.max(N_set_SP), np.max(N_set_PL)])
```

In [10]:

```python
# plot new simplified demographic history for PL and SP

fig, axes = plt.subplots(1, 2, figsize=(17,6))

axes[0].plot(T_set, N_set_SP)
axes[0].set_yscale('log')
axes[0].set_xlabel("generations")
axes[0].set_ylabel(r"$N_e$")
axes[0].set_title("SP")
axes[0].set_ylim([Ne_min/2, Ne_max*2])

axes[1].plot(T_set, N_set_PL)
axes[1].set_yscale('log')
axes[1].set_xlabel("generations")
axes[1].set_ylabel(r"$N_e$")
axes[1].set_title("PL")
axes[1].set_ylim([Ne_min/2, Ne_max*2])

fig.suptitle("Re-fit Split Time")
#fig.savefig("demography_refit_split.pdf", bbox_inches="tight")
```

Out[10]:

```
<matplotlib.text.Text at 0x7f51f79936d0>
```

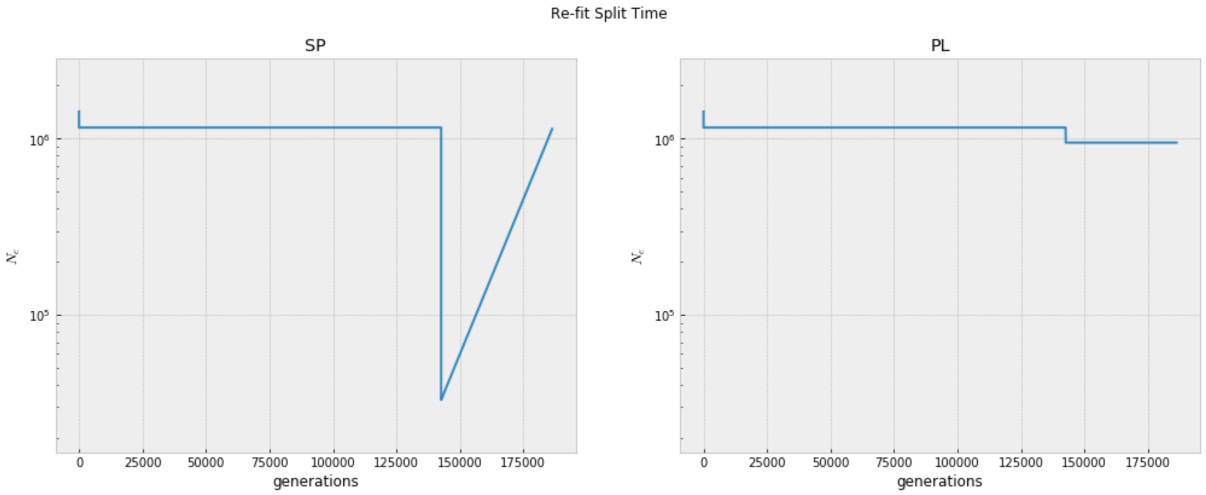


In [11]:

```
# define new demographic parameters as list to be used in further steps

demo_params_opt_PL = [nuB_opt, PLnu_opt, PLnu_opt, TB, TF, TI, opt]
demo_params_opt_SP = [nuB_opt, SPnuF2_opt, SPnuF1_opt, TB, TF, TI, opt]
demo_params_opt = [nuB_opt, PLnu_opt, PLnu_opt, SPnuF2_opt, SPnuF1_opt, TB,
TF, TI, opt]
```

In [12]:

```
# define functions to generate SFS under selection
# this needs the demographic parameters and a value of gamma
# function includes dadi functions for population size change and split of
populations
# to speed up calculations a minimum gamma value was chosen for very strong
selection
# dominance h set to 0.5 here
#
# SFS for a single population as output
def three_epoch_growth_sel_new(params, ns, pts):
    nuB, nuF2, nuF1, TB, TF, TI, pi, gamma = params
    T1 = (TB+TF+TI)*pi
    T2 = (TB+TF+TI)*(1-pi)
    xx = dadi.Numerics.default_grid(pts)
    min_gamma = -500 # minimum gamma value for strong selection
    if gamma < min_gamma:
        phi = dadi.PhiManip.phi_1D(xx, gamma=min_gamma)
    else:
        phi = dadi.PhiManip.phi_1D(xx, gamma=gamma)

    if nuB*gamma < min_gamma:
        phi = dadi.Integration.one_pop(phi, xx, T1, nuB, gamma=min_gamma)
    else:
        phi = dadi.Integration.one_pop(phi, xx, T1, nuB, gamma=gamma)

    if gamma == 0:
        nu_func = lambda t: nuF2*(nuF1/nuF2)**(t/TI)
```

```
    else:
        nu_func = lambda t: np.where(gamma*nuF2*(nuF1/nuF2)**(t/T2) < min_g
amma,
                                     np.abs(min_gamma/gamma),
                                     nuF2*(nuF1/nuF2)**(t/T2))

    phi = dadi.Integration.one_pop(phi, xx, T2, nu_func, gamma=gamma)
    fs = dadi.Spectrum.from_phi(phi, (28,), (xx,))
    return fs

# SFS for both populations as output
def three_epoch_growth_sel_both_new(params, ns, pts):
    min_gamma = -500
    nuB, PLnuF2, PLnuF1, SPnuF2, SPnuF1, TB, TF, TI, pi, gamma = params
    T1 = (TB+TF+TI)*pi
    T2 = (TB+TF+TI)*(1-pi)
    # define grid
    xx = dadi.Numerics.default_grid(pts)
    # equilibrium ancestral population
    if gamma < min_gamma:
        phi = dadi.PhiManip.phi_1D(xx, gamma=min_gamma)
    else:
        phi = dadi.PhiManip.phi_1D(xx, gamma=gamma)
    # population reduction in ancestral pop
    if nuB*gamma < min_gamma:
        phi_2 = dadi.Integration.one_pop(phi, xx, T1, nuB, gamma=min_gamma)
    else:
        phi_2 = dadi.Integration.one_pop(phi, xx, T1, nuB, gamma=gamma)
    # split between SP and PL and migration
    phi_3 = dadi.PhiManip.phi_1D_to_2D(xx, phi_2)
    # changes in population size in PL and SP, no migration
    if gamma == 0:
        nu_func1 = lambda t: PLnuF2*(PLnuF1/PLnuF2)**(t/T2)
        nu_func2 = lambda t: SPnuF2*(SPnuF1/SPnuF2)**(t/T2)
    else:
        nu_func1 = lambda t: np.where(gamma*PLnuF2*(PLnuF1/PLnuF2)**(t/T2)
< min_gamma,
                                     np.abs(min_gamma/gamma),
                                     PLnuF2*(PLnuF1/PLnuF2)**(t/T2))
        nu_func2 = lambda t: np.where(gamma*SPnuF2*(SPnuF1/SPnuF2)**(t/T2)
< min_gamma,
                                     np.abs(min_gamma/gamma),
                                     SPnuF2*(SPnuF1/SPnuF2)**(t/T2))
    phi_4 = dadi.Integration.two_pops(phi_3, xx, T2, nu1=nu_func1, nu2=nu_f
unc2, gamma1=gamma, gamma2=gamma)
    fs = dadi.Spectrum.from_phi(phi_4, (28,28), (xx,xx), pop_ids=["PL","SP"
])
    return fs
```

In [ ]:

```
# define array based on positions in the SFS
ns = np.array([28])

# run spectra function from fitdadi
# this allows to get the estimated SFS under the demographic parametrs for
a range of gammas
# this uses the previously defined function "three_epoch_growth_sel_new" an
d the simplified demographic parameters

# run for the SP population
spectra_SP_opt = spectra(demo_params_opt_SP, ns, three_epoch_growth_sel_new
, pts=1000,
                         int_bounds=(0.01, 1e5), Npts=500, echo=False, mp=Tr
ue)
# save object as the calculation takes some time
#pickle.dump(spectra_SP_opt, open("spectra_SP_break_model_new.sp","wb"))

# run for the PL population
spectra_PL_opt = spectra(demo_params_opt_PL, ns, three_epoch_growth_sel_new
, pts=1000,
                         int_bounds=(0.01, 1e5), Npts=500, echo=False, mp=Tr
ue)
#pickle.dump(spectra_PL_opt, open("spectra_PL_break_model_new.sp","wb"))

# load the object if previously saved
#spectra_SP_opt = pickle.load(open("spectra_SP_break_model_new.sp"))
#spectra_PL_opt = pickle.load(open("spectra_PL_break_model_new.sp"))
```

In [51]:

```
# calculate the ns population scaled mutation rate wich is used later
theta_PL = theta_PL_opt
# use the estimation for PL but multiply it by the SP/PL ratio that was cal
culated based on the data
theta_SP = theta_PL_opt*SP_PL_theta_ratio

theta_SP_opt_ns = theta_SP*2.76
theta_PL_opt_ns = theta_PL*2.76

# population scaled mutation rate theta
theta_SP_opt_ns = 49138.429315587644
theta_PL_opt_ns = 66378.65099732943
```

In [18]:

```
# fit the DFE by estimating the shape and scale of the gamma distribution
# optimization steps in fitdadi available, but we used scipy optimization a
s it allows more flexibility
# method for optimization in scipy was SLSQP
# for the optimization both a multinomial model (without using theta) and a
Poisson model (including theta) were used
```

```
# the optimization can be done both using a single population and a joint e
stimate (for both populations)
# the saved output from the fitdadi 'spectra' function is needed here

# the primary difference between the multinomial and the Poisson model is t
hat the multinomial model only fits the DFE for variation that is sufficien
tly weakly selected to be observed in the sample
# the reason is that the multinomial model only fits the proportions of all
eles observed at different frequencies and does not consider the decrease i
n per-site reduction in variation compared to the 4-fold sites
# strongly deleterious variation will largely be absent from our moderate s
ample size and therefore does not affect the shape of the SFS

# scipy optimization for SP under the multinomial model
SP_shape_set = np.logspace(start=-1, stop=1.0, num=250)
SP_scale_fit_set = np.ones_like(SP_shape_set)
SP_llhood_fit_set = np.ones_like(SP_shape_set)
for ii, shape in enumerate(SP_shape_set):
    def obj_funct(scale):
        sfs_SP = spectra_SP_opt.integrate([shape, scale], gamma_dist, 1).fo
ld()
        return -Inference.ll_multinom(sfs_SP, data_SP_0fold)
    fit = scipy.optimize.minimize(obj_funct, [SP_scale_fit_set[ii-1]], meth
od="SLSQP", bounds=[(np.exp(-8), np.exp(20))])
    SP_scale_fit_set[ii] = fit["x"][0]
    SP_llhood_fit_set[ii] = fit["fun"]
```

In [89]:

```
# plot the LL vs the shape parameter
fig, ax = plt.subplots(figsize=(7,4))
plt.plot(SP_shape_set, SP_llhood_fit_set-np.min(SP_llhood_fit_set))
plt.xscale("log")
plt.yscale("symlog", linthreshy=1)
plt.ylabel(r"$-LL-\min(LL)$")
plt.xlabel("shape parameter")
plt.title("SP gamma fit")
#fig.savefig("SP_only_gamma_llhood.pdf", bbox_inches="tight")
```

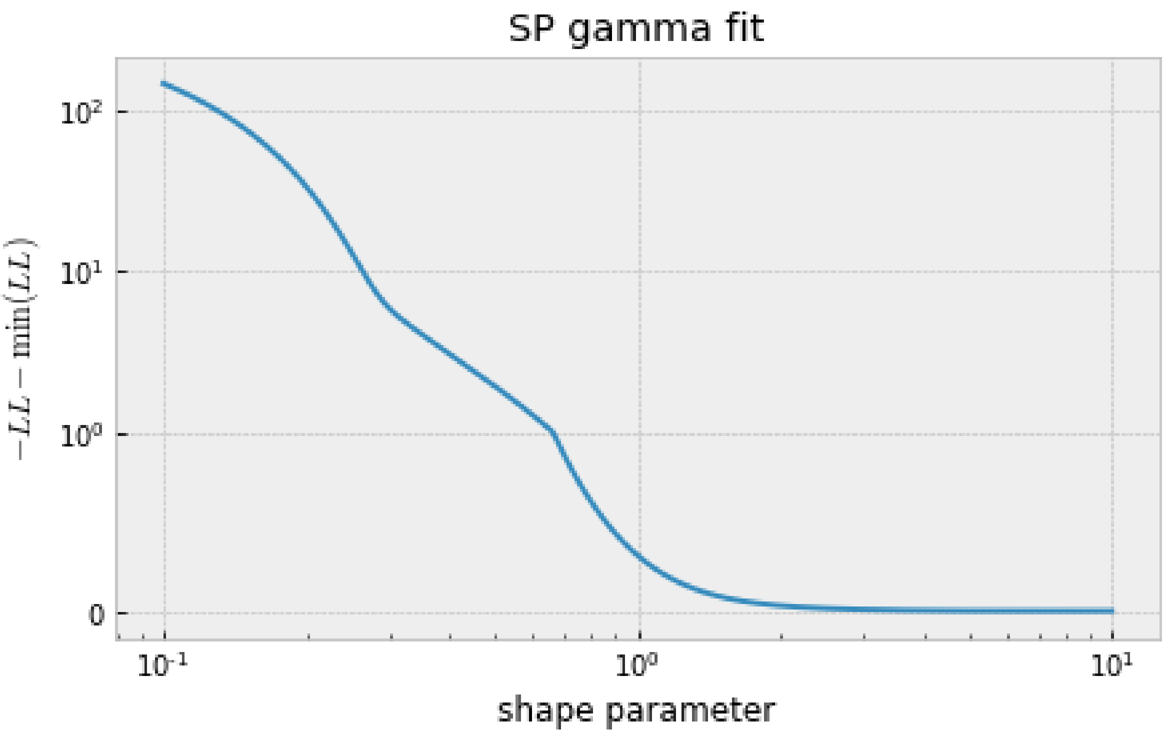


In [90]:

```
# plot the mean gamma vs the shape parameter
fig, ax = plt.subplots(figsize=(7,4))
plt.plot(SP_shape_set, SP_scale_fit_set*SP_shape_set)
plt.xscale("log")
plt.yscale("symlog", linthreshy=1)
plt.ylim([0,500])
plt.ylabel(r"mean $\gamma$")
plt.xlabel("shape parameter")
plt.title("SP gamma fit")
#fig.savefig("SP_only_gamma_mean.pdf", bbox_inches="tight")
```

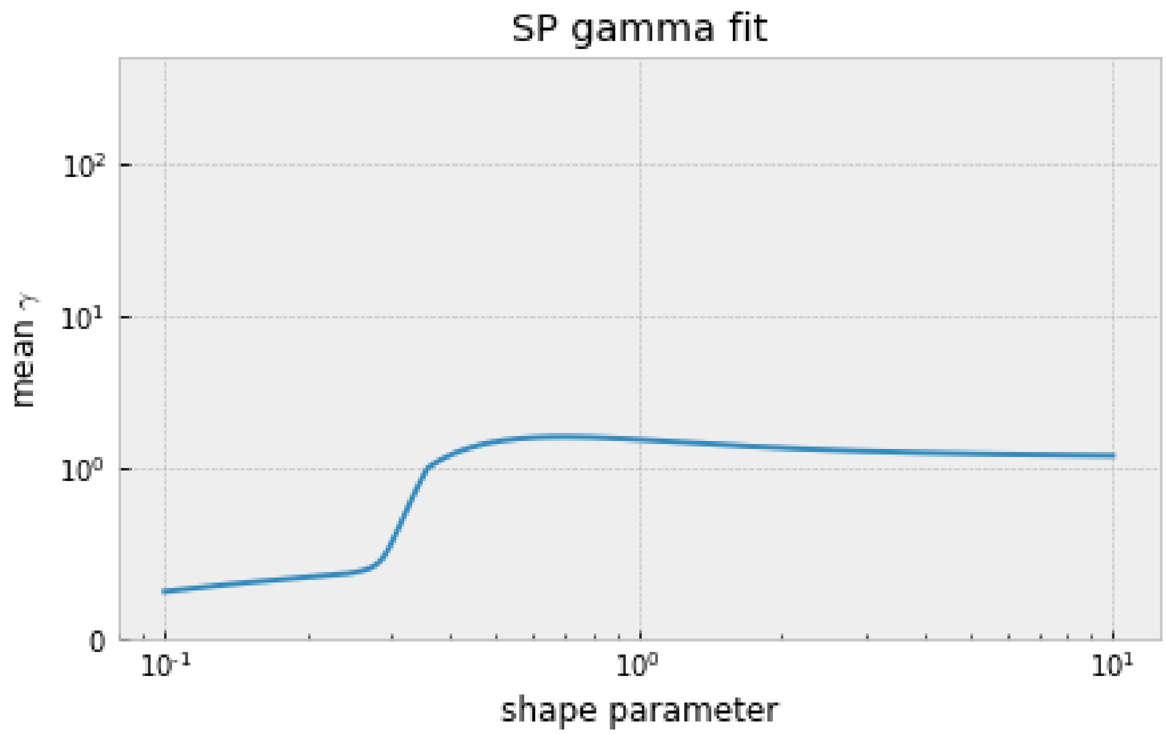


In [23]:

```
# scipy optimization for SP under the Poisson model including theta
SP_shape_set_t = np.logspace(start=-2, stop=0, num=250)
SP_scale_fit_set_t = np.ones_like(SP_shape_set_t)
SP_llhood_fit_set_t = np.ones_like(SP_shape_set_t)
for ii, shape in enumerate(SP_shape_set_t):
    def obj_funct(scale):
        sfs_SP = spectra_SP_opt.integrate([shape, np.exp(scale)], gamma_dis
t, theta_SP_opt_ns).fold()
        return -Inference.ll(sfs_SP, data_SP_0fold)
    fit = scipy.optimize.minimize(obj_funct, [np.log(100)], method="SLSQP",
bounds=[(-8, 20)])
    SP_scale_fit_set_t[ii] = np.exp(fit["x"][0])
    SP_llhood_fit_set_t[ii] = fit["fun"]
```

In [91]:

```
# plot the LL vs the shape parameter
fig, ax = plt.subplots(figsize=(7,4))
plt.plot(SP_shape_set_t, SP_llhood_fit_set_t-np.min(SP_llhood_fit_set_t), "
-")
plt.xscale("log")
plt.yscale("symlog", linthreshy=10)
plt.ylabel(r"$-LL-\min(LL)$")
plt.xlabel("shape parameter")
plt.title(r"SP gamma fit $\theta$")
#fig.savefig("SP_only_gamma_llhood_theta.pdf", bbox_inches="tight")
```

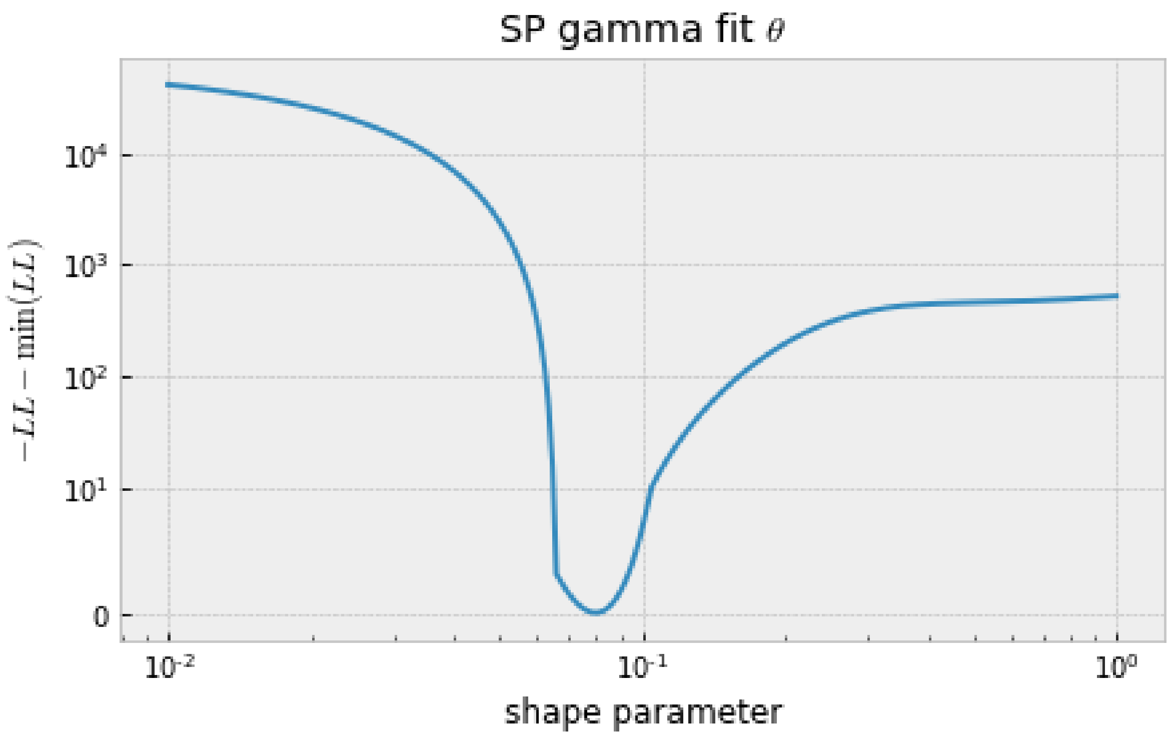


In [92]:

```
# plot the scale vs the shape parameter
fig, ax = plt.subplots(figsize=(7,4))
plt.plot(SP_shape_set_t, SP_scale_fit_set_t)
plt.xscale("log")
plt.yscale("symlog", linthreshy=2e4)
#plt.ylim([0,500])
plt.ylabel("scale parameter")
plt.xlabel("shape parameter")
plt.title(r"SP gamma fit $\theta$")
#fig.savefig("SP_only_gamma_scale_theta.pdf", bbox_inches="tight")
```

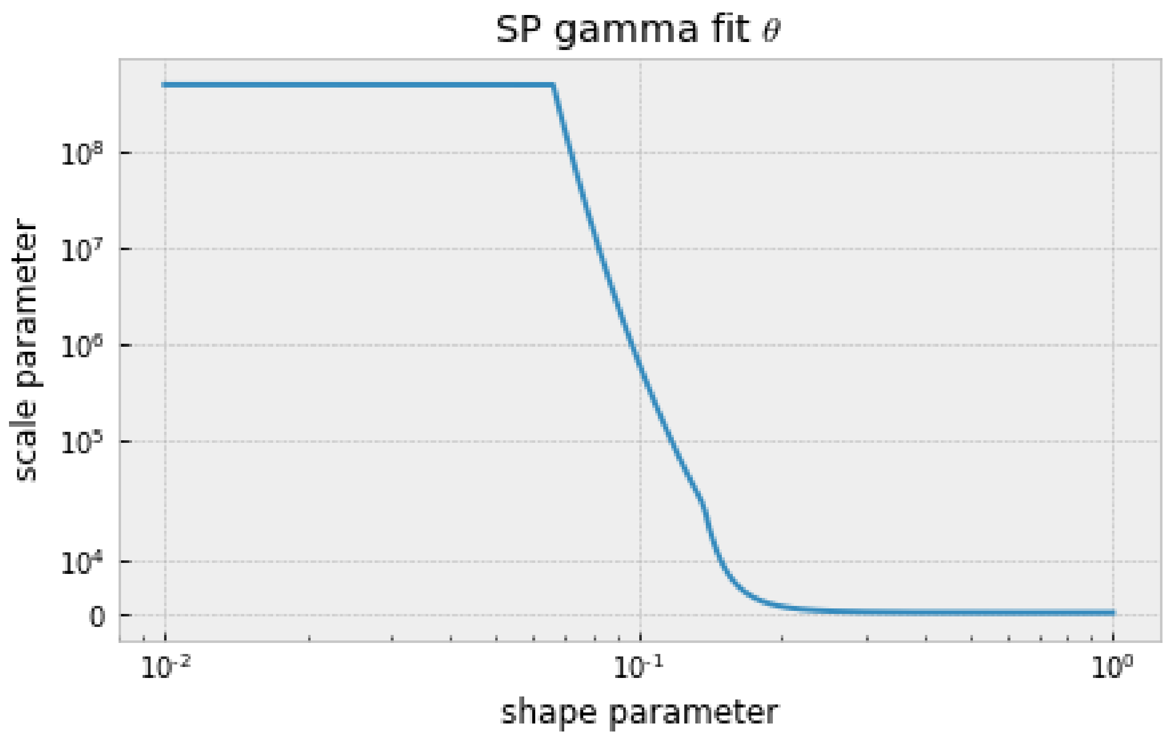


In [26]:

```
# scipy optimization for PL under the multinomial model

PL_shape_set = np.logspace(start=-1, stop=1, num=250)
PL_scale_fit_set = np.ones_like(PL_shape_set)
PL_llhood_fit_set = np.ones_like(PL_shape_set)
for ii, shape in enumerate(PL_shape_set):
    def obj_funct(scale):
        sfs_PL = spectra_PL_opt.integrate([shape, scale], gamma_dist, 1).fo
ld()
        return -Inference.ll_multinom(sfs_PL, data_PL_0fold)
    fit = scipy.optimize.minimize(obj_funct, [PL_scale_fit_set[ii-1]], meth
od="SLSQP", bounds=[(np.exp(-8), np.exp(20))])
    PL_scale_fit_set[ii] = fit["x"][0]
    PL_llhood_fit_set[ii] = fit["fun"]
```

In [93]:

```
# plot the LL vs the shape parameter
fig, ax = plt.subplots(figsize=(7,4))
plt.plot(PL_shape_set, PL_llhood_fit_set-np.min(PL_llhood_fit_set))
plt.xscale("log")
plt.yscale("symlog", linthreshy=10)
plt.ylabel(r"$-LL-\min(LL)$")
plt.xlabel("shape parameter")
plt.title("PL gamma fit")
#fig.savefig("PL_only_gamma_llhood.pdf", bbox_inches="tight")
```

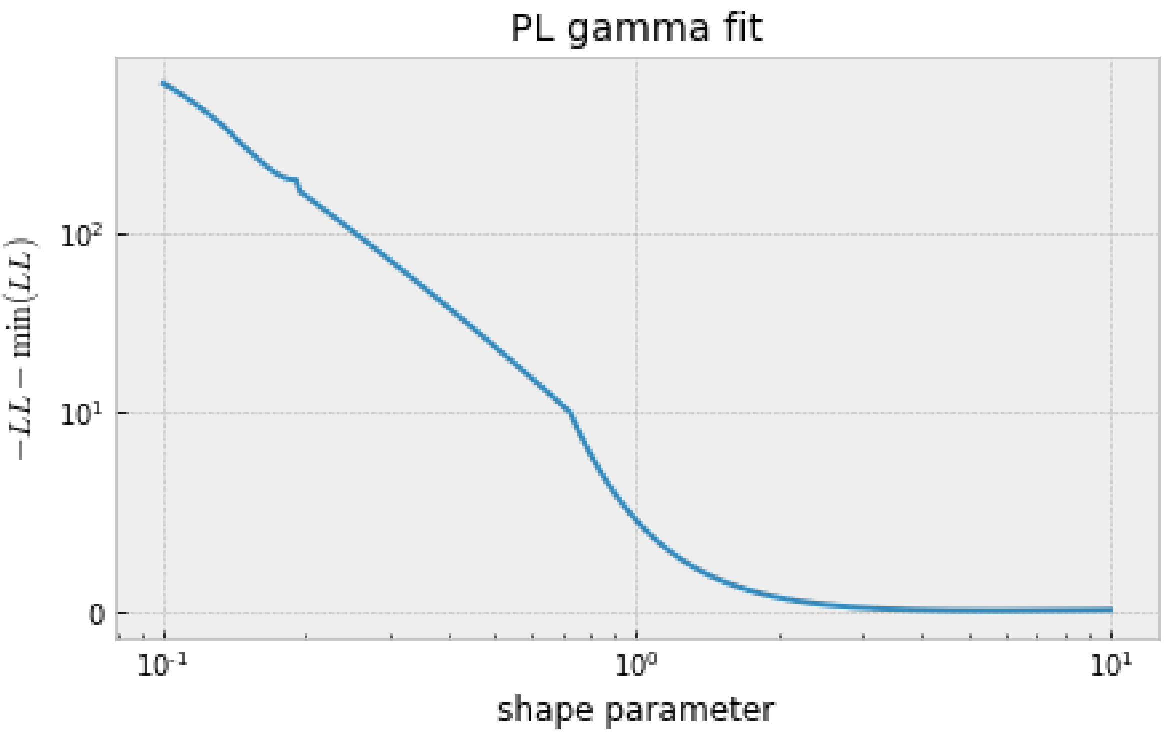


In [95]:

```
# plot the mean gamma vs the shape parameter
fig, ax = plt.subplots(figsize=(7,4))
plt.plot(PL_shape_set, PL_scale_fit_set*PL_shape_set)
plt.xscale("log")
plt.yscale("symlog", linthreshy=1)
plt.ylim([0,500])
plt.ylabel(r"mean $\gamma$")
plt.xlabel("shape parameter")
plt.title("PL gamma fit")
#fig.savefig("PL_only_gamma_mean.pdf", bbox_inches="tight")
```

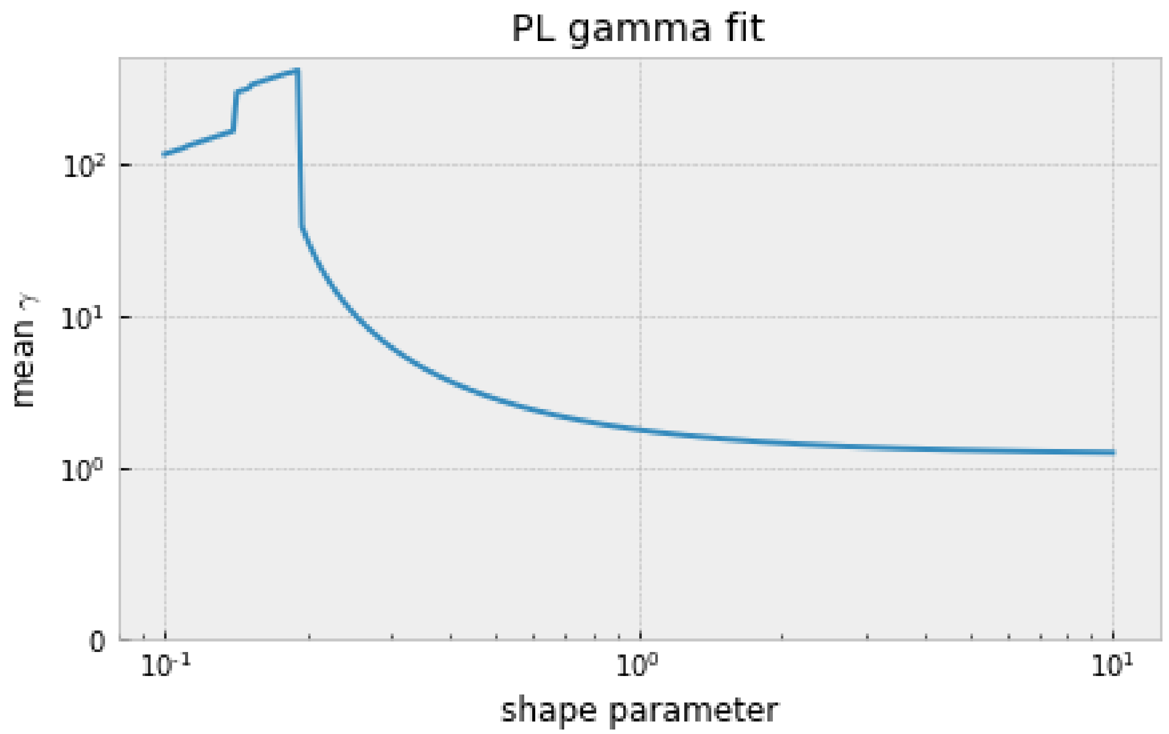


In [29]:

```
# scipy optimization for PL under the Poisson model including theta

PL_shape_set_t = np.logspace(start=-2, stop=0, num=250)
PL_scale_fit_set_t = np.ones_like(PL_shape_set_t)
PL_llhood_fit_set_t = np.ones_like(PL_shape_set_t)
for ii, shape in enumerate(PL_shape_set_t):
    def obj_funct(scale):
        sfs_PL = spectra_PL_opt.integrate([shape, np.exp(scale)], gamma_dis
t, theta_PL_opt_ns).fold()
        return -Inference.ll(sfs_PL, data_PL_0fold)
    fit = scipy.optimize.minimize(obj_funct, [np.log(100)], method="L-BFGS-
B", bounds=[(-8, 20)])
    PL_scale_fit_set_t[ii] = np.exp(fit["x"][0])
    PL_llhood_fit_set_t[ii] = fit["fun"]
```

In [96]:

```
# plot the LL vs the shape parameter
fig, ax = plt.subplots(figsize=(7,4))
plt.plot(PL_shape_set_t, PL_llhood_fit_set_t-np.min(PL_llhood_fit_set_t))
plt.xscale("log")
plt.yscale("symlog", linthreshy=10)
plt.ylabel(r"$-LL-\min(LL)$")
plt.xlabel("shape parameter")
plt.title(r"PL gamma fit $\theta$")
#fig.savefig("PL_only_gamma_llhood_theta.pdf", bbox_inches="tight")
```

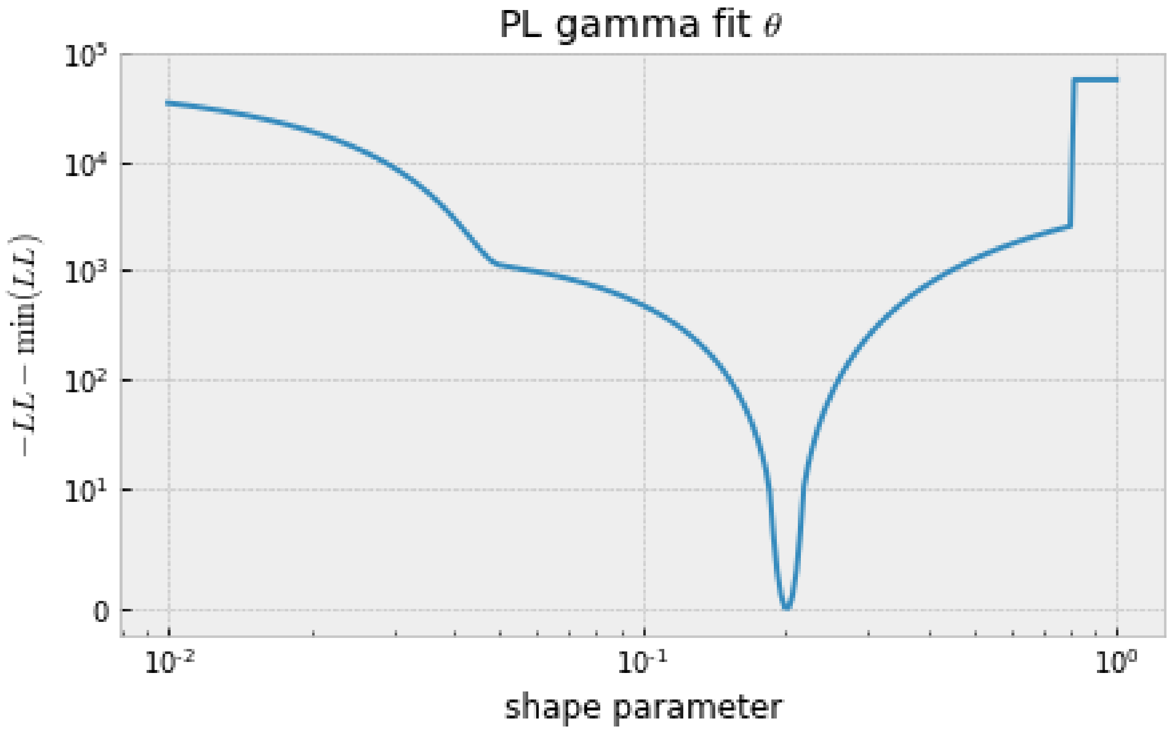


In [97]:

```
# plot the scale vs the shape parameter
fig, ax = plt.subplots(figsize=(7,4))
plt.plot(PL_shape_set_t, PL_scale_fit_set_t)
plt.xscale("log")
plt.yscale("symlog", linthreshy=1e3)
#plt.ylim([0,500])
plt.ylabel("scale parameter")
plt.xlabel("shape parameter")
plt.title(r"PL gamma fit $\theta$")
#fig.savefig("PL_only_gamma_scale_theta.pdf", bbox_inches="tight")
```

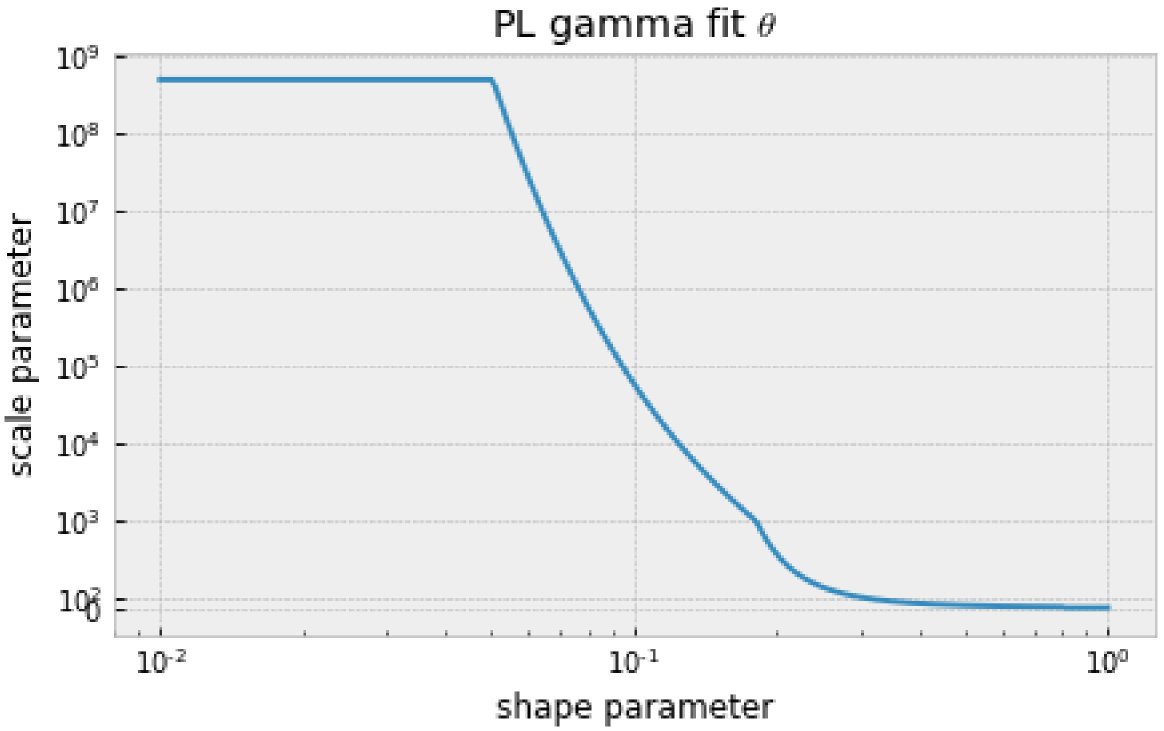


In [32]:

```
# joint scipy optimization for SP and PL under the multinomial model
shape_set = np.logspace(start=-1, stop=1.0, num=250)
scale_fit_set = np.ones_like(shape_set)
llhood_fit_set = np.ones_like(shape_set)
for ii, shape in enumerate(shape_set):
    def obj_funct(scale):
        sfs_SP = spectra_SP_opt.integrate([shape, scale], gamma_dist, 1).fo
ld()
        sfs_PL = spectra_PL_opt.integrate([shape, scale], gamma_dist, 1).fo
ld()
        return -Inference.ll_multinom(sfs_SP, data_SP_0fold) - Inference.ll
_multinom(sfs_PL, data_PL_0fold)
    fit = scipy.optimize.minimize(obj_funct, [scale_fit_set[ii-1]], method=
"SLSQP", bounds=[(np.exp(-8), np.exp(20))])
    scale_fit_set[ii] = fit["x"][0]
    llhood_fit_set[ii] = fit["fun"]
```

In [98]:

```
# plot the LL vs the shape parameter
fig, ax = plt.subplots(figsize=(7,4))
plt.plot(shape_set, llhood_fit_set-np.min(llhood_fit_set))
plt.xscale("log")
plt.yscale("symlog", linthreshy=10)
plt.ylabel(r"$-LL-\min(LL)$")
plt.xlabel("shape parameter")
plt.title("Both gamma fit")
#fig.savefig("both_gamma_llhood.pdf", bbox_inches="tight")
```

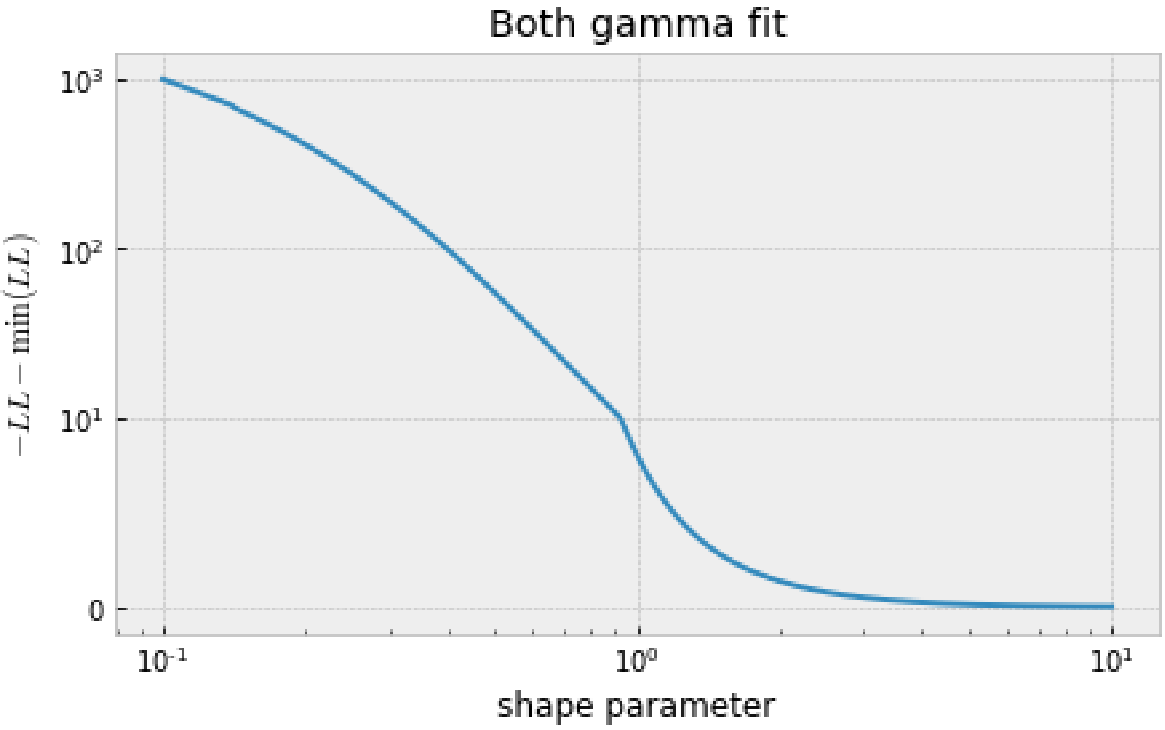


In [99]:

```
# plot the mean gamma vs the shape parameter
fig, ax = plt.subplots(figsize=(7,4))
plt.plot(shape_set, scale_fit_set*shape_set)
plt.xscale("log")
plt.yscale("symlog", linthreshy=1)
plt.ylim([0,500])
plt.ylabel(r"mean $\gamma$")
plt.xlabel("shape parameter")
plt.title("Both gamma fit")
#fig.savefig("both_gamma_mean.pdf", bbox_inches="tight")
```

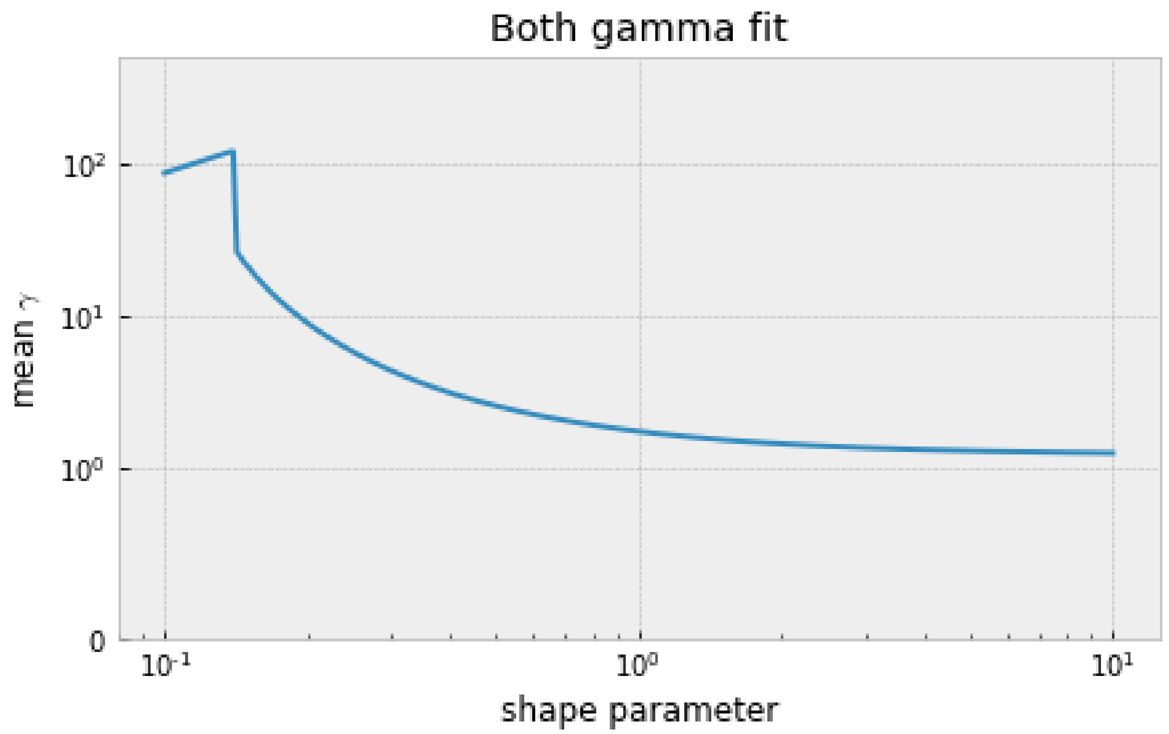


In [35]:

```
# joint scipy optimization for SP and PL under the Poisson model including
theta

shape_set_t = np.logspace(start=-2, stop=0, num=250)
scale_fit_set_t = np.ones_like(shape_set_t)
llhood_fit_set_t = np.zeros_like(shape_set_t)
for ii, shape in enumerate(shape_set_t):
    def obj_funct(scale):
        sfs_SP = spectra_SP_opt.integrate([shape, np.exp(scale)], gamma_dis
t, theta_SP_opt_ns).fold()
        sfs_PL = spectra_PL_opt.integrate([shape, np.exp(scale)], gamma_dis
t, theta_PL_opt_ns).fold()
        return -Inference.ll(sfs_SP, data_SP_0fold) - Inference.ll(sfs_PL,
data_PL_0fold)
    fit = scipy.optimize.minimize(obj_funct, [np.log(100)], method="SLSQP",
bounds=[(-8, 20)])
    scale_fit_set_t[ii] = np.exp(fit["x"][0])
    llhood_fit_set_t[ii] = fit["fun"]
```

In [100]:

```
# plot the LL vs the shape parameter
fig, ax = plt.subplots(figsize=(7,4))
plt.plot(shape_set_t, llhood_fit_set_t-np.min(llhood_fit_set_t))
plt.xscale("log")
plt.yscale("symlog", linthreshy=10)
plt.ylabel(r"$-LL-\min(LL)$")
plt.xlabel("shape parameter")
plt.title(r"Both gamma fit $\theta$")
#fig.savefig("both_gamma_llhood_theta.pdf", bbox_inches="tight")
```

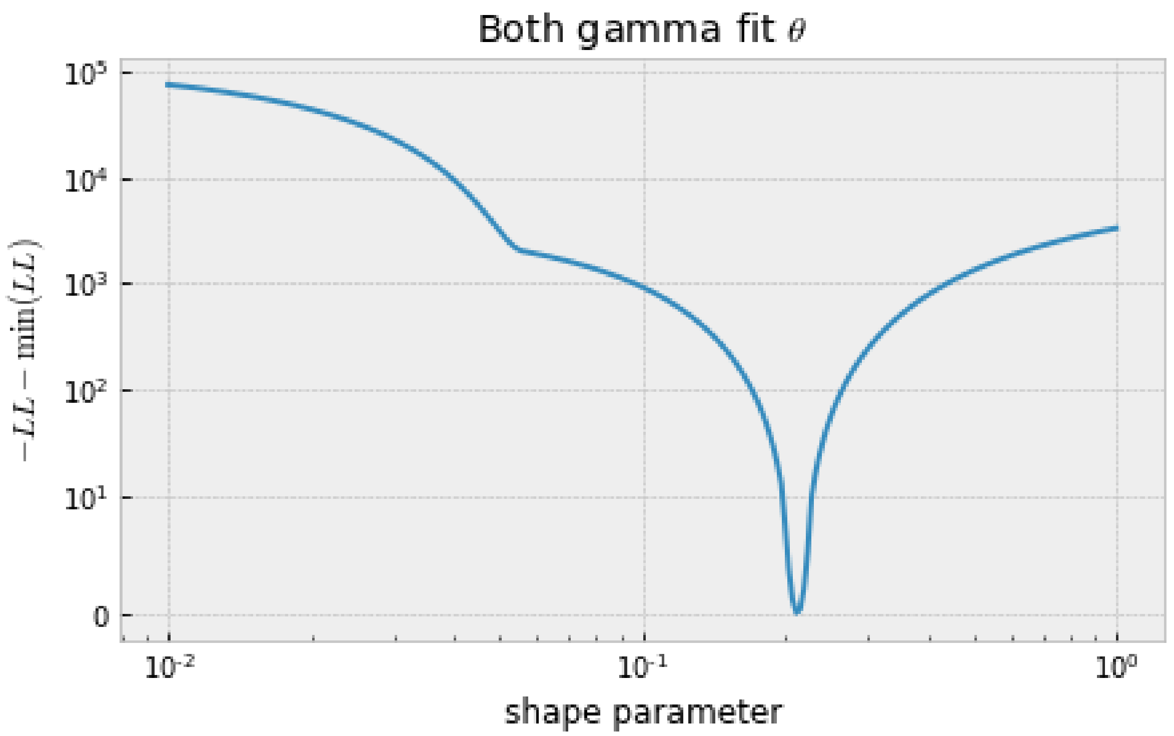


In [101]:

```
# plot the scale vs the shape parameter
fig, ax = plt.subplots(figsize=(7,4))
plt.plot(shape_set_t, scale_fit_set_t)
plt.xscale("log")
plt.yscale("symlog", linthreshy=10)
#plt.ylim([0,500])
plt.ylabel(r"scale parameter")
plt.xlabel("shape parameter")
plt.title(r"Both gamma fit $\theta$")
#fig.savefig("both_gamma_scale_theta.pdf", bbox_inches="tight")
```

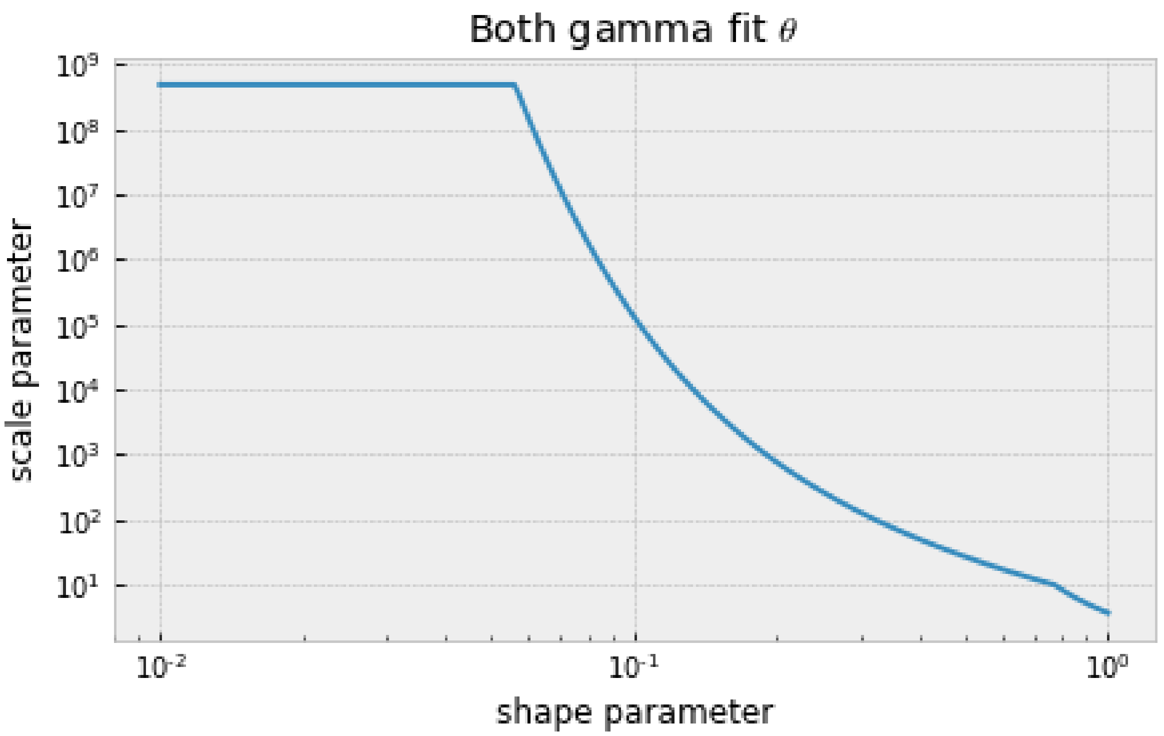


In [38]:

```
# multinomial likelihood converges on a point mass for a single s value at
~ 1.2
# calculated by shape * scale parameter
# this is true for the joint and single population estimations
np.prod([shape_set[246], scale_fit_set[246]])
```

Out[38]:

```
1.2577171629039776
```

In [39]:

```
np.prod([PL_shape_set[246], PL_scale_fit_set[246]])
```

Out[39]:

```
1.2662500256216924
```

In [40]:

```
np.prod([SP_shape_set[246], SP_scale_fit_set[246]])
```

Out[40]:

```
1.2096378596471853
```

In [127]:

```
# visualising the observed SFS and expected SFS for the multinomial model
# the expected SFS can be extracted from the object calculated using the fi
tdadi 'spectra' function

fig, ax = plt.subplots(figsize=(7,7))
fit_SP_both = spectra_SP_opt.integrate([shape_set[246], scale_fit_set[246]]
, gamma_dist, 1)
fit_PL_both = spectra_PL_opt.integrate([shape_set[246], scale_fit_set[246]]
, gamma_dist, 1)
```

```
fit_SP_only = spectra_SP_opt.integrate([SP_shape_set[246], SP_scale_fit_set
[246]], gamma_dist, 1)
fit_PL_only = spectra_PL_opt.integrate([PL_shape_set[246], PL_scale_fit_set
[246]], gamma_dist, 1)

plt.plot(fit_SP_both.fold()/np.sum(fit_SP_both.fold()),
         color="red", label="SP gamma both")
plt.plot(fit_PL_both.fold()/np.sum(fit_PL_both.fold()),
         color="blue", label="PL gamma both")
plt.plot(fit_SP_only.fold()/np.sum(fit_SP_only.fold()),
         "o", color="red", label="SP gamma only")
plt.plot(fit_PL_only.fold()/np.sum(fit_PL_only.fold()),
         "o", color="blue", label="PL gamma only")
plt.plot(data_SP_0fold/np.sum(data_SP_0fold), "--",
         color="red", label="SP (0-fold)")
plt.plot(data_PL_0fold/np.sum(data_PL_0fold), "--",
         color="blue", label="PL (0-fold)")

plt.yscale("log")
plt.ylabel("proportion of sites")
plt.xlabel("sample count (folded)")
plt.legend()
#fig.savefig("gamma_fits.pdf", bbox_inches="tight")
```

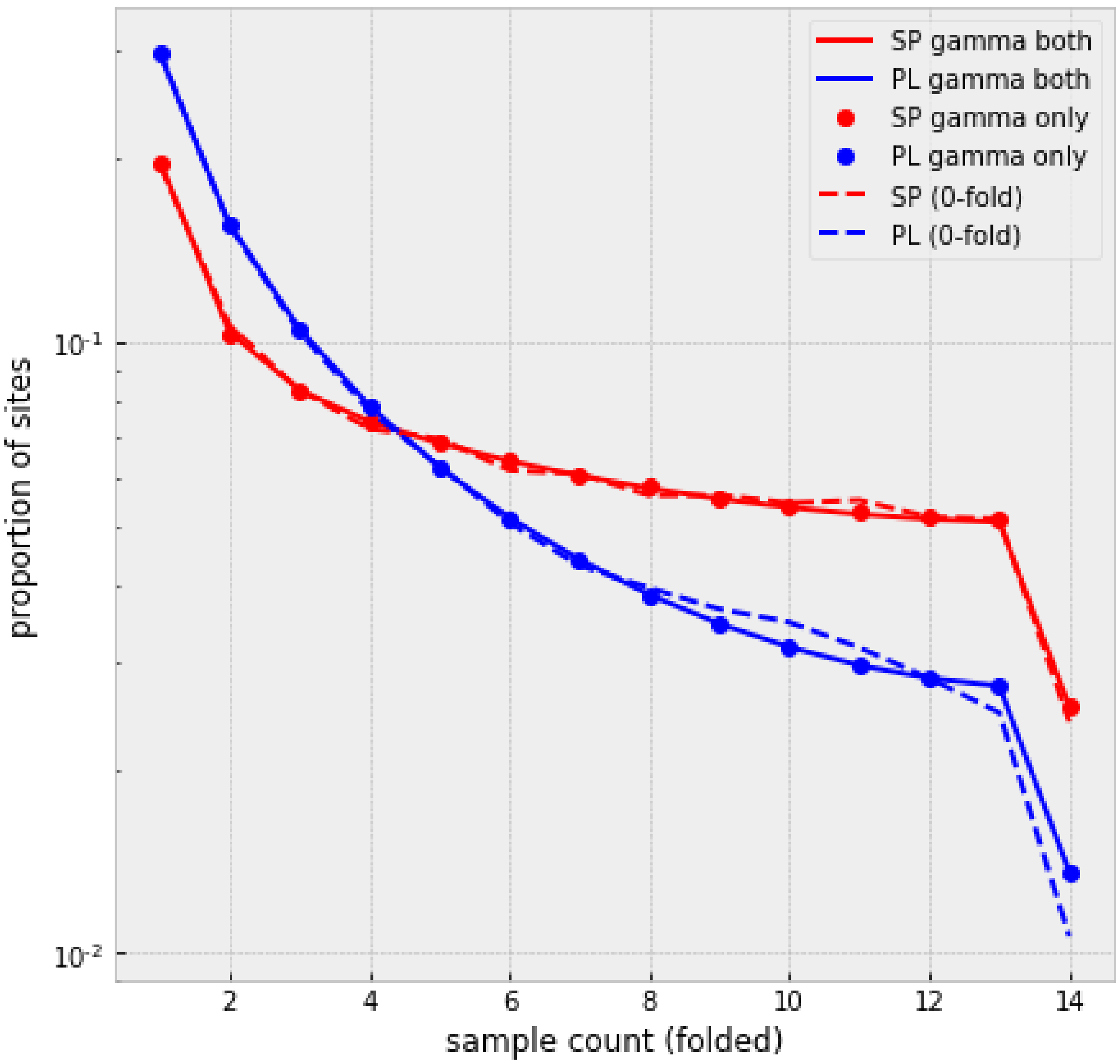


In [129]:

```
# visualising the observed SFS and expected SFS for the Poisson model inclu
ding theta

fig, ax = plt.subplots(figsize=(8,8))
min_both = np.argmin(llhood_fit_set_t)
fit_SP_both = spectra_SP_opt.integrate([shape_set_t[min_both], scale_fit_se
t_t[min_both]], gamma_dist, theta_SP_opt_ns)
fit_PL_both = spectra_PL_opt.integrate([shape_set_t[min_both], scale_fit_se
t_t[min_both]], gamma_dist, theta_PL_opt_ns)

min_PL = np.argmin(PL_llhood_fit_set_t)
min_SP = np.argmin(SP_llhood_fit_set_t)
fit_SP_only = spectra_SP_opt.integrate([SP_shape_set_t[min_SP], SP_scale_fi
t_set_t[min_SP]], gamma_dist, theta_SP_opt_ns)
fit_PL_only = spectra_PL_opt.integrate([PL_shape_set_t[min_PL], PL_scale_fi
t_set_t[min_PL]], gamma_dist, theta_PL_opt_ns)

plt.plot(fit_SP_both.fold(), color="red", label="SP gamma both")
```

```
plt.plot(fit_PL_both.fold(), color="blue", label="PL gamma both")
plt.plot(fit_SP_only.fold(),  "o", color="red", label="SP gamma only")
plt.plot(fit_PL_only.fold(), "o", color="blue", label="PL gamma only")
plt.plot(data_SP_0fold, "--", color="red", label="SP (0-fold)")
plt.plot(data_PL_0fold, "--", color="blue", label="PL (0-fold)")

plt.yscale("log")
plt.ylabel("number of sites")
plt.xlabel("sample count (folded)")
plt.legend()
#fig.savefig("gamma_fits_theta.pdf", bbox_inches="tight")
```

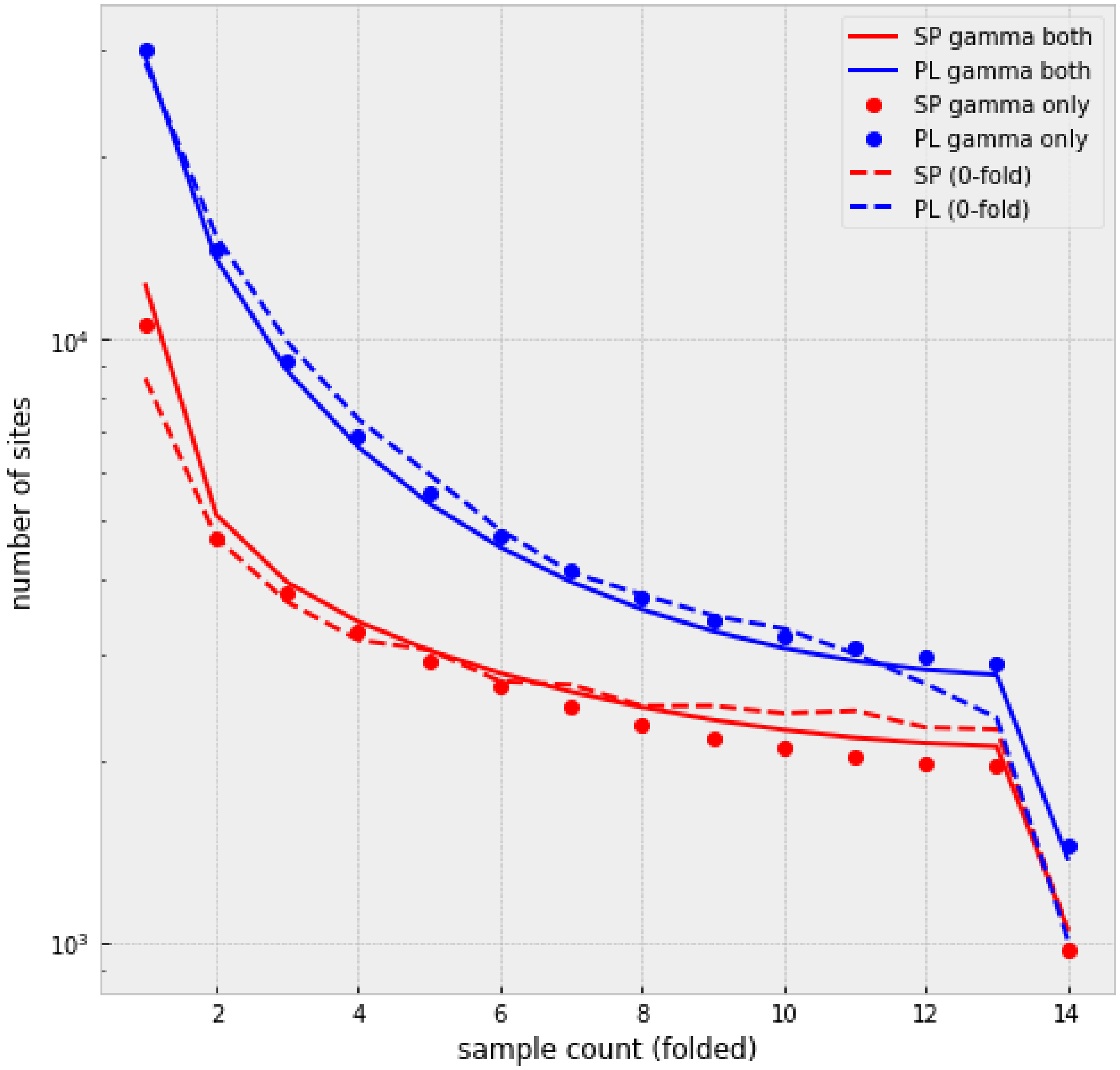


In [130]:

```
fig, ax = plt.subplots(figsize=(7,7))

plt.plot(fit_SP_both.fold()/np.sum(fit_SP_both.fold()), color="red", label=
"SP gamma both")
plt.plot(fit_PL_both.fold()/np.sum(fit_PL_both.fold()), color="blue", label
="PL gamma both")
```

```
plt.plot(fit_SP_only.fold()/np.sum(fit_SP_only.fold()),  "o", color="red",
label="SP gamma only")
plt.plot(fit_PL_only.fold()/np.sum(fit_PL_only.fold()), "o", color="blue",
label="PL gamma only")
plt.plot(data_SP_0fold/np.sum(data_SP_0fold), "--", color="red", label="SP
(0-fold)")
plt.plot(data_PL_0fold/np.sum(data_PL_0fold), "--", color="blue", label="PL
(0-fold)")


plt.yscale("log")
plt.ylabel("propotion of sites")
plt.xlabel("sample count (folded)")
plt.legend()
#fig.savefig("gamma_fits_theta_prop.pdf", bbox_inches="tight")
```

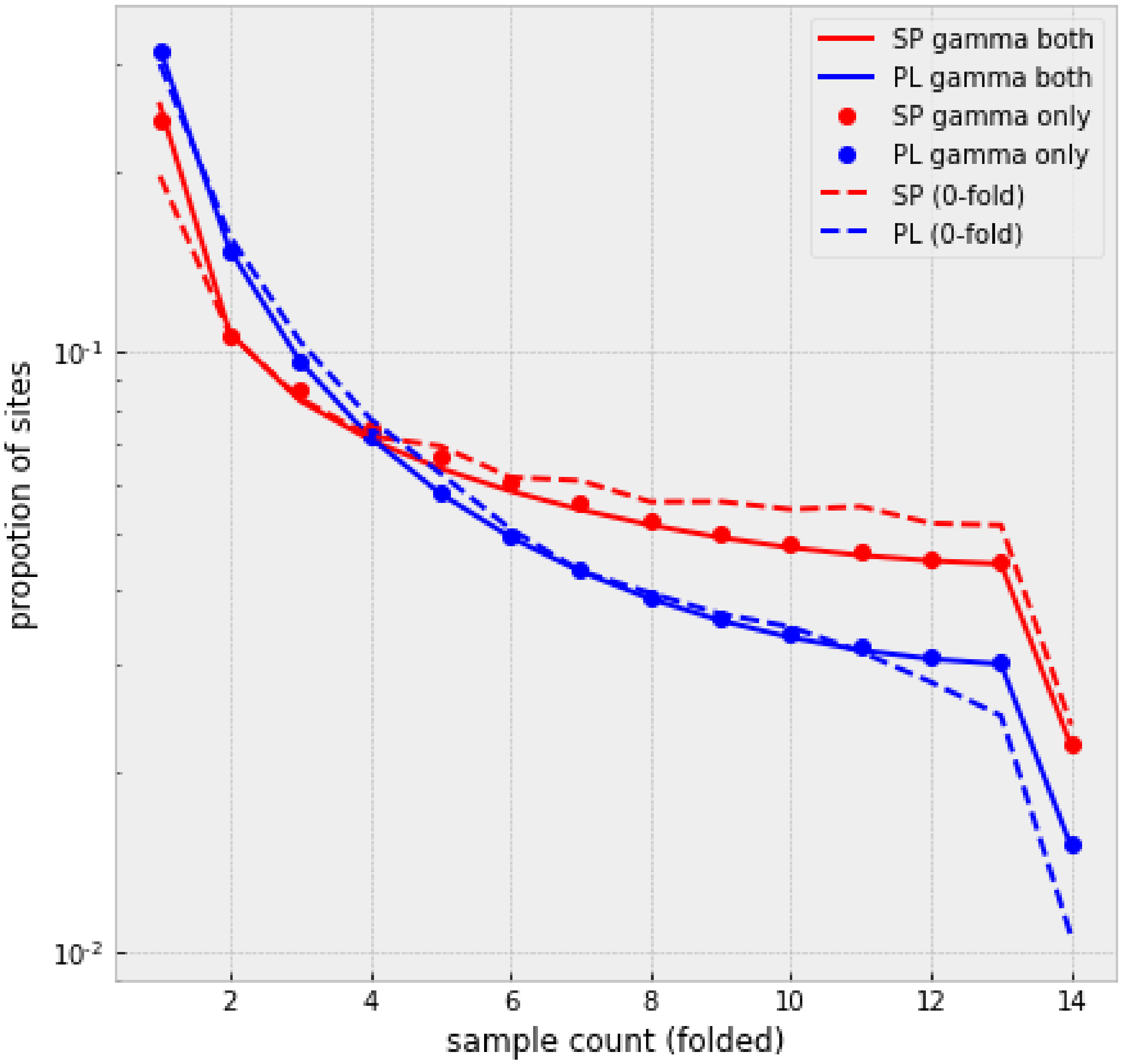


In [104]:

```
# visualising the observed SFS and expected SFS for the point estimate (mul
tinomial model)
fig, ax = plt.subplots(figsize=(8,8))
```

```
fit_SP_both = spectra_SP_opt.integrate([shape_set[246], scale_fit_set[246]]
, gamma_dist, theta_SP_opt_ns)
fit_PL_both = spectra_PL_opt.integrate([shape_set[246], scale_fit_set[246]]
, gamma_dist, theta_PL_opt_ns)

fit_SP_only = spectra_SP_opt.integrate([SP_shape_set[246], SP_scale_fit_set
[246]], gamma_dist, theta_SP_opt_ns)
fit_PL_only = spectra_PL_opt.integrate([PL_shape_set[246], PL_scale_fit_set
[246]], gamma_dist, theta_PL_opt_ns)

# 44% of mutations strongly enough selected to not show up in the sample
# we therefore multiply by 0.56
# this fits the multinomial DFE to the observed counts

plt.plot(fit_SP_both.fold()*.56,
         color="red", label="SP gamma both")
plt.plot(fit_PL_both.fold()*.56,
         color="blue", label="PL gamma both")
plt.plot(fit_SP_only.fold()*.56,
         "o", color="red", label="SP gamma only")
plt.plot(fit_PL_only.fold()*.56,
         "o", color="blue", label="PL gamma only")
plt.plot(data_SP_0fold, "--",
         color="red", label="SP (0-fold)")
plt.plot(data_PL_0fold, "--",
         color="blue", label="PL (0-fold)")

plt.yscale("log")
plt.ylabel("number of sites")
plt.xlabel("sample count (folded)")
plt.legend()
plt.title("Point DFE")
#fig.savefig("point_DFE_fits.pdf", bbox_inches="tight")

# the point estimate (multinomial model) has a very good fit to the data he
re
# this is true for both the joint as well as the independent optimization
```

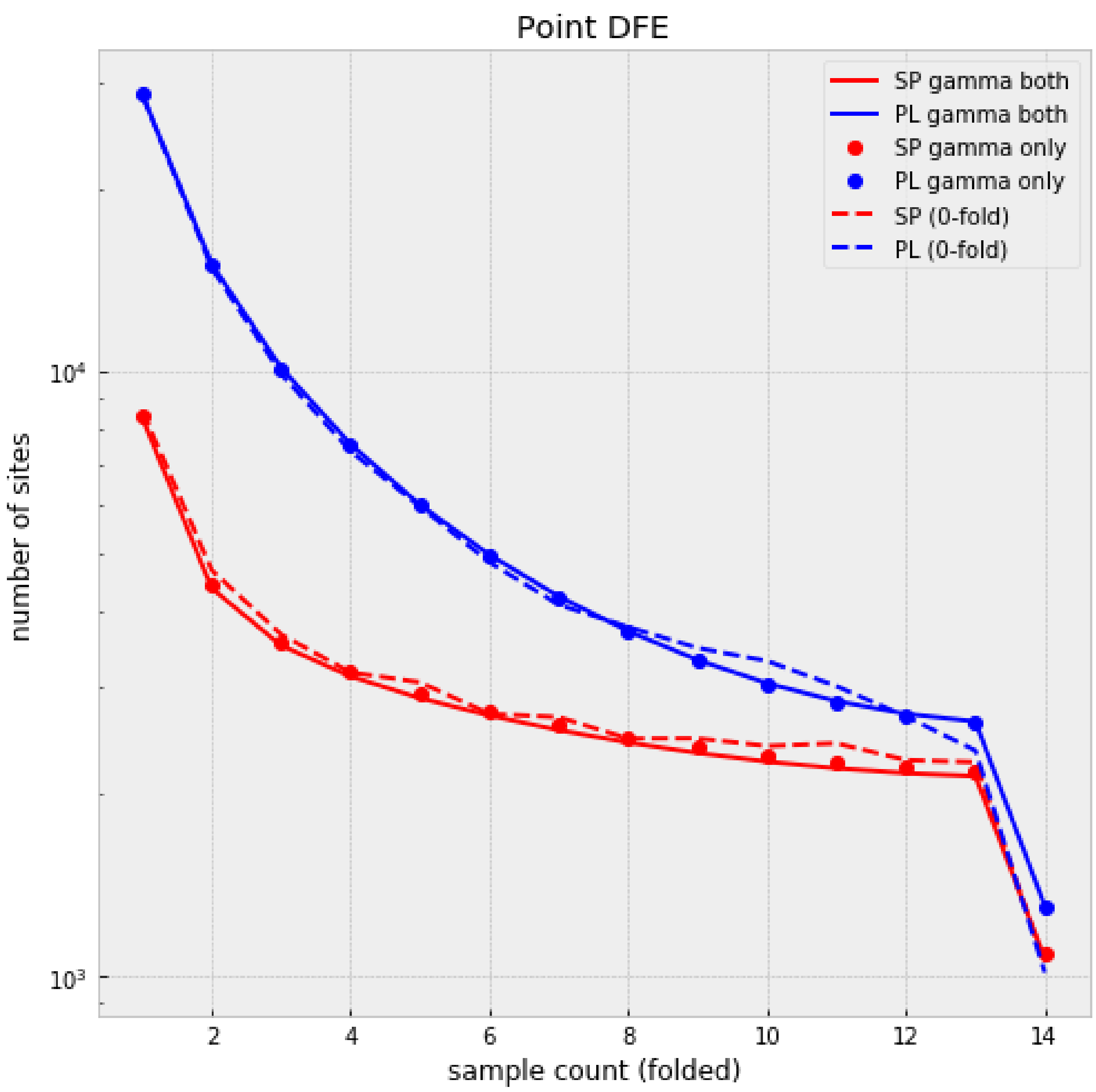


In [105]:

```
# visualising the cummulative distribution function (CDF) of the DFE
# the gamma distribution is calculated using the 'gamma.cdf' function and t
he previously calculated shape and scale parameters
# for SP two population sizes are shown, at the split with PL and the final
size

# calculating the CDF
from scipy.stats import gamma, lognorm
i = 246
Ns_SP_set = -spectra_SP_opt.gammas[::-1]
# point estimate for the multinomial model
DFE_cdf_both = gamma.cdf(Ns_SP_set, a=shape_set[i], scale=scale_fit_set[i])
# estimates from the Poisson model
DFE_cdf_both_t = gamma.cdf(Ns_SP_set, a=shape_set_t[min_both], scale=scale_
fit_set_t[min_both])
DFE_cdf_both_SP_t = gamma.cdf(Ns_SP_set, a=SP_shape_set_t[min_SP], scale=SP
_scale_fit_set_t[min_SP])
```

```
DFE_cdf_both_PL_t = gamma.cdf(Ns_SP_set, a=PL_shape_set_t[min_PL], scale=PL
_scale_fit_set_t[min_PL])

# plot CDF for the multinomial model
fig, ax = plt.subplots(figsize=(8,8))
ax.plot(Ns_SP_set*PLnu_opt, DFE_cdf_both, "-", color="blue", label="PL fina
l")
ax.plot(Ns_SP_set*SPnuF2_opt, DFE_cdf_both, "--", color="red", label="SP po
st-split")
ax.plot(Ns_SP_set*SPnuF1_opt, DFE_cdf_both, "-", color="red", label="SP fin
al")

ax.set(xscale="log", xlabel=r"$2N_e s$", ylabel="CDF", title="Multinomial C
DFs")
ax.legend()
#fig.savefig("SP_PL_DFE_multinom.pdf")
```

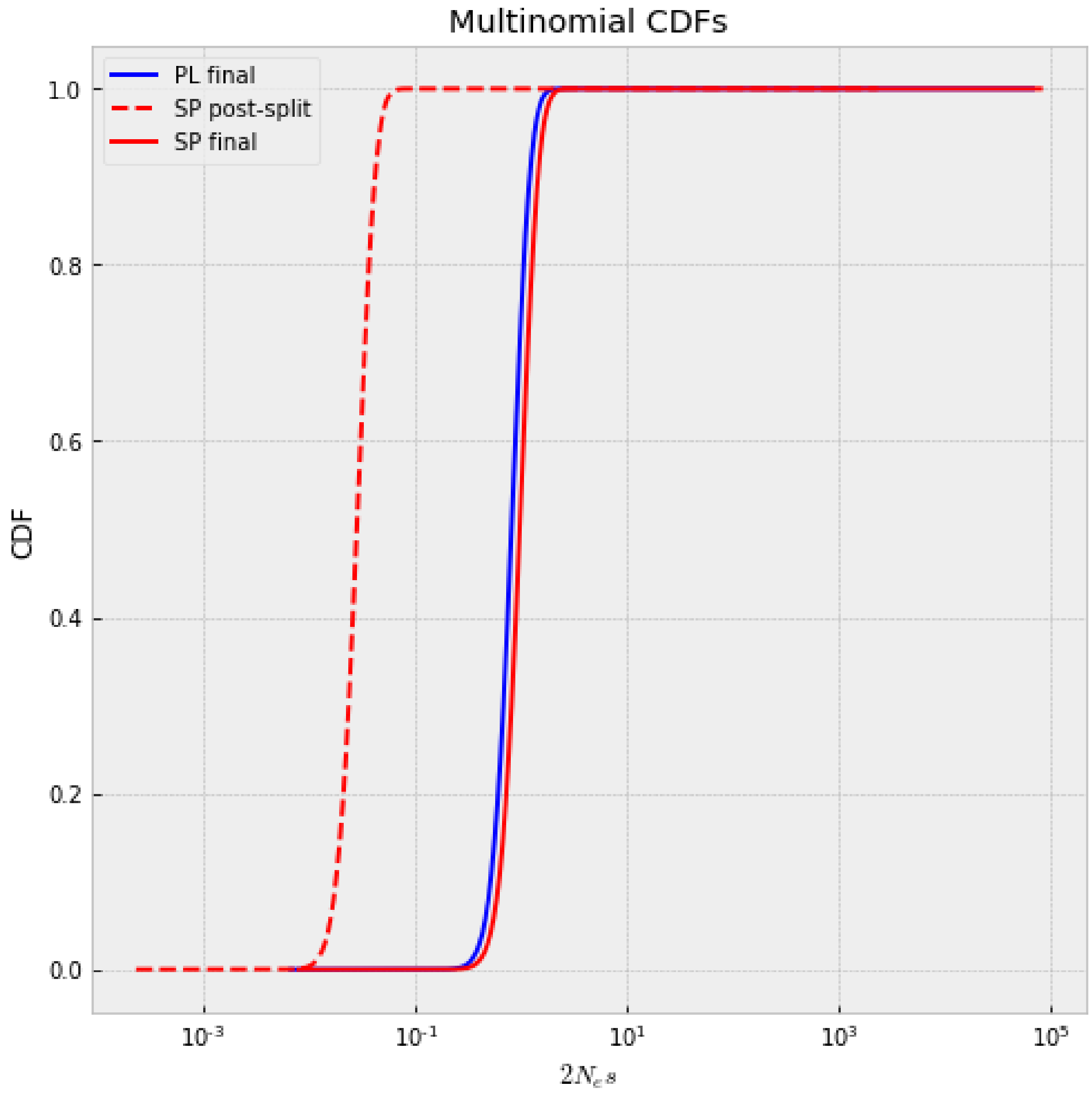


In [106]:

```
# plot CDF for the Poisson model (joint estimate)

fig, ax = plt.subplots(figsize=(8,8))
ax.plot(Ns_SP_set*PLnu_opt, DFE_cdf_both_t, "-", color="blue", label="PL")
ax.plot(Ns_SP_set*SPnuF2_opt, DFE_cdf_both_t, "--", color="red", label="SP
post-split")
ax.plot(Ns_SP_set*SPnuF1_opt, DFE_cdf_both_t, "-", color="red", label="SP f
inal")

ax.set(xscale="log", xlabel=r"$2N_e s$", ylabel="CDF", title=r"$\theta$ CDF
s both")
ax.legend()
#fig.savefig("SP_PL_DFE_theta_both.pdf")
```

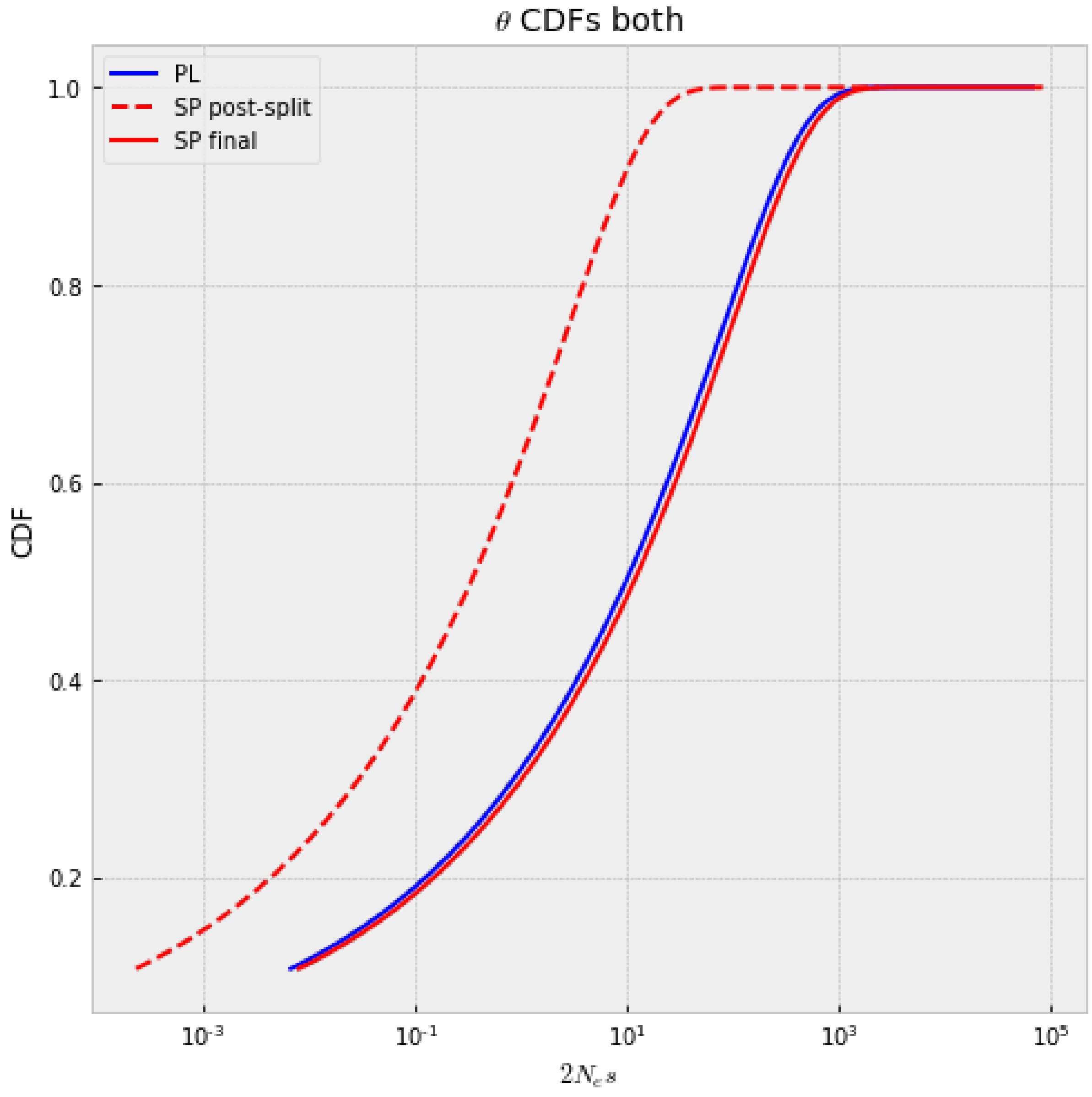


In [46]:

```
# calculate the proportion of segregating alleles for each bin of s for the
SFS
```

```
# for this we calculate the expected SFS for each value of s under the deog
raphic scenario
# and also the gamma distribution under the joint Poisson model for SP and
PL
# by applying Bayes' rule we can calculate the distribution of s for the va
riants in each count of the SFS

# calculate the expected SFS for a range of gammas
# this uses the demographic parameters and the selection function that was
previously defined
gamma_set = np.logspace(-1, 4, 100)
spectra_full_sel = []
for ii, gamma_tmp in enumerate(tqdm_notebook(gamma_set)):
    spectra_full_sel.append(three_epoch_growth_sel_both_new(demo_params_opt
+ [-gamma_tmp], 1, 100))
```

```
/home/ksteige/anaconda2/lib/python2.7/site-packages/ipykernel_launcher.py:1
0: TqdmDeprecationWarning: This function will be removed in tqdm==5.0.0
Please use `tqdm.notebook.tqdm` instead of `tqdm.tqdm_notebook`
  # Remove the CWD from sys.path while we load stuff.
```

In [47]:

```
# get the gamma distribution using the joint Poisson model
# using the shape and scale parameter with the best LL for both populations
# the gamma distribution is calculated by the 'gamma.cdf' function

F_gamma_both = gamma.cdf(gamma_set, a=shape_set_t[min_both], scale=scale_fi
t_set_t[min_both])
p_gamma_both = F_gamma_both - np.concatenate([[0], F_gamma_both[0:-1]])
p_gamma_both[-1] += 1 - p_gamma_both[-1]
```

In [48]:

```
# combine information of the expected SFS for each s and the gamma distribu
tion
# this allows us to get the proportion of s for each count of the SFS
PL_cDFE_set = []
SP_cDFE_set = []

# get proportion of s for each count of the SFS
from copy import deepcopy

for ii in range(14):
    cDFE_PL_tmp = np.zeros_like(gamma_set)
    cDFE_SP_tmp = np.zeros_like(gamma_set)
    for jj, gamma_tmp in enumerate(gamma_set):
        cDFE_PL_tmp[jj] = p_gamma_both[jj]*spectra_full_sel[jj].marginalize
([1]).fold()[ii+1]
        cDFE_SP_tmp[jj] = p_gamma_both[jj]*spectra_full_sel[jj].marginalize
([0]).fold()[ii+1]
    PL_cDFE_set.append(deepcopy(cDFE_PL_tmp/np.sum(cDFE_PL_tmp)))
    SP_cDFE_set.append(deepcopy(cDFE_SP_tmp/np.sum(cDFE_SP_tmp)))
```

```python
# get bins for s
gamma_bins = np.digitize(gamma_set, np.array([-1, 0.1, 1, 10, 100, 1e6]), r
ight=True)
bins = np.arange(1, 6)
PL_bin_prob_set = []
SP_bin_prob_set = []
for ii in range(14):
    PL_bin_probs = [0]*5
    SP_bin_probs = [0]*5
    for jj, bin_ind in enumerate(bins):
        PL_bin_probs[jj] = np.sum(PL_cDFE_set[ii][gamma_bins == bin_ind])
        SP_bin_probs[jj] = np.sum(SP_cDFE_set[ii][gamma_bins == bin_ind])
    PL_bin_probs = np.array(PL_bin_probs)
    SP_bin_probs = np.array(SP_bin_probs)
    PL_bin_probs[np.array(PL_bin_probs) < 0] = 0
    SP_bin_probs[np.array(SP_bin_probs) < 0] = 0
    PL_bin_prob_set.append(deepcopy(PL_bin_probs)/np.sum(PL_bin_probs))
    SP_bin_prob_set.append(deepcopy(SP_bin_probs)/np.sum(SP_bin_probs))

PL_bin_prob_set = np.array(PL_bin_prob_set)
SP_bin_prob_set = np.array(SP_bin_prob_set)
```

In [49]:

```python
from matplotlib.patches import Polygon
from matplotlib.collections import PatchCollection
from matplotlib import cm

cmap =["darkgrey","navy","cornflowerblue","mediumturquoise","darkorchid","i
ndigo"]

counts = np.arange(1, 15)

# plot binned s for each count of the SFS for PL
fig, axes = plt.subplots(1, 3, figsize=(27,7))

foo1 = Polygon(list(zip(counts, np.cumsum(PL_bin_prob_set,1)[:,0])) + [(14,
0.0), (1, 0.0)], color=cmap[5])
foo2 = Polygon(list(zip(counts, np.cumsum(PL_bin_prob_set,1)[:,0])) +
              list(reversed(list(zip(counts, np.cumsum(PL_bin_prob_set,1)[
:,1])))), color=cmap[4])
foo3 = Polygon(list(zip(counts, np.cumsum(PL_bin_prob_set,1)[:,1])) +
              list(reversed(list(zip(counts, np.cumsum(PL_bin_prob_set,1)[
:,2])))), color=cmap[3])
foo4 = Polygon(list(zip(counts, np.cumsum(PL_bin_prob_set,1)[:,2])) +
              list(reversed(list(zip(counts, np.cumsum(PL_bin_prob_set,1)[
:,3])))), color=cmap[2])
foo5 = Polygon(list(zip(counts, np.cumsum(PL_bin_prob_set,1)[:,3])) + [(1,1
)], color=cmap[1])
```

```
p = PatchCollection([foo1, foo2, foo3, foo4, foo5], match_original=True, cm
ap=cm.jet)
axes[0].add_collection(p)
for ii, _ in enumerate(bins):
    axes[0].scatter(counts, np.cumsum(PL_bin_prob_set,1)[:,ii], c=np.array(
[cmap[6-ii-1]]*14), edgecolor="black", s=60)

axes[0].set(ylim=[-0.05,1.05], ylabel="Proportion of segregating alleles",
xlabel="Sample count", title="PL")

# plot binned s for each count of the SFS for SP
foo1 = Polygon(list(zip(counts, np.cumsum(SP_bin_prob_set,1)[:,0])) + [(14,
0.), (1, 0.)], color=cmap[5])
foo2 = Polygon(list(zip(counts, np.cumsum(SP_bin_prob_set,1)[:,0])) +
               list(reversed(list(zip(counts, np.cumsum(SP_bin_prob_set,1)[
:,1])))), color=cmap[4])
foo3 = Polygon(list(zip(counts, np.cumsum(SP_bin_prob_set,1)[:,1])) +
               list(reversed(list(zip(counts, np.cumsum(SP_bin_prob_set,1)[
:,2])))), color=cmap[3])
foo4 = Polygon(list(zip(counts, np.cumsum(SP_bin_prob_set,1)[:,2])) +
               list(reversed(list(zip(counts, np.cumsum(SP_bin_prob_set,1)[
:,3])))), color=cmap[2])
foo5 = Polygon(list(zip(counts, np.cumsum(SP_bin_prob_set,1)[:,3])) + [(1,1
)], color=cmap[1])
p = PatchCollection([foo1, foo2, foo3, foo4, foo5], match_original=True, cm
ap=cm.jet)
axes[1].add_collection(p)
for ii, _ in enumerate(bins):
    axes[1].scatter(counts, np.cumsum(SP_bin_prob_set,1)[:,ii], c=np.array(
[cmap[6-ii-1]]*14), edgecolor="black", s=60)

axes[1].set(ylim=[-0.05,1.05], ylabel="Proportion of segregating alleles",
xlabel="Sample count", title="SP")

labels=[r"$\gamma \leq 0.1$", r"$0.1 > \gamma \leq 1$", r"$1 > \gamma \leq
10$", r"$10 > \gamma \leq 100$", r"$\gamma > 100$"]

# plot ratio PL / SP
for ii, _ in enumerate(bins):
    axes[2].plot(counts,(PL_bin_prob_set[:,ii]/SP_bin_prob_set[:,ii]), c=cm
ap[6-ii-1])
    axes[2].scatter(counts, (PL_bin_prob_set[:,ii]/SP_bin_prob_set[:,ii]),
c=np.array([cmap[6-ii-1]]*14), edgecolor="black", s=60, label=labels[ii])
axes[2].set(ylabel="PL/SP ratio", xlabel="Sample count")
axes[2].legend(loc=(.6, 0.1))

#fig.savefig("seg_effects_both_DFE_SP_PL.pdf", bbox_inches="tight")
```

Out[49]:

<matplotlib.legend.Legend at 0x7f51d375dd10>

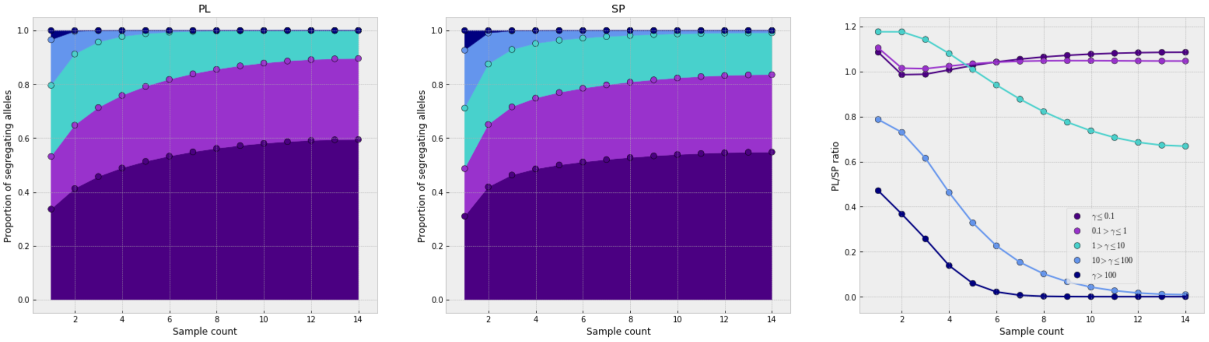


In [52]:

```
# calculate the burden difference between the populations
# using point estimate for s of -1.2 from the multinomial model to calculat
e the expected SFS
# the estimate of -1.2 is gained by the single populations as well as the j
oint estimate
# the selection function previously defined and the demographic parameters
are used here
# the difference in the burden between the populations is based on the cont
ribution of deleterious variants depending on their count in the population

sfs_single = three_epoch_growth_sel_both_new(demo_params_opt + [-1.2], 1, 2
000)
single_PL = sfs_single.marginalize([1])
single_SP = sfs_single.marginalize([0])
single_PL.mask=False
single_SP.mask=False

# 44% of mutations strongly enough selected to not show up in the sample
# we therefore multiply by 0.56
# this fits the multinomial DFE to the observed counts

fig, ax = plt.subplots(figsize=(8,8))
ax.plot(np.cumsum(single_PL*np.arange(29)/28*theta_PL_opt_ns*2*0.56) -
        np.cumsum(single_SP*np.arange(29)/28*theta_PL_opt_ns*2*0.56))
plt.yscale("symlog", linthreshy=100)
plt.ylabel("burden difference between PL and SP")
plt.xlabel("derived allele count")
plt.title("Point DFE")
#fig.savefig("single_DFE_burden_difference_PL_SP.pdf", bbox_inches="tight")
```

Out[52]:

<matplotlib.text.Text at 0x7f51c1a351d0>

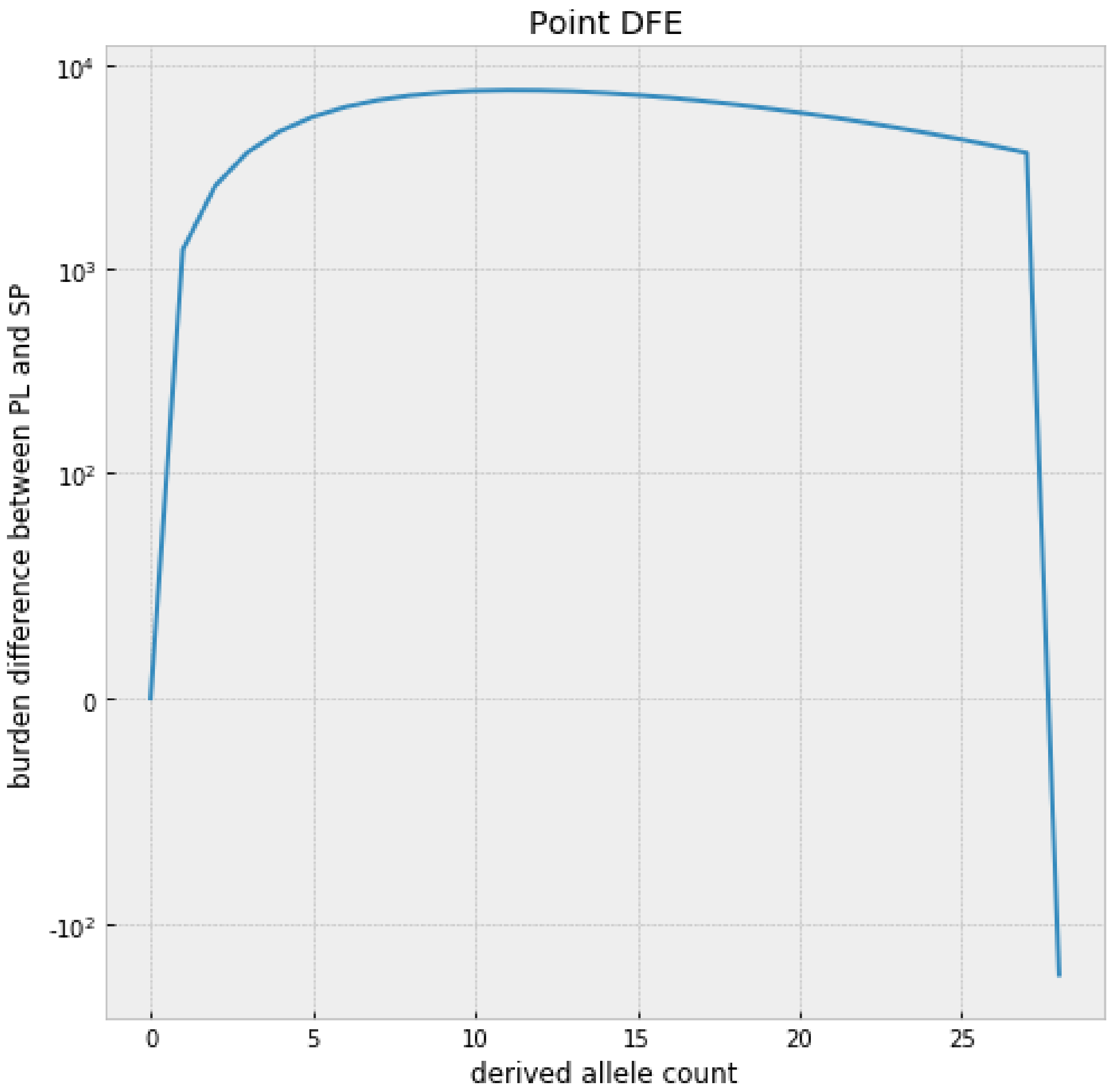


In [53]:

```
# also analysing the data when dominance h=0
#
# define functions to generate SFS under selection
# this needs the demographic parameters and a value of gamma
# function includes dadi functions for population size change and split of
populations
# to speed up calculations a minimum gamma value was chosen for very strong
selection
# dominance h set to 0 here
#
# SFS for a single population as output
def three_epoch_growth_sel_new_h0(params, ns, pts):
    nuB, nuF2, nuF1, TB, TF, TI, pi, gamma = params
    T1 = (TB+TF+TI)*pi
    T2 = (TB+TF+TI)*(1-pi)
    xx = dadi.Numerics.default_grid(pts)
    min_gamma = -500
```

```
    if gamma < min_gamma:
        phi = dadi.PhiManip.phi_1D(xx, gamma=min_gamma, h=0)
    else:
        phi = dadi.PhiManip.phi_1D(xx, gamma=gamma, h=0)

    if nuB*gamma < min_gamma:
        phi = dadi.Integration.one_pop(phi, xx, T1, nuB, gamma=min_gamma, h
=0)
    else:
        phi = dadi.Integration.one_pop(phi, xx, T1, nuB, gamma=gamma, h=0)

    if gamma == 0:
        nu_func = lambda t: nuF2*(nuF1/nuF2)**(t/TI)
    else:
        nu_func = lambda t: np.where(gamma*nuF2*(nuF1/nuF2)**(t/T2) < min_g
amma,
                                     np.abs(min_gamma/gamma),
                                     nuF2*(nuF1/nuF2)**(t/T2))

    phi = dadi.Integration.one_pop(phi, xx, T2, nu_func, gamma=gamma, h=0)
    fs = dadi.Spectrum.from_phi(phi, (28,), (xx,))
    return fs

# SFS for both populations as output
def three_epoch_growth_sel_both_new_h0(params, ns, pts):
    min_gamma = -500
    nuB, PLnuF2, PLnuF1, SPnuF2, SPnuF1, TB, TF, TI, pi, gamma = params
    T1 = (TB+TF+TI)*pi
    T2 = (TB+TF+TI)*(1-pi)
    # define grid
    xx = dadi.Numerics.default_grid(pts)
    # equilibrium ancestral population
    if gamma < min_gamma:
        phi = dadi.PhiManip.phi_1D(xx, gamma=min_gamma, h=0)
    else:
        phi = dadi.PhiManip.phi_1D(xx, gamma=gamma, h=0)
    # population reduction in ancestral pop
    if nuB*gamma < min_gamma:
        phi_2 = dadi.Integration.one_pop(phi, xx, T1, nuB, gamma=min_gamma,
h=0)
    else:
        phi_2 = dadi.Integration.one_pop(phi, xx, T1, nuB, gamma=gamma, h=0
)
    # split between SP and PL and migration
    phi_3 = dadi.PhiManip.phi_1D_to_2D(xx, phi_2)
    # changes in population size in PL and SP, no migration
    if gamma == 0:
        nu_func1 = lambda t: PLnuF2*(PLnuF1/PLnuF2)**(t/T2)
        nu_func2 = lambda t: SPnuF2*(SPnuF1/SPnuF2)**(t/T2)
```

```
    else:
        nu_func1 = lambda t: np.where(gamma*PLnuF2*(PLnuF1/PLnuF2)**(t/T2)
< min_gamma,
                                     np.abs(min_gamma/gamma),
                                     PLnuF2*(PLnuF1/PLnuF2)**(t/T2))
        nu_func2 = lambda t: np.where(gamma*SPnuF2*(SPnuF1/SPnuF2)**(t/T2)
< min_gamma,
                                     np.abs(min_gamma/gamma),
                                     SPnuF2*(SPnuF1/SPnuF2)**(t/T2))
    phi_4 = dadi.Integration.two_pops(phi_3, xx, T2, nu1=nu_func1, nu2=nu_f
unc2, gamma1=gamma, gamma2=gamma, h1=0, h2=0)
    fs = dadi.Spectrum.from_phi(phi_4, (28,28), (xx,xx), pop_ids=["PL","SP"
])
    return fs
```

In [ ]:

```
# define array based on positions in the SFS
ns = np.array([28])

# run spectra function from fitdadi
# this allows to get the estimated SFS under the demographic parametrs for
a range of gammas
# this uses the previously defined function "three_epoch_growth_sel_new" an
d the simplified demographic parameters

# run for the SP population
spectra_SP_opt_h0 = spectra(demo_params_opt_SP, ns, three_epoch_growth_sel_
new_h0, pts=1000,
                        int_bounds=(0.01, 350), Npts=500, echo=False, mp=Tr
ue)
# save object as the calculation takes some time
#pickle.dump(spectra_SP_opt_h0, open("spectra_SP_break_model_new_h0.sp","wb
"))

# run for the PL population
spectra_PL_opt_h0 = spectra(demo_params_opt_PL, ns, three_epoch_growth_sel_
new_h0, pts=1000,
                        int_bounds=(0.01, 350), Npts=500, echo=False, mp=Tr
ue)
#pickle.dump(spectra_PL_opt_h0, open("spectra_PL_break_model_new_h0.sp","wb
"))

# int_bounds reduced
# values higher lead to nan for the SFS
# they were therefore excluded here

# load the object if previously saved
#spectra_SP_opt_h0 = pickle.load(open("spectra_SP_break_model_new_h0.sp"))
#spectra_PL_opt_h0 = pickle.load(open("spectra_PL_break_model_new_h0.sp"))
```

In [55]:

```
# fit the DFE by estimating the shape and scale of the gamma distribution
# optimization steps in fitdadi available, but we used scipy optimization a
s it allows more flexibility
# method for optimization in scipy was SLSQP
# for the optimization both a multinomial model (without using theta) and a
Poisson model (including theta) were used
# the optimization can be done both using a single population and a joint e
stimate (for both populations)
# the saved output from the fitdadi 'spectra' function is needed here

# here we redo the optimization steps for h=0

# scipy optimization for SP under the multinomial model
SP_shape_set_h0 = np.logspace(start=-1, stop=1.0, num=250)
SP_scale_fit_set_h0 = np.ones_like(SP_shape_set_h0)
SP_llhood_fit_set_h0 = np.ones_like(SP_shape_set_h0)
for ii, shape in enumerate(SP_shape_set_h0):
    def obj_funct(scale):
        sfs_SP_h0 = spectra_SP_opt_h0.integrate([shape, scale], gamma_dist,
1).fold()
        return -Inference.ll_multinom(sfs_SP_h0, data_SP_0fold)
    fit = scipy.optimize.minimize(obj_funct, [SP_scale_fit_set_h0[ii-1]], m
ethod="SLSQP", bounds=[(np.exp(-8), np.exp(20))])
    SP_scale_fit_set_h0[ii] = fit["x"][0]
    SP_llhood_fit_set_h0[ii] = fit["fun"]
```

In [107]:

```
# plot the LL vs the shape parameter
fig, ax = plt.subplots(figsize=(7,4))
plt.plot(SP_shape_set_h0, SP_llhood_fit_set_h0-np.min(SP_llhood_fit_set_h0)
)
plt.xscale("log")
plt.yscale("symlog", linthreshy=1)
plt.ylabel(r"$-LL-\min(LL)$")
plt.xlabel("shape parameter")
plt.title("SP gamma fit")
#fig.savefig("SP_only_gamma_llhood_h0.pdf", bbox_inches="tight")
```

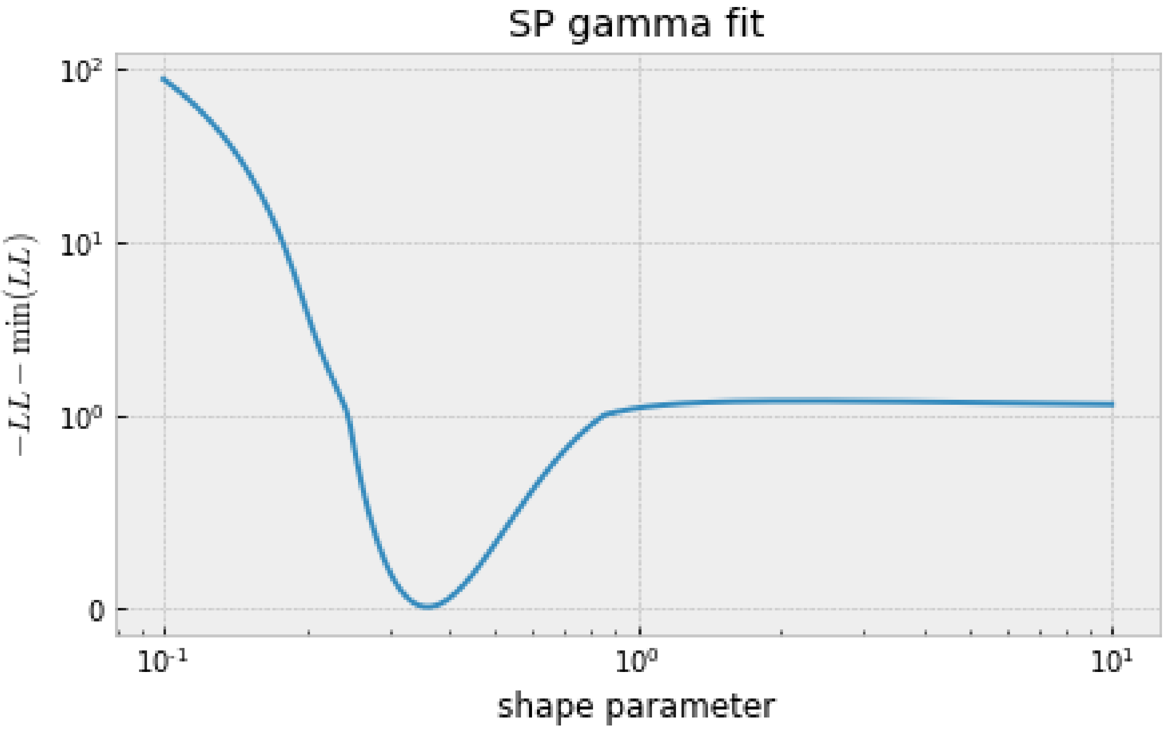


In [137]:

```
# plot the scale vs the shape parameter
fig, ax = plt.subplots(figsize=(7,4))
plt.plot(SP_shape_set_h0, SP_scale_fit_set_h0)
plt.xscale("log")
plt.yscale("symlog", linthreshy=2e4)
#plt.ylim([0,500])
plt.ylabel("scale parameter")
plt.xlabel("shape parameter")
plt.title(r"SP gamma fit $\theta$")
#fig.savefig("SP_only_gamma_scale_h0.pdf", bbox_inches="tight")
```

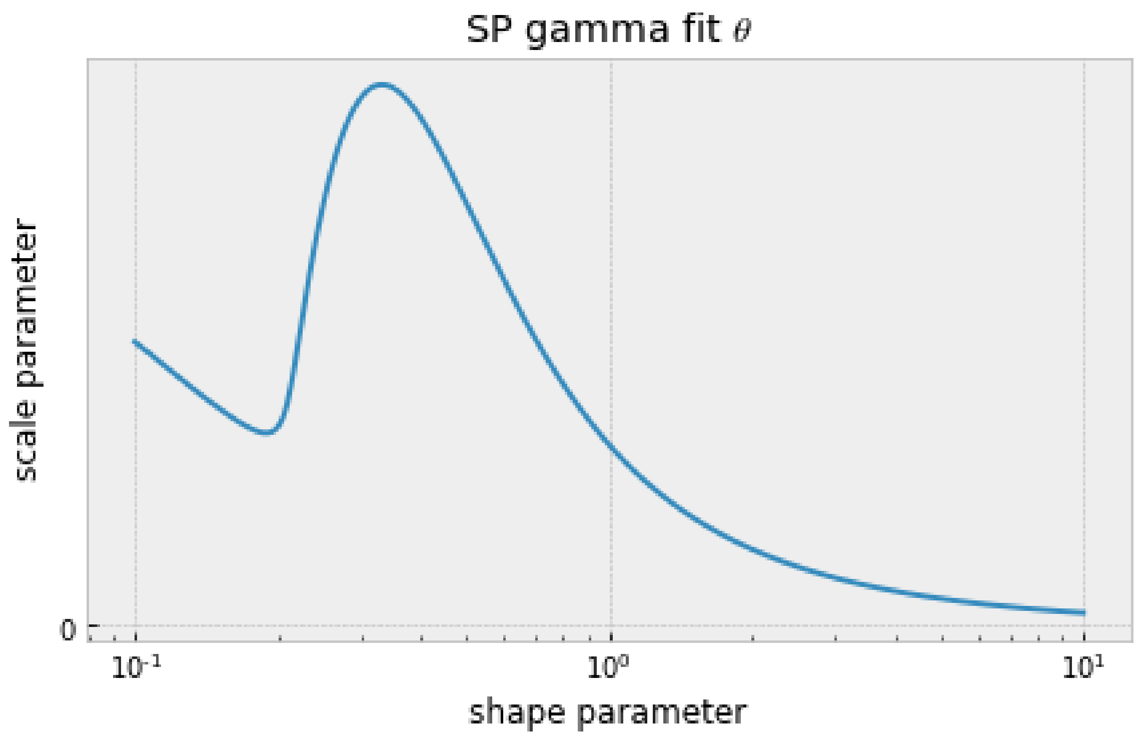


In [58]:

```
# scipy optimization for SP under the Poisson model including theta
SP_shape_set_t_h0 = np.logspace(start=-2, stop=0, num=150)
SP_scale_fit_set_t_h0 = np.ones_like(SP_shape_set_t_h0)
SP_llhood_fit_set_t_h0 = np.ones_like(SP_shape_set_t_h0)
for ii, shape in enumerate(SP_shape_set_t_h0):
    def obj_funct(scale):
        sfs_SP_h0 = spectra_SP_opt_h0.integrate([shape, np.exp(scale)], gam
ma_dist, theta_SP_opt_ns).fold()
        return -Inference.ll(sfs_SP_h0, data_SP_0fold)
    fit = scipy.optimize.minimize(obj_funct, [np.log(100)], method="SLSQP",
bounds=[(-8, 20)])
    SP_scale_fit_set_t_h0[ii] = np.exp(fit["x"][0])
    SP_llhood_fit_set_t_h0[ii] = fit["fun"]
```

In [109]:

```
# plot the LL vs the shape parameter
fig, ax = plt.subplots(figsize=(7,4))
plt.plot(SP_shape_set_t_h0, SP_llhood_fit_set_t_h0-np.min(SP_llhood_fit_set
_t_h0), "-")
plt.xscale("log")
plt.yscale("symlog", linthreshy=10)
plt.ylabel(r"$-LL-\min(LL)$")
plt.xlabel("shape parameter")
plt.title(r"SP gamma fit $\theta$")
#fig.savefig("SP_only_gamma_llhood_theta_h0.pdf", bbox_inches="tight")
```

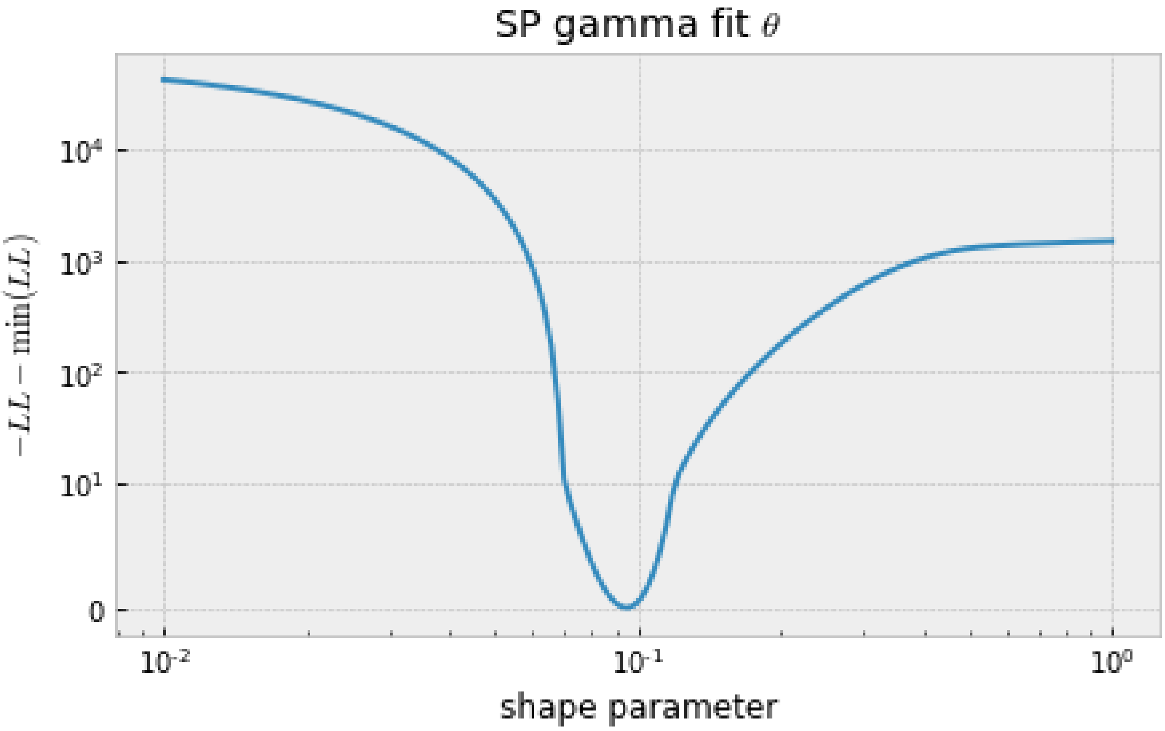


In [110]:

```
# plot the scale vs the shape parameter
fig, ax = plt.subplots(figsize=(7,4))
plt.plot(SP_shape_set_t_h0, SP_scale_fit_set_t_h0)
plt.xscale("log")
plt.yscale("symlog", linthreshy=2e4)
#plt.ylim([0,500])
plt.ylabel("scale parameter")
plt.xlabel("shape parameter")
plt.title(r"SP gamma fit $\theta$")
#fig.savefig("SP_only_gamma_scale_theta_h0.pdf", bbox_inches="tight")
```

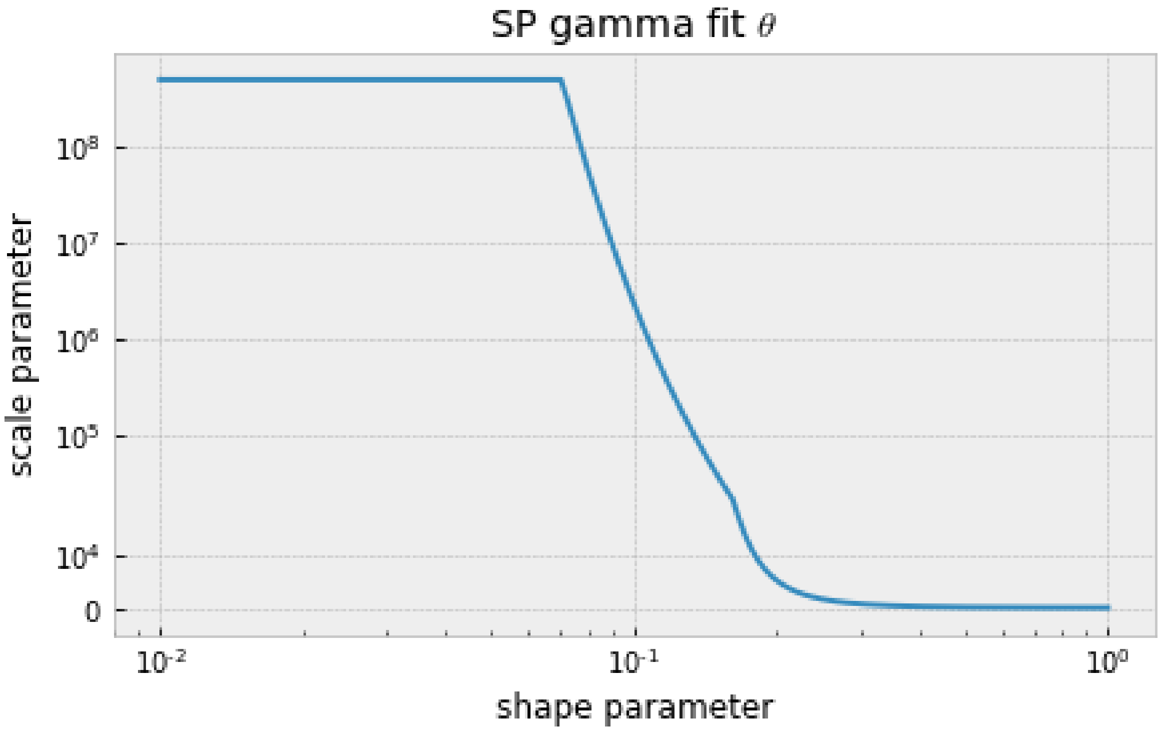


In [61]:

```
# scipy optimization for PL under the multinomial model

PL_shape_set_h0 = np.logspace(start=-1, stop=1, num=250)
PL_scale_fit_set_h0 = np.ones_like(PL_shape_set_h0)
PL_llhood_fit_set_h0 = np.ones_like(PL_shape_set_h0)
for ii, shape in enumerate(PL_shape_set_h0):
    def obj_funct(scale):
        sfs_PL_h0 = spectra_PL_opt_h0.integrate([shape, scale], gamma_dist,
1).fold()
        return -Inference.ll_multinom(sfs_PL_h0, data_PL_0fold)
    fit = scipy.optimize.minimize(obj_funct, [PL_scale_fit_set_h0[ii-1]], m
ethod="SLSQP", bounds=[(np.exp(-8), np.exp(20))])
    PL_scale_fit_set_h0[ii] = fit["x"][0]
    PL_llhood_fit_set_h0[ii] = fit["fun"]
```

In [111]:

```
# plot the LL vs the shape parameter
fig, ax = plt.subplots(figsize=(7,4))
plt.plot(PL_shape_set_h0, PL_llhood_fit_set_h0-np.min(PL_llhood_fit_set_h0)
)
plt.xscale("log")
plt.yscale("symlog", linthreshy=10)
plt.ylabel(r"$-LL-\min(LL)$")
plt.xlabel("shape parameter")
plt.title("PL gamma fit")
#fig.savefig("PL_only_gamma_llhood_h0.pdf", bbox_inches="tight")
```

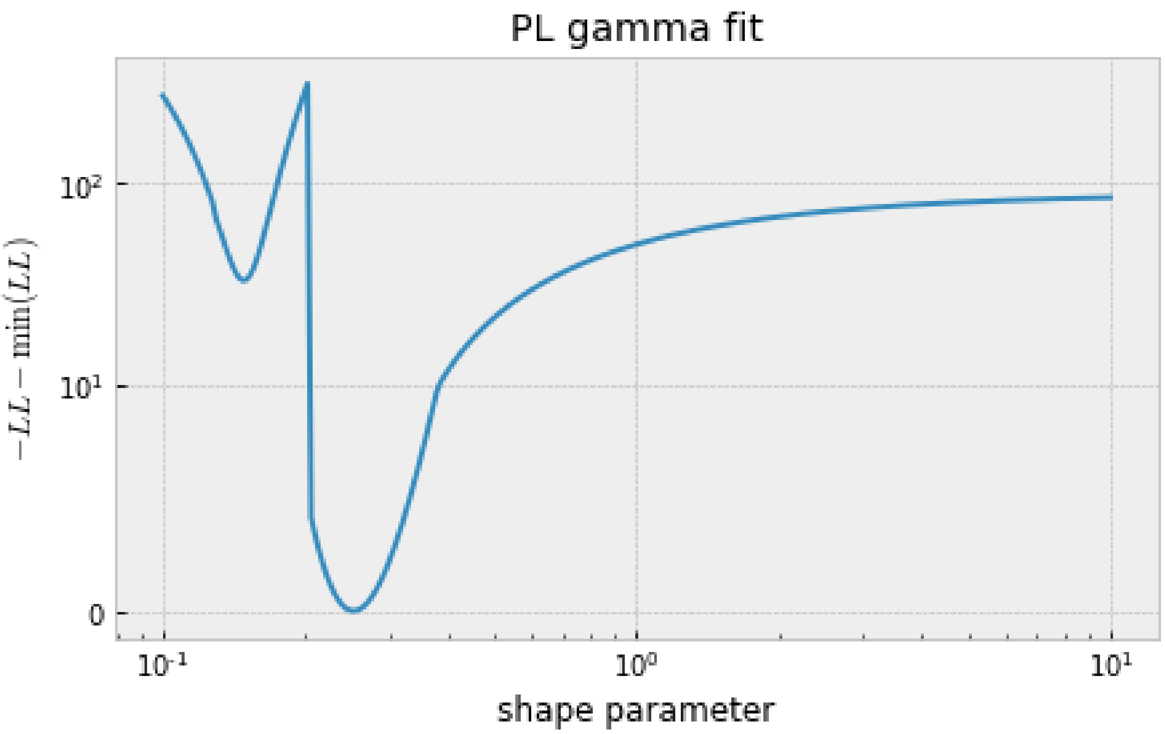


In [112]:

```
# plot the scale vs the shape parameter
fig, ax = plt.subplots(figsize=(7,4))
plt.plot(PL_shape_set_h0, PL_scale_fit_set_h0)
plt.xscale("log")
plt.yscale("symlog", linthreshy=1e3)
#plt.ylim([0,500])
plt.ylabel("scale parameter")
plt.xlabel("shape parameter")
plt.title(r"PL gamma fit $\theta$")
#fig.savefig("PL_only_gamma_scale_h0.pdf", bbox_inches="tight")
```

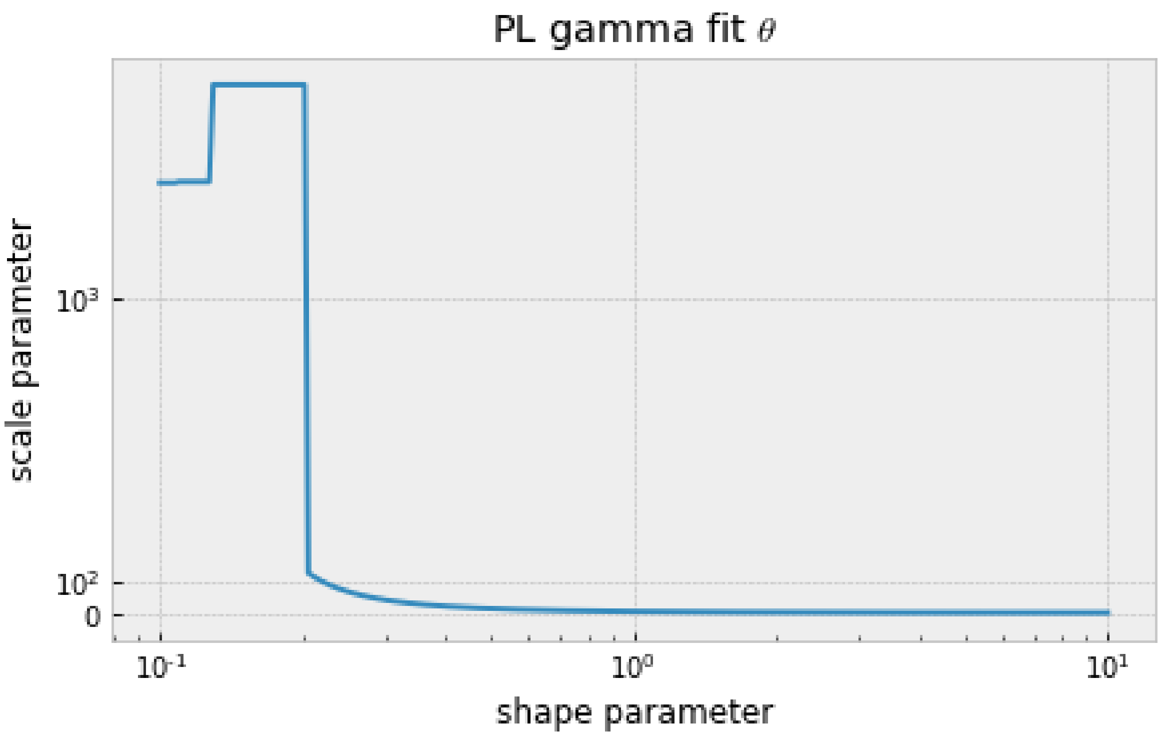


In [64]:

```
# scipy optimization for PL under the Poisson model including theta

PL_shape_set_t_h0 = np.logspace(start=-2, stop=0, num=250)
PL_scale_fit_set_t_h0 = np.ones_like(PL_shape_set_t_h0)
PL_llhood_fit_set_t_h0 = np.ones_like(PL_shape_set_t_h0)
for ii, shape in enumerate(PL_shape_set_t_h0):
    def obj_funct(scale):
        sfs_PL_h0 = spectra_PL_opt_h0.integrate([shape, np.exp(scale)], gam
ma_dist, theta_PL_opt_ns).fold()
        return -Inference.ll(sfs_PL_h0, data_PL_0fold)
    fit = scipy.optimize.minimize(obj_funct, [np.log(100)], method="L-BFGS-
B", bounds=[(-8, 20)])
    PL_scale_fit_set_t_h0[ii] = np.exp(fit["x"][0])
    PL_llhood_fit_set_t_h0[ii] = fit["fun"]
```

In [113]:

```
# plot the LL vs the shape parameter
fig, ax = plt.subplots(figsize=(7,4))
plt.plot(PL_shape_set_t_h0, PL_llhood_fit_set_t_h0-np.min(PL_llhood_fit_set
_t_h0))
plt.xscale("log")
plt.yscale("symlog", linthreshy=10)
plt.ylabel(r"$-LL-\min(LL)$")
plt.xlabel("shape parameter")
plt.title(r"PL gamma fit $\theta$")
#fig.savefig("PL_only_gamma_llhood_theta_h0.pdf", bbox_inches="tight")
```

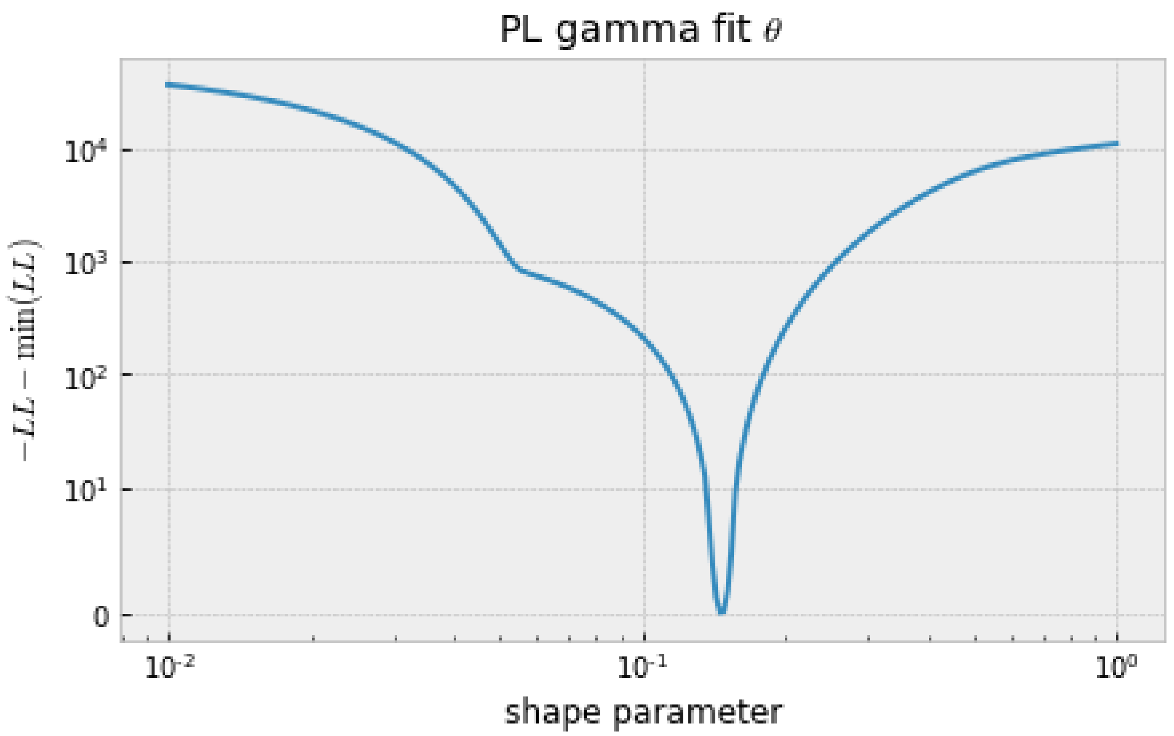


In [114]:

```
# plot the scale vs the shape parameter
fig, ax = plt.subplots(figsize=(7,4))
plt.plot(PL_shape_set_t_h0, PL_scale_fit_set_t_h0)
plt.xscale("log")
plt.yscale("symlog", linthreshy=1e3)
#plt.ylim([0,500])
plt.ylabel("scale parameter")
plt.xlabel("shape parameter")
plt.title(r"PL gamma fit $\theta$")
#fig.savefig("PL_only_gamma_scale_theta_h0.pdf", bbox_inches="tight")
```

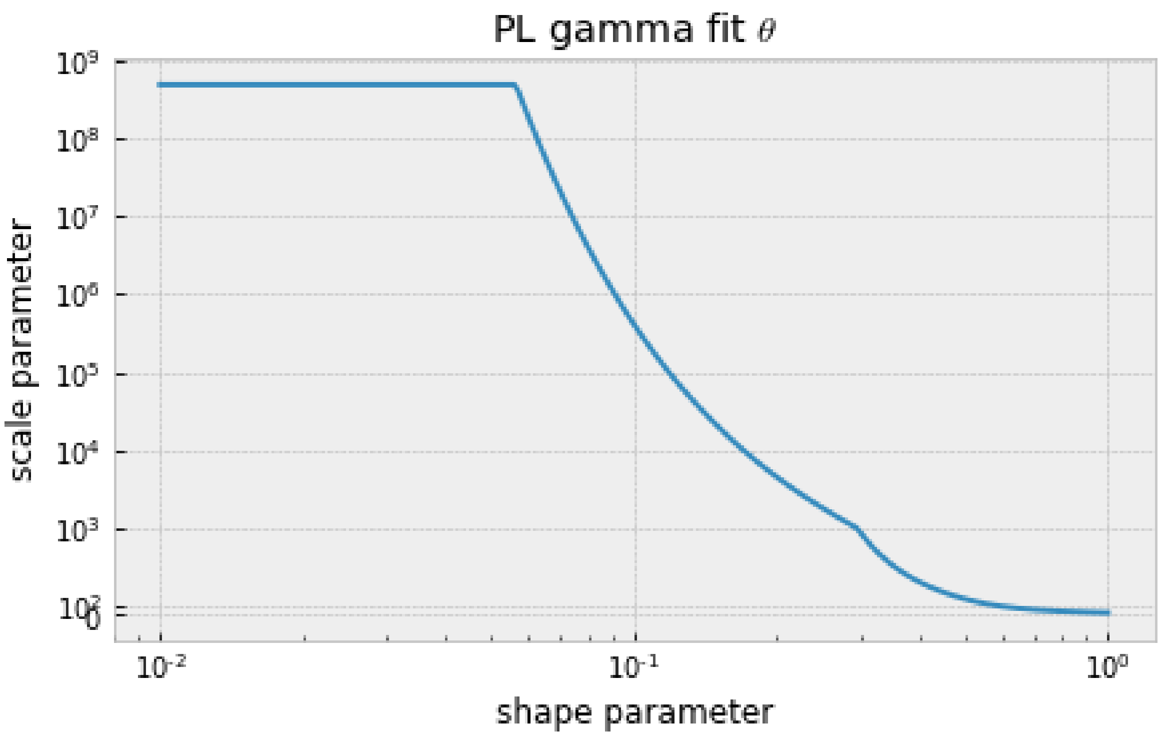


In [67]:

```
# joint scipy optimization for SP and PL under the multinomial model
shape_set_h0 = np.logspace(start=-1, stop=1.0, num=250)
scale_fit_set_h0 = np.ones_like(shape_set_h0)
llhood_fit_set_h0 = np.ones_like(shape_set_h0)
for ii, shape in enumerate(shape_set):
    def obj_funct(scale):
        sfs_SP_h0 = spectra_SP_opt_h0.integrate([shape, scale], gamma_dist,
1).fold()
        sfs_PL_h0 = spectra_PL_opt_h0.integrate([shape, scale], gamma_dist,
1).fold()
        return -Inference.ll_multinom(sfs_SP_h0, data_SP_0fold) - Inference
.ll_multinom(sfs_PL_h0, data_PL_0fold)
    fit = scipy.optimize.minimize(obj_funct, [scale_fit_set_h0[ii-1]], meth
od="SLSQP", bounds=[(np.exp(-8), np.exp(20))])
    scale_fit_set_h0[ii] = fit["x"][0]
    llhood_fit_set_h0[ii] = fit["fun"]
```

In [138]:

```
# plot the LL vs the shape parameter
fig, ax = plt.subplots(figsize=(7,4))
plt.plot(shape_set_h0, llhood_fit_set_h0-np.min(llhood_fit_set_h0))
plt.xscale("log")
plt.yscale("symlog", linthreshy=10)
plt.ylabel(r"$-LL-\min(LL)$")
plt.xlabel("shape parameter")
plt.title("Both gamma fit")
#fig.savefig("both_gamma_llhood_h0.pdf", bbox_inches="tight")
```

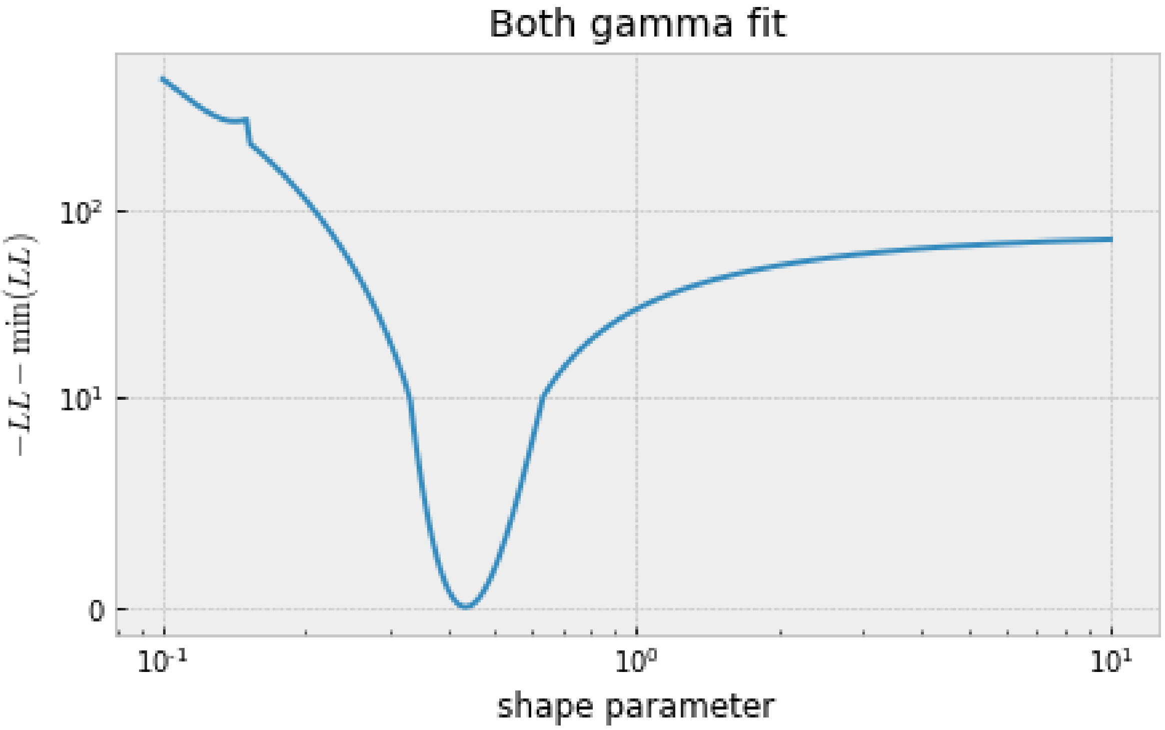


In [115]:

```
# plot the scale vs the shape parameter
fig, ax = plt.subplots(figsize=(7,4))
plt.plot(shape_set_h0, scale_fit_set_h0)
plt.xscale("log")
plt.yscale("symlog", linthreshy=1e3)
#plt.ylim([0,500])
plt.ylabel("scale parameter")
plt.xlabel("shape parameter")
plt.title(r"both gamma fit")
#fig.savefig("both_gamma_scale_h0.pdf", bbox_inches="tight")
```

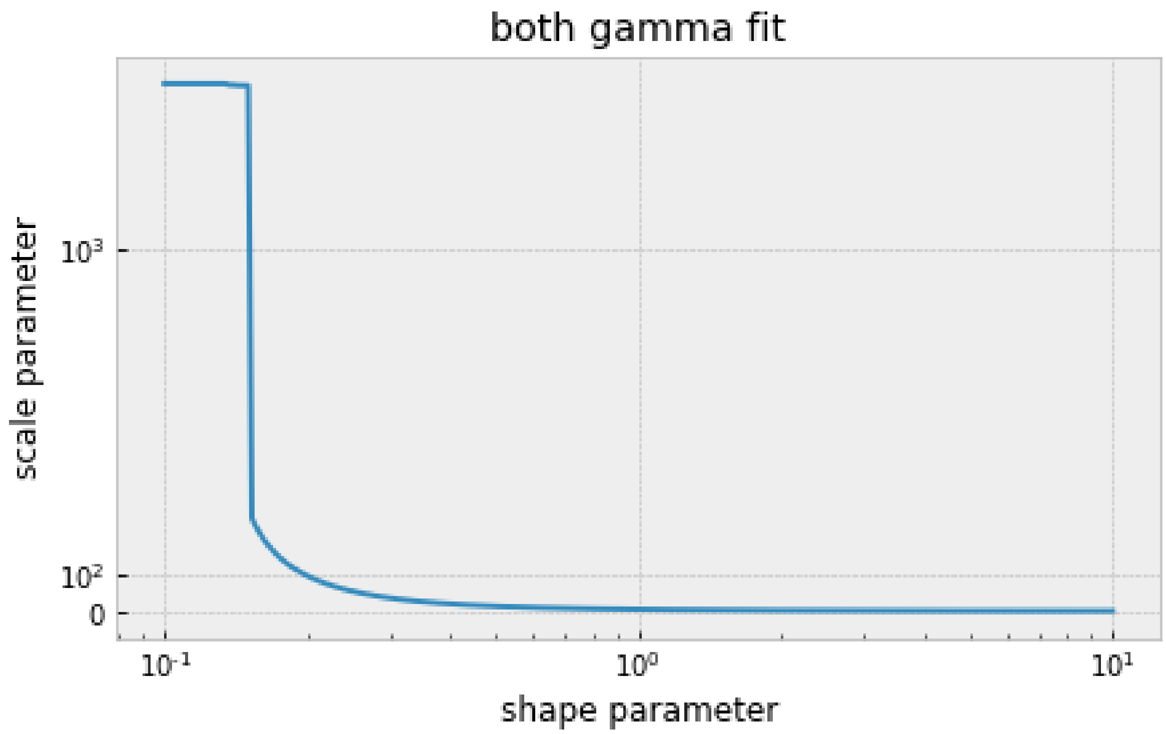


In [75]:

```
# joint scipy optimization for SP and PL under the Poisson model including
theta

shape_set_t_h0 = np.logspace(start=-2, stop=0, num=250)
scale_fit_set_t_h0 = np.ones_like(shape_set_t_h0)
llhood_fit_set_t_h0 = np.zeros_like(shape_set_t_h0)
for ii, shape in enumerate(shape_set_t_h0):
    def obj_funct(scale):
        sfs_SP_h0 = spectra_SP_opt_h0.integrate([shape, np.exp(scale)], gam
ma_dist, theta_SP_opt_ns).fold()
        sfs_PL_h0 = spectra_PL_opt_h0.integrate([shape, np.exp(scale)], gam
ma_dist, theta_PL_opt_ns).fold()
        return -Inference.ll(sfs_SP_h0, data_SP_0fold) - Inference.ll(sfs_P
L_h0, data_PL_0fold)
    fit = scipy.optimize.minimize(obj_funct, [np.log(100)], method="SLSQP",
bounds=[(-8, 20)])
    scale_fit_set_t_h0[ii] = np.exp(fit["x"][0])
    llhood_fit_set_t_h0[ii] = fit["fun"]
```

In [116]:

```
# plot the LL vs the shape parameter
fig, ax = plt.subplots(figsize=(7,4))
plt.plot(shape_set_t_h0, llhood_fit_set_t_h0-np.min(llhood_fit_set_t_h0))
plt.xscale("log")
plt.yscale("symlog", linthreshy=10)
plt.ylabel(r"$-LL-\min(LL)$")
plt.xlabel("shape parameter")
plt.title(r"Both gamma fit $\theta$")
#fig.savefig("both_gamma_llhood_theta_h0.pdf", bbox_inches="tight")
```

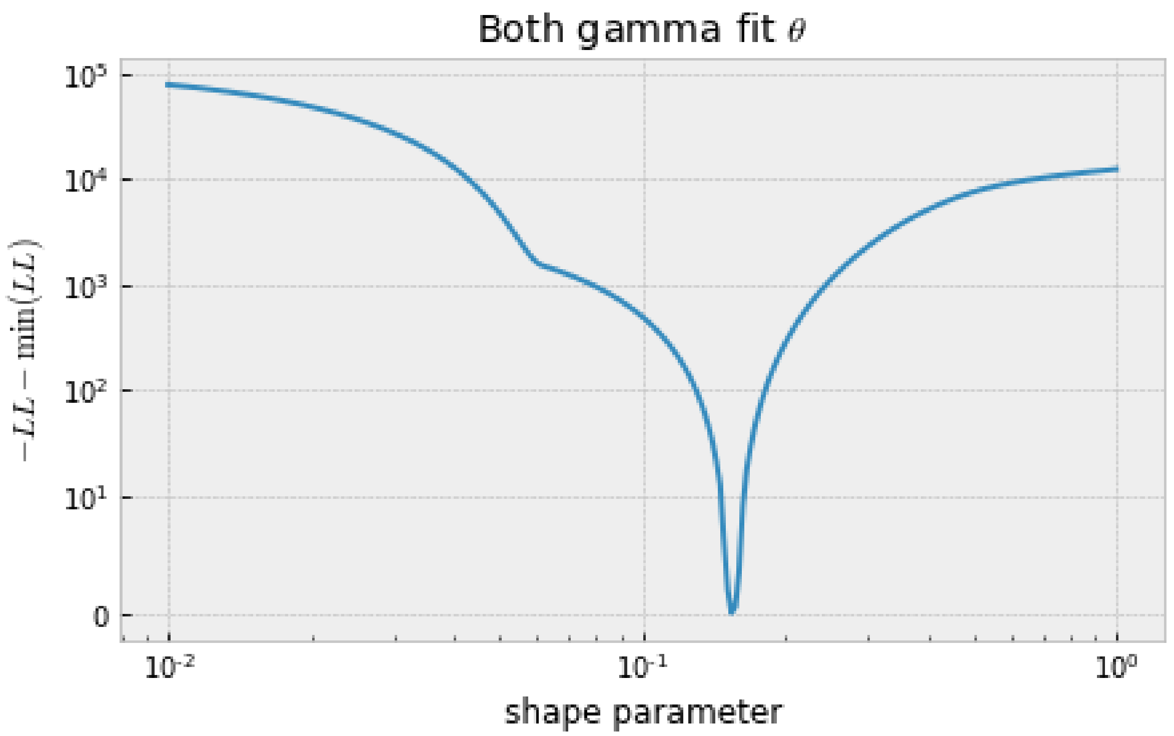


In [117]:

```
# plot the scale vs the shape parameter
fig, ax = plt.subplots(figsize=(7,4))
plt.plot(shape_set_t_h0, scale_fit_set_t_h0)
plt.xscale("log")
plt.yscale("symlog", linthreshy=10)
#plt.ylim([0,500])
plt.ylabel(r"scale parameter")
plt.xlabel("shape parameter")
plt.title(r"Both gamma fit $\theta$")
#fig.savefig("both_gamma_scale_theta_h0.pdf", bbox_inches="tight")
```

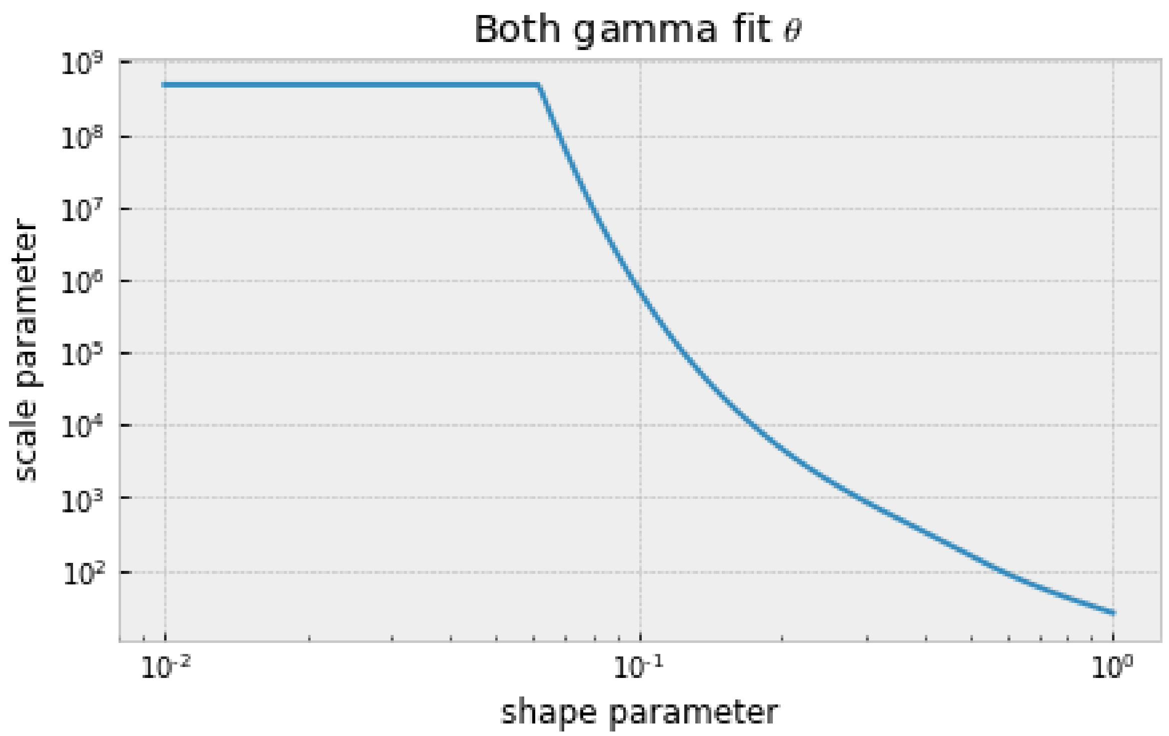


In [132]:

```
# visualising the observed SFS and expected SFS for the multinomial model
# the expected SFS can be extracted from the object calculated using the fi
tdadi 'spectra' function

fig, ax = plt.subplots(figsize=(7,7))

min_both_h0 = np.argmin(llhood_fit_set_h0)
fit_SP_both_h0 = spectra_SP_opt_h0.integrate([shape_set_h0[min_both_h0], sc
ale_fit_set_h0[min_both_h0]], gamma_dist, 1)
fit_PL_both_h0 = spectra_PL_opt_h0.integrate([shape_set_h0[min_both_h0], sc
ale_fit_set_h0[min_both_h0]], gamma_dist, 1)

min_PL_h0 = np.argmin(PL_llhood_fit_set_h0)
min_SP_h0 = np.argmin(SP_llhood_fit_set_h0)
fit_SP_only_h0 = spectra_SP_opt_h0.integrate([SP_shape_set_h0[min_SP_h0], S
P_scale_fit_set_h0[min_SP_h0]], gamma_dist, 1)
fit_PL_only_h0 = spectra_PL_opt_h0.integrate([PL_shape_set_h0[min_PL_h0], P
L_scale_fit_set_h0[min_PL_h0]], gamma_dist, 1)


plt.plot(fit_SP_both_h0.fold()/np.sum(fit_SP_both_h0.fold()),
         color="red", label="SP gamma both")
plt.plot(fit_PL_both_h0.fold()/np.sum(fit_PL_both_h0.fold()),
         color="blue", label="PL gamma both")
plt.plot(fit_SP_only_h0.fold()/np.sum(fit_SP_only_h0.fold()),
         "o", color="red", label="SP gamma only")
plt.plot(fit_PL_only_h0.fold()/np.sum(fit_PL_only_h0.fold()),
         "o", color="blue", label="PL gamma only")
```

```
plt.plot(data_SP_0fold/np.sum(data_SP_0fold), "--",
         color="red", label="SP (0-fold)")
plt.plot(data_PL_0fold/np.sum(data_PL_0fold), "--",
         color="blue", label="PL (0-fold)")

plt.yscale("log")
plt.ylabel("proportion of sites")
plt.xlabel("sample count (folded)")
plt.legend()

#fig.savefig("gamma_fits_h0.pdf", bbox_inches="tight")
```

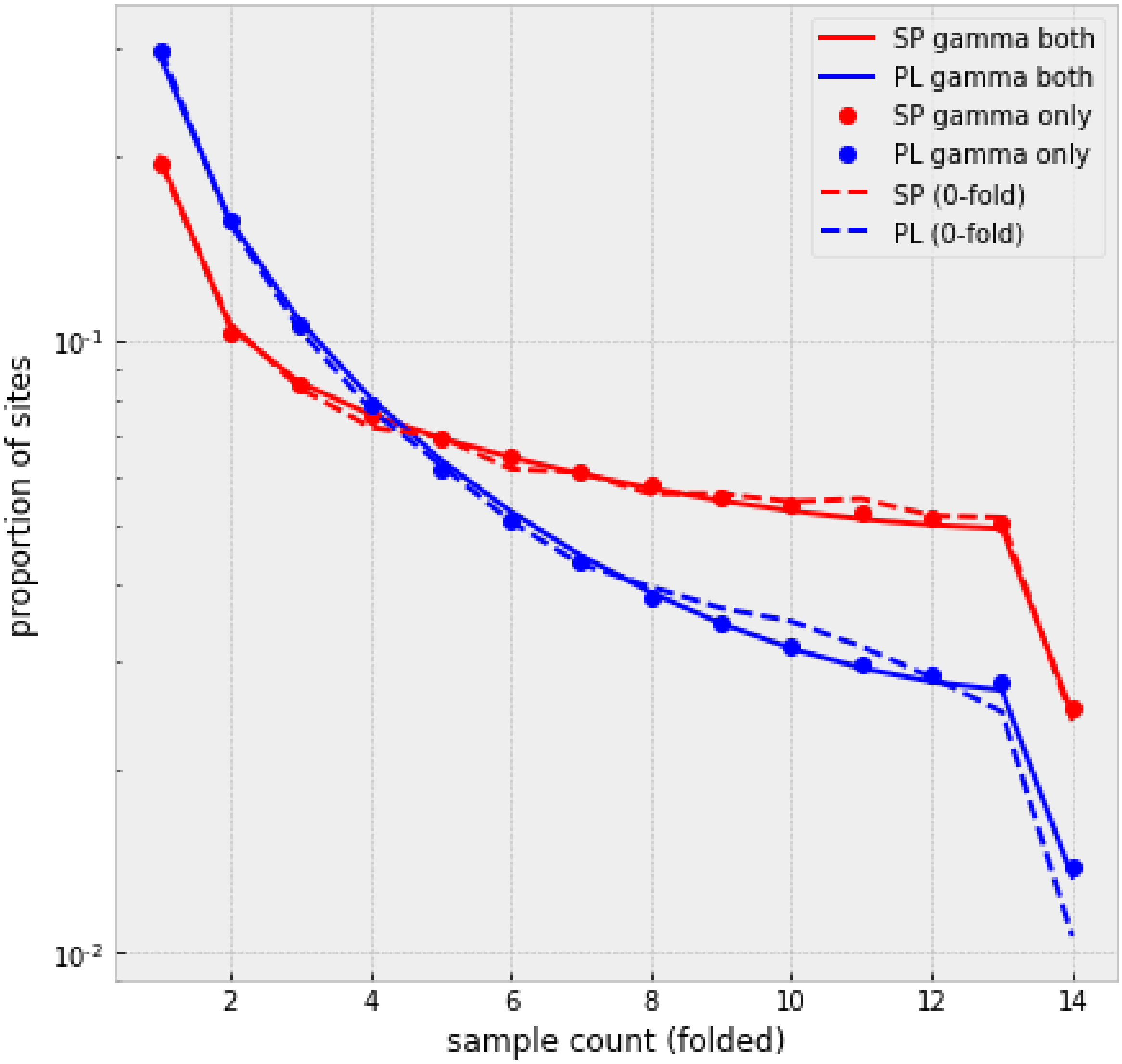


In [155]:

```
shape_set_h0[min_both_h0], scale_fit_set_h0[min_both_h0]
```

Out[155]:

```
(0.43910691980449945, 16.807932710318287)
```

In [119]:

```
# visualising the observed SFS and expected SFS for the Poisson model

fig, ax = plt.subplots(figsize=(8,8))
```

```
min_both_t_h0 = np.argmin(llhood_fit_set_t_h0)
fit_SP_both_t_h0 = spectra_SP_opt_h0.integrate([shape_set_t_h0[min_both_t_h
0], scale_fit_set_t_h0[min_both_t_h0]], gamma_dist, theta_SP_opt_ns)
fit_PL_both_t_h0 = spectra_PL_opt_h0.integrate([shape_set_t_h0[min_both_t_h
0], scale_fit_set_t_h0[min_both_t_h0]], gamma_dist, theta_PL_opt_ns)

min_PL_t_h0 = np.argmin(PL_llhood_fit_set_t_h0)
min_SP_t_h0 = np.argmin(SP_llhood_fit_set_t_h0)
fit_SP_only_t_h0 = spectra_SP_opt_h0.integrate([SP_shape_set_t_h0[min_SP_t_
h0], SP_scale_fit_set_t_h0[min_SP_t_h0]], gamma_dist, theta_SP_opt_ns)
fit_PL_only_t_h0 = spectra_PL_opt_h0.integrate([PL_shape_set_t_h0[min_PL_t_
h0], PL_scale_fit_set_t_h0[min_PL_t_h0]], gamma_dist, theta_PL_opt_ns)

plt.plot(fit_SP_both_t_h0.fold(), color="red", label="SP gamma both")
plt.plot(fit_PL_both_t_h0.fold(), color="blue", label="PL gamma both")
plt.plot(fit_SP_only_t_h0.fold(),  "o", color="red", label="SP gamma only")
plt.plot(fit_PL_only_t_h0.fold(), "o", color="blue", label="PL gamma only")
plt.plot(data_SP_0fold, "--", color="red", label="SP (0-fold)")
plt.plot(data_PL_0fold, "--", color="blue", label="PL (0-fold)")

plt.yscale("log")
plt.ylabel("number of sites")
plt.xlabel("sample count (folded)")
plt.legend()
#fig.savefig("gamma_fits_theta_h0.pdf", bbox_inches="tight")
```

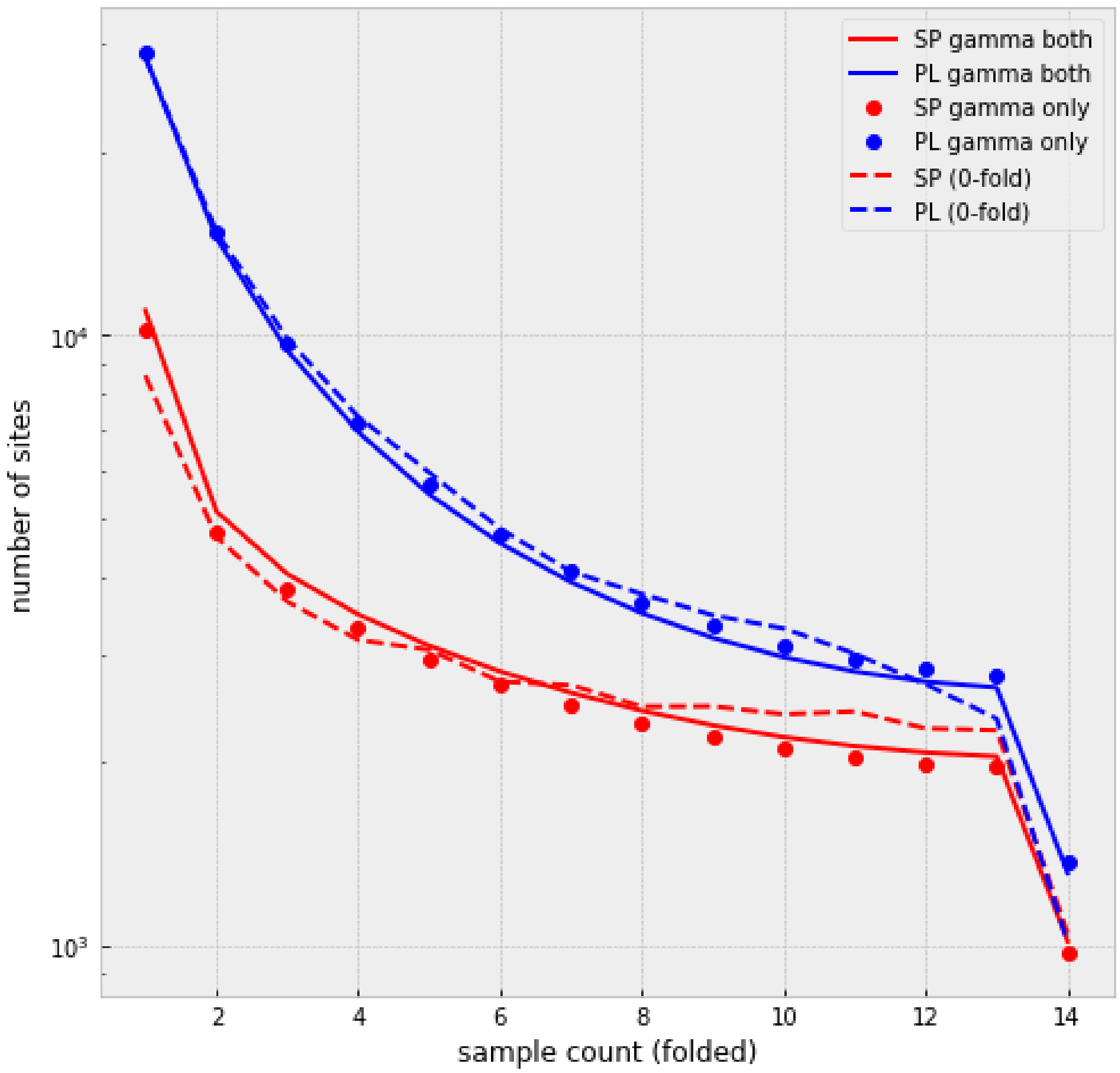


In [156]:

```
shape_set_t_h0[min_both_t_h0], scale_fit_set_t_h0[min_both_t_h0]
```

Out[156]:

```
(0.15443830272249795, 20396.03041368727)
```

In [131]:

```
fig, ax = plt.subplots(figsize=(7,7))

plt.plot(fit_SP_both_t_h0.fold()/np.sum(fit_SP_both_t_h0.fold()), color="re
d", label="SP gamma both")
plt.plot(fit_PL_both_t_h0.fold()/np.sum(fit_PL_both_t_h0.fold()), color="bl
ue", label="PL gamma both")
plt.plot(fit_SP_only_t_h0.fold()/np.sum(fit_SP_only_t_h0.fold()),  "o", col
or="red", label="SP gamma only")
plt.plot(fit_PL_only_t_h0.fold()/np.sum(fit_PL_only_t_h0.fold()), "o", colo
r="blue", label="PL gamma only")
plt.plot(data_SP_0fold/np.sum(data_SP_0fold), "--", color="red", label="SP
(0-fold)")
```

```
plt.plot(data_PL_0fold/np.sum(data_PL_0fold), "--", color="blue", label="PL
(0-fold)")

plt.yscale("log")
plt.ylabel("proportion of sites")
plt.xlabel("sample count (folded)")
plt.legend()
#fig.savefig("gamma_fits_theta_prop_h0.pdf", bbox_inches="tight")
```

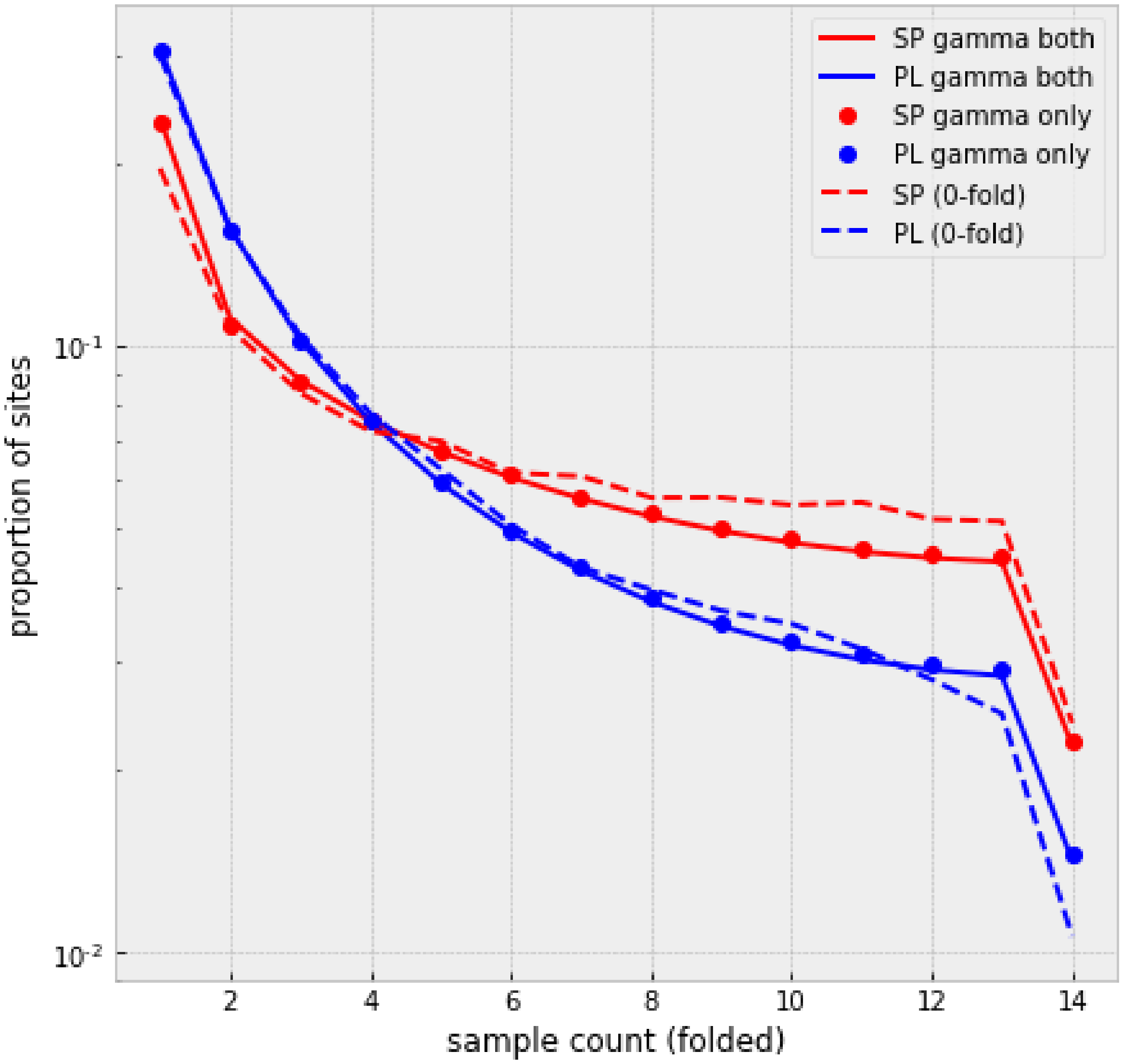


In [80]:

```
# visualising the cummulative distribution function (CDF) of the DFE
# the gamma distribution is calculated using the 'gamma.cdf' function and t
he previously calculated shape and scale parameters
# for SP two population sizes are shown, at the split with PL and the final
size

# calculating the CDF
from scipy.stats import gamma, lognorm
Ns_SP_set_h0 = -spectra_SP_opt_h0.gammas[::-1]
```

```
DFE_cdf_both_h0 = gamma.cdf(Ns_SP_set_h0, a=shape_set_h0[min_both_h0], scal
e=scale_fit_set_h0[min_both_h0])
DFE_cdf_both_t_h0 = gamma.cdf(Ns_SP_set_h0, a=shape_set_t_h0[min_both_t_h0]
, scale=scale_fit_set_t_h0[min_both_t_h0])
DFE_cdf_both_SP_t_h0 = gamma.cdf(Ns_SP_set_h0, a=SP_shape_set_t_h0[min_SP_t
_h0], scale=SP_scale_fit_set_t_h0[min_SP_t_h0])
DFE_cdf_both_PL_t_h0 = gamma.cdf(Ns_SP_set_h0, a=PL_shape_set_t_h0[min_PL_t
_h0], scale=PL_scale_fit_set_t_h0[min_PL_t_h0])

# plot CDF for the multinomial model
fig, ax = plt.subplots(figsize=(8,8))
ax.plot(Ns_SP_set_h0*PLnu_opt, DFE_cdf_both_h0, "-", color="blue", label="P
L final")
ax.plot(Ns_SP_set_h0*SPnuF2_opt, DFE_cdf_both_h0, "--", color="red", label=
"SP post-split")
ax.plot(Ns_SP_set_h0*SPnuF1_opt, DFE_cdf_both_h0, "-", color="red", label="
SP final")

ax.set(xscale="log", xlabel=r"$2N_e s$", ylabel="CDF", title="Multinom CDFs
")
ax.legend()
#fig.savefig("SP_PL_DFE_multinom_both_h0.pdf")
```

Out[80]:

```
<matplotlib.legend.Legend at 0x7f51bad54750>
```

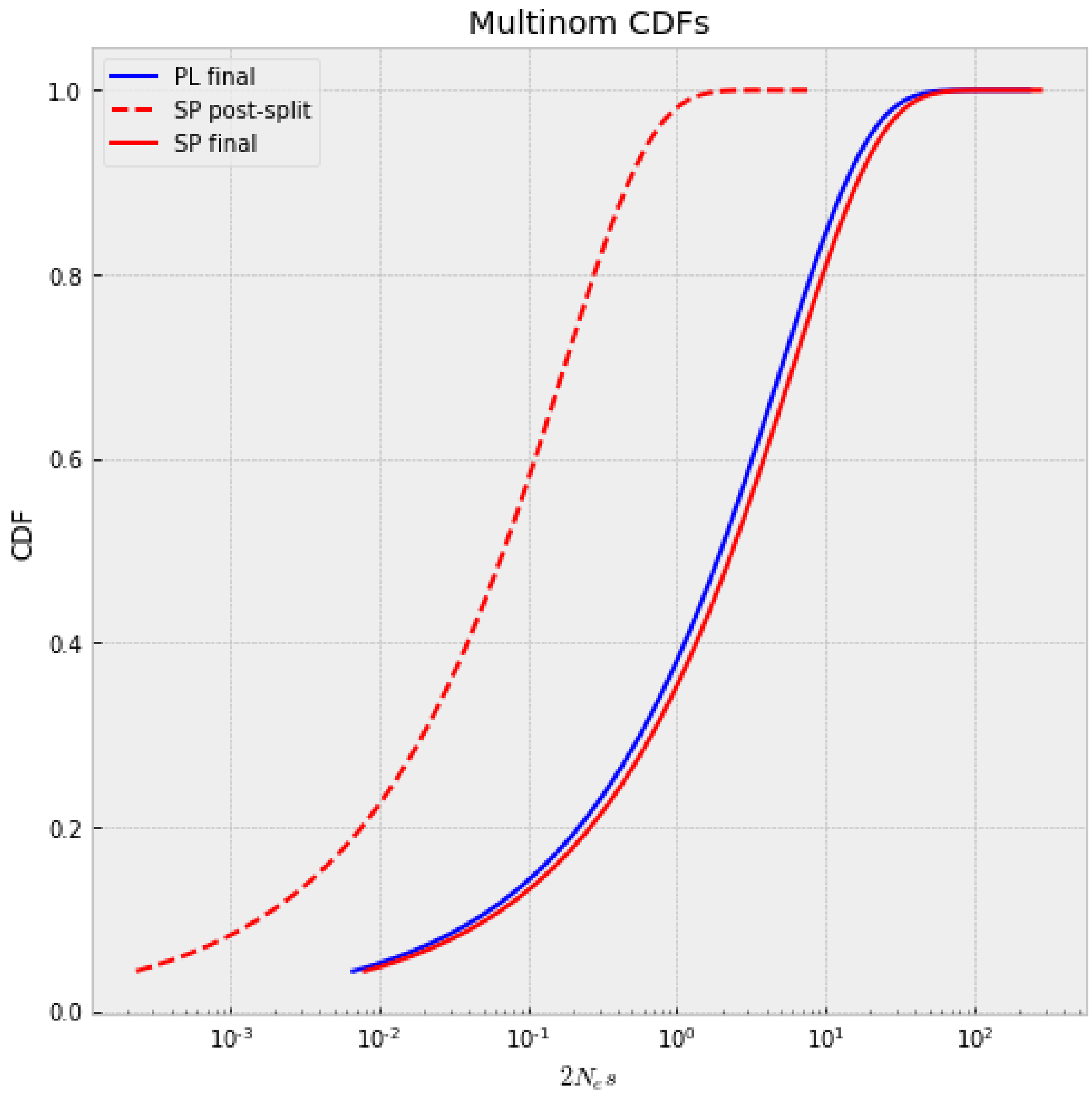


In [81]:

```
# plot CDF for the Poisson model (joint estimate)

fig, ax = plt.subplots(figsize=(8,8))
ax.plot(Ns_SP_set_h0*PLnu_opt, DFE_cdf_both_t_h0, "-", color="blue", label=
"PL")
ax.plot(Ns_SP_set_h0*SPnuF2_opt, DFE_cdf_both_t_h0, "--", color="red", labe
l="SP post-split")
ax.plot(Ns_SP_set_h0*SPnuF1_opt, DFE_cdf_both_t_h0, "-", color="red", label
="SP final")

ax.set(xscale="log", xlabel=r"$2N_e s$", ylabel="CDF", title=r"$\theta$ CDF
s both")
ax.legend()
#fig.savefig("SP_PL_DFE_theta_both_h0.pdf")
```

Out[81]:

```
<matplotlib.legend.Legend at 0x7f51bb073350>
```

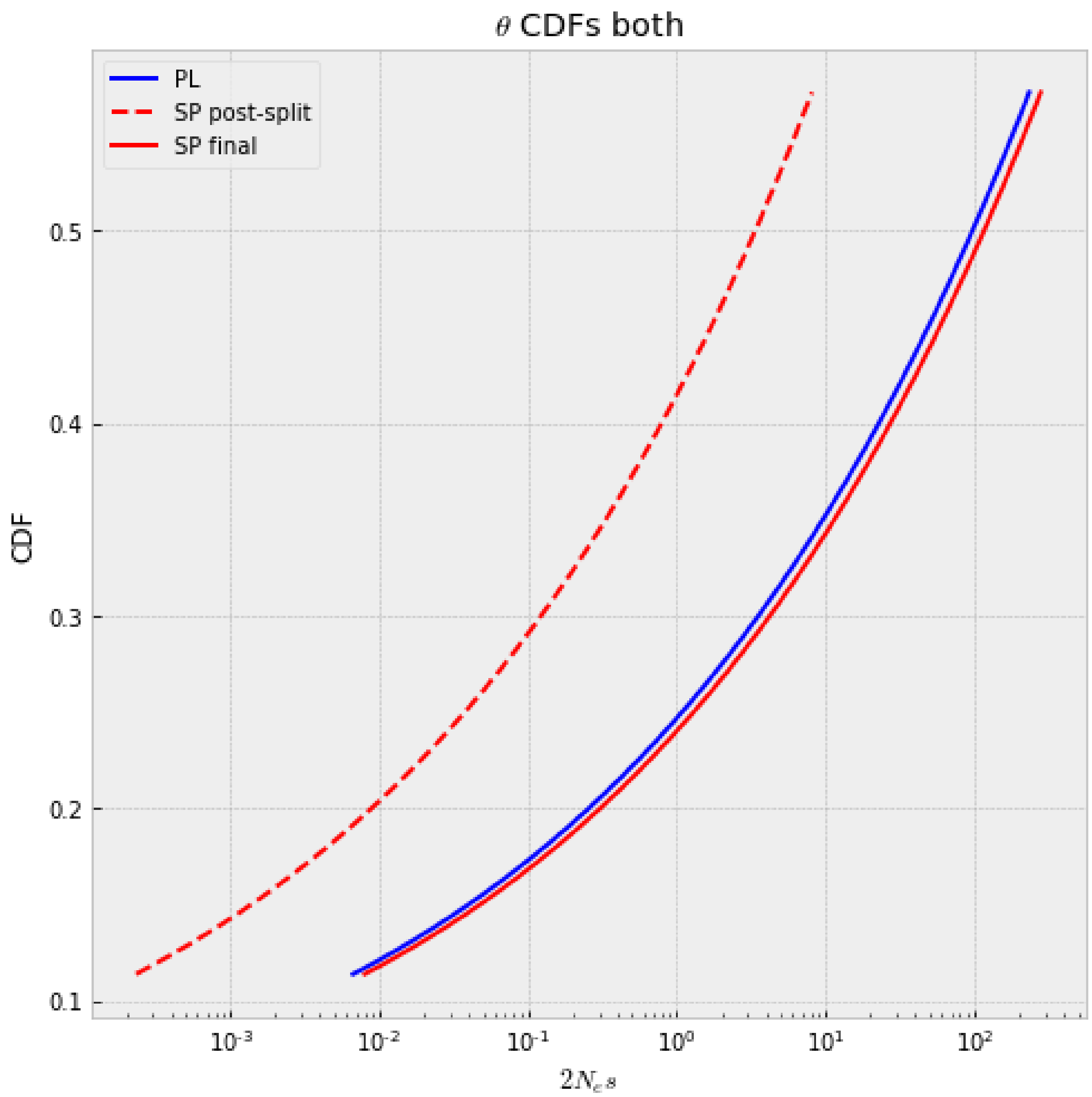


In [82]:

```
# calculate the proportion of segregating alleles for each bin of s for the
SFS
# for this we calculate the expected SFS for each value of s under the deog
raphic scenario
# and also the gamma distribution under the joint Poisson model for SP and
PL
# by applying Bayes' rule we can calculate the distribution of s for the va
riants in each count of the SFS

# calculate the expected SFS for a range of gammas
# this uses the demographic parameters and the selection function that was
previously defined
gamma_set_h0 = np.logspace(-1, 2.54, 100)
spectra_full_sel_h0 = []
for ii, gamma_tmp in enumerate(tqdm_notebook(gamma_set_h0)):
```

```
    spectra_full_sel_h0.append(three_epoch_growth_sel_both_new_h0(demo_para
ms_opt + [-gamma_tmp], 1, 100))
```

```
/home/ksteige/anaconda2/lib/python2.7/site-packages/ipykernel_launcher.py:1
0: TqdmDeprecationWarning: This function will be removed in tqdm==5.0.0
Please use `tqdm.notebook.tqdm` instead of `tqdm.tqdm_notebook`
  # Remove the CWD from sys.path while we load stuff.
```

In [83]:

```
# get the gamma distribution using the joint Poisson model
# using the shape and scale parameter with the best LL for both populations
# the gamma distribution is calculated by the 'gamma.cdf' function

F_gamma_both_h0 = gamma.cdf(gamma_set_h0, a=shape_set_t_h0[min_both_t_h0],
scale=scale_fit_set_t_h0[min_both_t_h0])
p_gamma_both_h0 = F_gamma_both_h0 - np.concatenate([[0], F_gamma_both_h0[0:
-1]])
p_gamma_both_h0[-1] += 1 - p_gamma_both_h0[-1]
```

In [84]:

```
# combine information of the expected SFS for each s and the gamma distribu
tion
# this allows us to get the proportion of s for each count of the SFS
PL_cDFE_set_h0 = []
SP_cDFE_set_h0 = []

# get proportion of s for each count of the SFS
from copy import deepcopy

for ii in range(14):
    cDFE_PL_tmp = np.zeros_like(gamma_set_h0)
    cDFE_SP_tmp = np.zeros_like(gamma_set_h0)
    for jj, gamma_tmp in enumerate(gamma_set_h0):
        cDFE_PL_tmp[jj] = p_gamma_both_h0[jj]*spectra_full_sel_h0[jj].margi
nalize([1]).fold()[ii+1]
        cDFE_SP_tmp[jj] = p_gamma_both_h0[jj]*spectra_full_sel_h0[jj].margi
nalize([0]).fold()[ii+1]
    PL_cDFE_set_h0.append(deepcopy(cDFE_PL_tmp/np.sum(cDFE_PL_tmp)))
    SP_cDFE_set_h0.append(deepcopy(cDFE_SP_tmp/np.sum(cDFE_SP_tmp)))

# get bins for s
gamma_bins = np.digitize(gamma_set_h0, np.array([-1, 0.1, 1, 10, 100, 1e6])
, right=True)
bins = np.arange(1, 6)
PL_bin_prob_set_h0 = []
SP_bin_prob_set_h0 = []
for ii in range(14):
    PL_bin_probs_h0 = [0]*5
    SP_bin_probs_h0 = [0]*5
    for jj, bin_ind in enumerate(bins):
```

```
        PL_bin_probs_h0[jj] = np.sum(PL_cDFE_set_h0[ii][gamma_bins == bin_i
nd])
        SP_bin_probs_h0[jj] = np.sum(SP_cDFE_set_h0[ii][gamma_bins == bin_i
nd])
    PL_bin_probs_h0 = np.array(PL_bin_probs_h0)
    SP_bin_probs_h0 = np.array(SP_bin_probs_h0)
    PL_bin_probs_h0[np.array(PL_bin_probs_h0) < 0] = 0
    SP_bin_probs_h0[np.array(SP_bin_probs_h0) < 0] = 0
    PL_bin_prob_set_h0.append(deepcopy(PL_bin_probs_h0)/np.sum(PL_bin_probs
_h0))
    SP_bin_prob_set_h0.append(deepcopy(SP_bin_probs_h0)/np.sum(SP_bin_probs
_h0))

PL_bin_prob_set_h0 = np.array(PL_bin_prob_set_h0)
SP_bin_prob_set_h0 = np.array(SP_bin_prob_set_h0)
```

In [85]:

```
from matplotlib.patches import Polygon
from matplotlib.collections import PatchCollection
from matplotlib import cm

cmap =["darkgrey","navy","cornflowerblue","mediumturquoise","darkorchid","i
ndigo"]

counts = np.arange(1, 15)

# plot binned s for each count of the SFS for PL
fig, axes = plt.subplots(1, 3, figsize=(27,7))

foo1 = Polygon(list(zip(counts, np.cumsum(PL_bin_prob_set_h0,1)[:,0])) + [(
14, 0.0), (1, 0.0)], color=cmap[5])
foo2 = Polygon(list(zip(counts, np.cumsum(PL_bin_prob_set_h0,1)[:,0])) +
               list(reversed(list(zip(counts, np.cumsum(PL_bin_prob_set_h0,
1)[:,1])))), color=cmap[4])
foo3 = Polygon(list(zip(counts, np.cumsum(PL_bin_prob_set_h0,1)[:,1])) +
               list(reversed(list(zip(counts, np.cumsum(PL_bin_prob_set_h0,
1)[:,2])))), color=cmap[3])
foo4 = Polygon(list(zip(counts, np.cumsum(PL_bin_prob_set_h0,1)[:,2])) +
               list(reversed(list(zip(counts, np.cumsum(PL_bin_prob_set_h0,
1)[:,3])))), color=cmap[2])
foo5 = Polygon(list(zip(counts, np.cumsum(PL_bin_prob_set_h0,1)[:,3])) + [(
1,1)], color=cmap[1])
p = PatchCollection([foo1, foo2, foo3, foo4, foo5], match_original=True, cm
ap=cm.jet)
axes[0].add_collection(p)
for ii, _ in enumerate(bins):
    axes[0].scatter(counts, np.cumsum(PL_bin_prob_set_h0,1)[:,ii], c=np.arr
ay([cmap[6-ii-1]]*14), edgecolor="black", s=60)
```

```
axes[0].set(ylim=[-0.05,1.05], ylabel="Proportion of segregating alleles",
xlabel="Sample count", title="PL")

# plot binned s for each count of the SFS for SP
foo1 = Polygon(list(zip(counts, np.cumsum(SP_bin_prob_set_h0,1)[:,0])) + [(
14, 0.), (1, 0.)], color=cmap[5])
foo2 = Polygon(list(zip(counts, np.cumsum(SP_bin_prob_set_h0,1)[:,0])) +
               list(reversed(list(zip(counts, np.cumsum(SP_bin_prob_set_h0,
1)[:,1])))), color=cmap[4])
foo3 = Polygon(list(zip(counts, np.cumsum(SP_bin_prob_set_h0,1)[:,1])) +
               list(reversed(list(zip(counts, np.cumsum(SP_bin_prob_set_h0,
1)[:,2])))), color=cmap[3])
foo4 = Polygon(list(zip(counts, np.cumsum(SP_bin_prob_set_h0,1)[:,2])) +
               list(reversed(list(zip(counts, np.cumsum(SP_bin_prob_set_h0,
1)[:,3])))), color=cmap[2])
foo5 = Polygon(list(zip(counts, np.cumsum(SP_bin_prob_set_h0,1)[:,3])) + [(
1,1)], color=cmap[1])
p = PatchCollection([foo1, foo2, foo3, foo4, foo5], match_original=True, cm
ap=cm.jet)
axes[1].add_collection(p)
for ii, _ in enumerate(bins):
    axes[1].scatter(counts, np.cumsum(SP_bin_prob_set_h0,1)[:,ii], c=np.arr
ay([cmap[6-ii-1]]*14), edgecolor="black", s=60)

axes[1].set(ylim=[-0.05,1.05], ylabel="Proportion of segregating alleles",
xlabel="Sample count", title="SP")

labels=[r"$\gamma \leq 0.1$", r"$0.1 > \gamma \leq 1$", r"$1 > \gamma \leq
10$", r"$10 > \gamma \leq 100$", r"$\gamma > 100$"]

# plot ratio PL / SP
for ii, _ in enumerate(bins):
    axes[2].plot(counts,(PL_bin_prob_set_h0[:,ii]/SP_bin_prob_set_h0[:,ii])
, c=cmap[6-ii-1])
    axes[2].scatter(counts, (PL_bin_prob_set_h0[:,ii]/SP_bin_prob_set_h0[:,
ii]), c=np.array([cmap[6-ii-1]]*14), edgecolor="black", s=60, label=labels[
ii])
axes[2].set(ylabel="PL/SP ratio", xlabel="Sample count")
axes[2].legend(loc=(.6, 0.1))

#fig.savefig("seg_effects_both_DFE_SP_PL_h0.pdf", bbox_inches="tight")
```

Out[85]:

```
<matplotlib.legend.Legend at 0x7f51b9f53690>
```

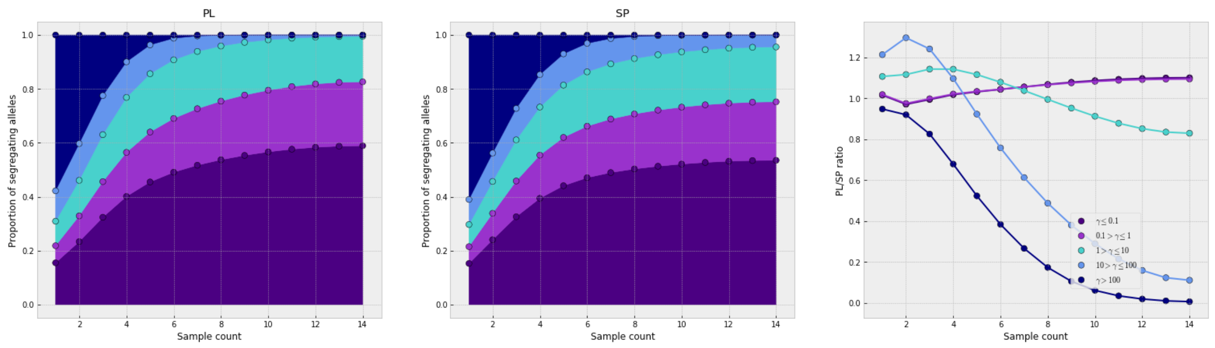


In [158]:

```
# calculate the burden difference between the populations
# using the Poisson model to calculate the expected SFS
# the difference in the burden between the populations is based on the cont
ribution of deleterious variants depending on their count in the population

min_both_m_h0 = np.argmin(llhood_fit_set_h0)
min_PL_m_h0 = np.argmin(PL_llhood_fit_set_h0)
min_SP_m_h0 = np.argmin(SP_llhood_fit_set_h0)

F_gamma_both_m_h0 = gamma.cdf(gamma_set, a=shape_set_t_h0[min_both_t_h0], s
cale=scale_fit_set_t_h0[min_both_t_h0])
p_gamma_both_m_h0 = F_gamma_both_m_h0 - np.concatenate([[0], F_gamma_both_m
_h0[0:-1]])
p_gamma_both_m_h0[-1] += 1 - p_gamma_both_m_h0[-1]

F_gamma_PL_m_h0 = gamma.cdf(gamma_set,  a=PL_shape_set_t_h0[min_PL_t_h0], s
cale=PL_scale_fit_set_t_h0[min_PL_t_h0])
p_gamma_PL_m_h0 = F_gamma_PL_m_h0 - np.concatenate([[0], F_gamma_PL_m_h0[0:
-1]])
p_gamma_PL_m_h0[-1] += 1 - p_gamma_PL_m_h0[-1]

F_gamma_SP_m_h0 = gamma.cdf(gamma_set,  a=SP_shape_set_t_h0[min_SP_t_h0], s
cale=SP_scale_fit_set_t_h0[min_SP_t_h0])
p_gamma_SP_m_h0 = F_gamma_SP_m_h0 - np.concatenate([[0], F_gamma_SP_m_h0[0:
-1]])
p_gamma_SP_m_h0[-1] += 1 - p_gamma_SP_m_h0[-1]
```

In [159]:

```
# get difference in burden from full DFE multinomial

from copy import deepcopy

full_PL_both_m_h0 = spectra_full_sel_h0[0].marginalize([1])
full_PL_both_m_h0.mask = False
full_PL_both_m_h0 *= 0
full_PL_PL_m_h0 = deepcopy(full_PL_both_m_h0)
full_PL_SP_m_h0 = deepcopy(full_PL_both_m_h0)

full_SP_both_m_h0 = spectra_full_sel_h0[0].marginalize([0])
```

```python
full_SP_both_m_h0.mask = False
full_SP_both_m_h0 *= 0
full_SP_PL_m_h0 = deepcopy(full_SP_both_m_h0)
full_SP_SP_m_h0 = deepcopy(full_SP_both_m_h0)

for ii, gamma_tmp in enumerate(gamma_set):
    tmp_PL = spectra_full_sel_h0[ii].marginalize([1])
    tmp_SP = spectra_full_sel_h0[ii].marginalize([0])
    tmp_PL.mask = False
    tmp_SP.mask = False
    full_PL_both_m_h0 += tmp_PL*p_gamma_both_m_h0[ii]
    full_SP_both_m_h0 += tmp_SP*p_gamma_both_m_h0[ii]
    full_PL_PL_m_h0 += tmp_PL*p_gamma_PL_m_h0[ii]
    full_SP_PL_m_h0 += tmp_SP*p_gamma_PL_m_h0[ii]
    full_PL_SP_m_h0 += tmp_PL*p_gamma_SP_m_h0[ii]
    full_SP_SP_m_h0 += tmp_SP*p_gamma_SP_m_h0[ii]
```

In [160]:

```python
fig, ax = plt.subplots(figsize=(8,8))cd ..
plt.plot(np.cumsum(full_PL_both_m_h0*np.arange(29)/28 - full_SP_both_m_h0*n
p.arange(29)/28)*theta_PL_opt_ns*2, label="Both DFE")
plt.plot(np.cumsum(full_PL_PL_m_h0*np.arange(29)/28 - full_SP_PL_m_h0*np.ar
ange(29)/28)*theta_PL_opt_ns*2, label="PL DFE")
plt.plot(np.cumsum(full_PL_SP_m_h0*np.arange(29)/28 - full_SP_SP_m_h0*np.ar
ange(29)/28)*theta_PL_opt_ns*2, label="SP DFE")
plt.yscale("symlog", linthreshy=100)
plt.ylabel("burden difference between PL and SP")
plt.xlabel("derived allele count")
plt.title("Full DFE")
plt.legend()

#fig.savefig("full_DFE_burden_h0_poisson.pdf", bbox_inches="tight")
```

```
WARNING:Spectrum_mod:Arithmetic between Spectra with different pop_ids. Res
ulting pop_id may not be correct.
WARNING:Spectrum_mod:Arithmetic between Spectra with different pop_ids. Res
ulting pop_id may not be correct.
WARNING:Spectrum_mod:Arithmetic between Spectra with different pop_ids. Res
ulting pop_id may not be correct.
```

Out[160]:

```
<matplotlib.legend.Legend at 0x7f51b85ffc90>
```

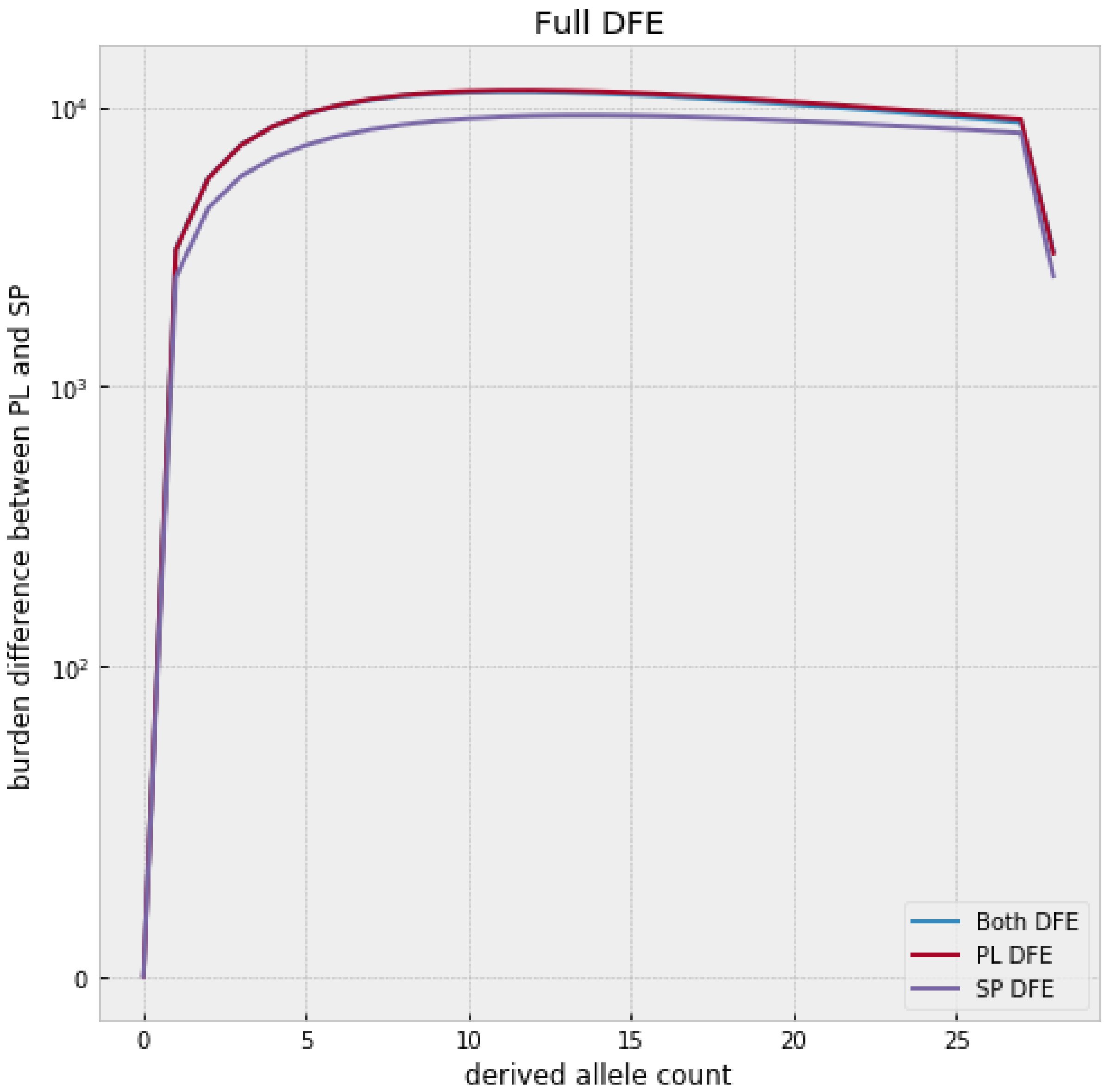

Full DFE
burden difference between PL and SP
derived allele count
10^4
10^3
10^2
0
0
5
10
15
20
25
Both DFE
PL DFE
SP DFE


In [ ]:
